# A universal method for gene expression engineering

**DOI:** 10.1101/644989

**Authors:** Rahmi Lale, Lisa Tietze, Jenny Nesje, Ingerid Onsager, Kerstin Engelhardt, Che Fai Alex Wong, Madina Akan, Niklas Hummel, Jörn Kalinowski, Christian Rückert, Martin Frank Hohmann-Marriott

**Author notes:** Correspondence and requests for materials should be addressed to Rahmi Lale and Martin F. Hohmann-Marriott.

## Abstract

The precise expression of genes is one of the foundations of biotechnology. Here we present GeneEE, a straightforward method for generating artificial gene expression systems. We demonstrate that GeneEE segments, containing a 200 nucleotide DNA with random nucleotide composition, can facilitate constitutive and inducible gene expression. To highlight the universal character of our method, we demonstrate GeneEE-mediated gene and protein expression in six bacterial species and Baker’s yeast.

## Introduction

Gene expression is modulated by the interaction of a complex transcription machinery with DNA^1^. In eubacteria, RNA polymerase initiates transcription after forming a complex with a sigma(σ)- factor, which recognises specific DNA sequence motifs. In addition, a multitude of additional proteins and RNA molecules participate in the modulation of gene expression. This complexity often thwarts the computational design of DNA sequences that result in desired levels of gene expression. In the absence of such computational tools, most of the gene expression studies rely on the use of native or modifications of existing promoters^2^.

Here, we present a Gene Expression Engineering (GeneEE) platform for generating artificial gene expression systems specifically designed for the gene of interest in the host of interest. The GeneEE platform leads to generation of both: artificial promoters, that recruit the native transcription machinery of the host microorganism; and 5′ untranslated regions (UTR), that prompt the translation of protein-coding DNA sequences. With this approach, GeneEE eliminates the reliance on previously characterised promoters and 5′ UTRs, and leads to generation of tailored gene expression systems.

The starting point for developing GeneEE was a back-of-the-envelope calculation for the probability of finding a σ-factor binding sequence within a segment of DNA with random nucleotide composition. The housekeeping *Escherichia coli* σ^70^ has two DNA binding domains that both interact with six nucleotides. If we assume that proper interaction of eight nucleotides with σ^70^ is enough to facilitate transcription initiation, then it would be expected that transcription initiation will occur with a probability of 1 in 65,536 DNA segments. For a DNA segment with 200 random nucleotides, 1 in 325 DNA segments would lead to transcription initiation—a surprisingly high proportion.

## Results & Discussion

To assess the validity of our calculations, we constructed two plasmid DNA libraries by cloning a stretch of DNA with random nucleotide composition (GeneEE segment) upstream of a β-lactamase coding sequence, that confers ampicillin resistance. To maintain the random DNA sequence composition, we developed a cloning strategy that avoids introduction of any scars upon cloning of the GeneEE segments [Supplementary Online Material (SOM) Figure 1, SOM Table 1 & 2, SOM sections 1.1-1.3]. For most chromosomal genes in *E. coli*, a Shine-Dalgarno (SD) sequence within the 5′ UTR of mRNA facilitates the recruitment of ribosomes^3^. Therefore we constructed plasmid DNA libraries with 211 nt long GeneEE segment containing a defined SD sequence (GGAG), GeneEE(+SD) along with a 200 nt long GeneEE segment consisting of entirely random nucleotides, GeneEE(–SD) [Figure 1a & b, SOM Figure 1]. After the transformation of both constructed plasmid DNAs into competent *E. coli* cells, aliquots were plated on agar plates supplemented with 50 μg/mL kanamycin, to determine the total number of constructs in the libraries; as well as on agar plates containing ampicillin (50, 500 and 1,000 μg/mL), to determine the frequency of clones that carry functional Artificial Promoter and 5′ UTR (ArtPromU) sequences [SOM section 1.4.8]. We found that 42% of the *E. coli* library, constructed with the GeneEE(–SD) segment, had *E. coli* clones that carry functional ArtPromU sequences [Figure 1c, SOM Table 9], whereas 33% of the *E. coli* library, constructed with the Gene(+SD) segment, had *E. coli* clones that carry functional ArtPromU sequences [Figure 1d]. The observation that a random nucleotide sequence with no defined SD sequence is more likely to initiate protein expression than SD-containing sequences indicates the prevalence of the use of alternative translation initiation mechanisms in many bacteria including *E. coli* ^4, 5^.

**Figure 1:**
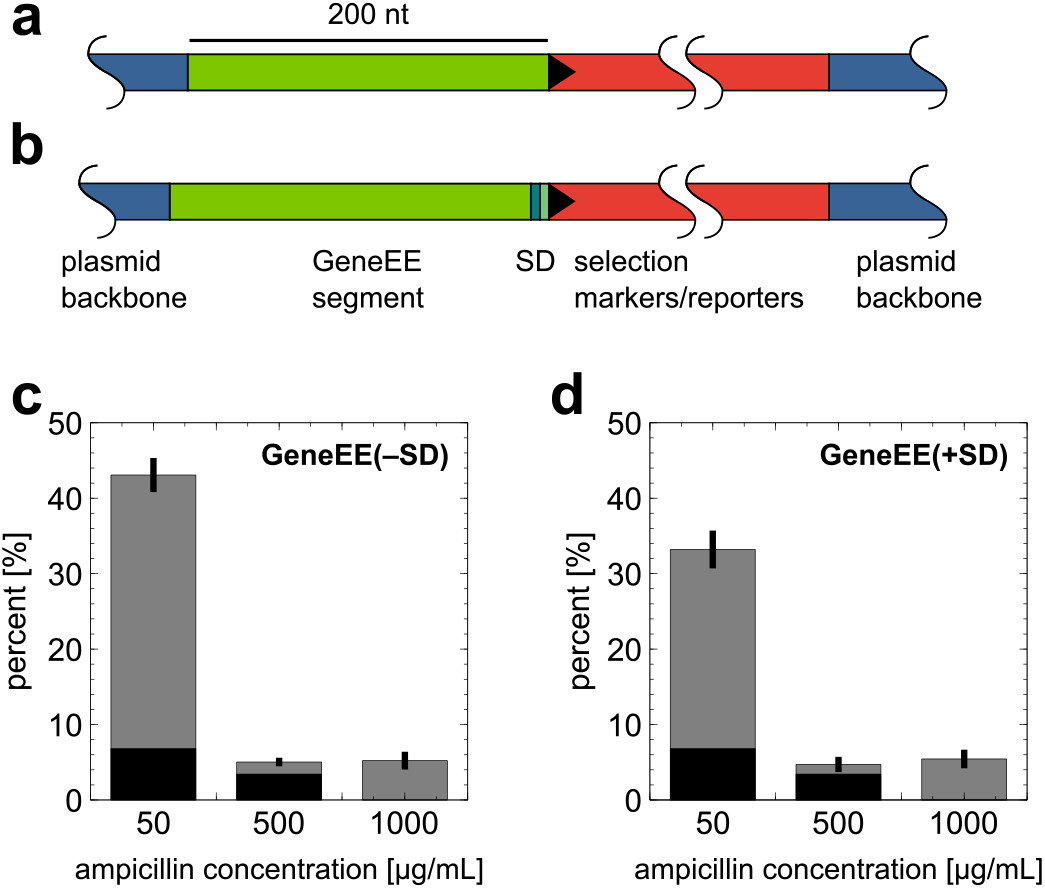
Schematic overview of the GeneEE segments, and the relative presence of clones with functional ArtPromU sequences in *E. coli*. Two plasmid DNA libraries were constructed: one with a GeneEE segment, GeneEE(–SD), containing a 200 nt DNA with random nucleotide composition (a); and second with a GeneEE segment with a defined SD sequence (GGAG), GeneEE(+SD) (b). The relative presence of clones with functional ArtPromU sequences was assessed by the cloning of the GeneEE segments upstream of the β-lactamase coding sequence (gene orientation indicated by an arrow head). The total number of clones in each library was determined by the count of kanamaycin resistant cells (dark columns). The relative presence of clones with different concentration of ampicillin resistance (grey columns) in each clone libraries carrying GeneEE(–SD) (c) and GeneEE(+SD) (d) are presented in percentage.

**Table 1:**
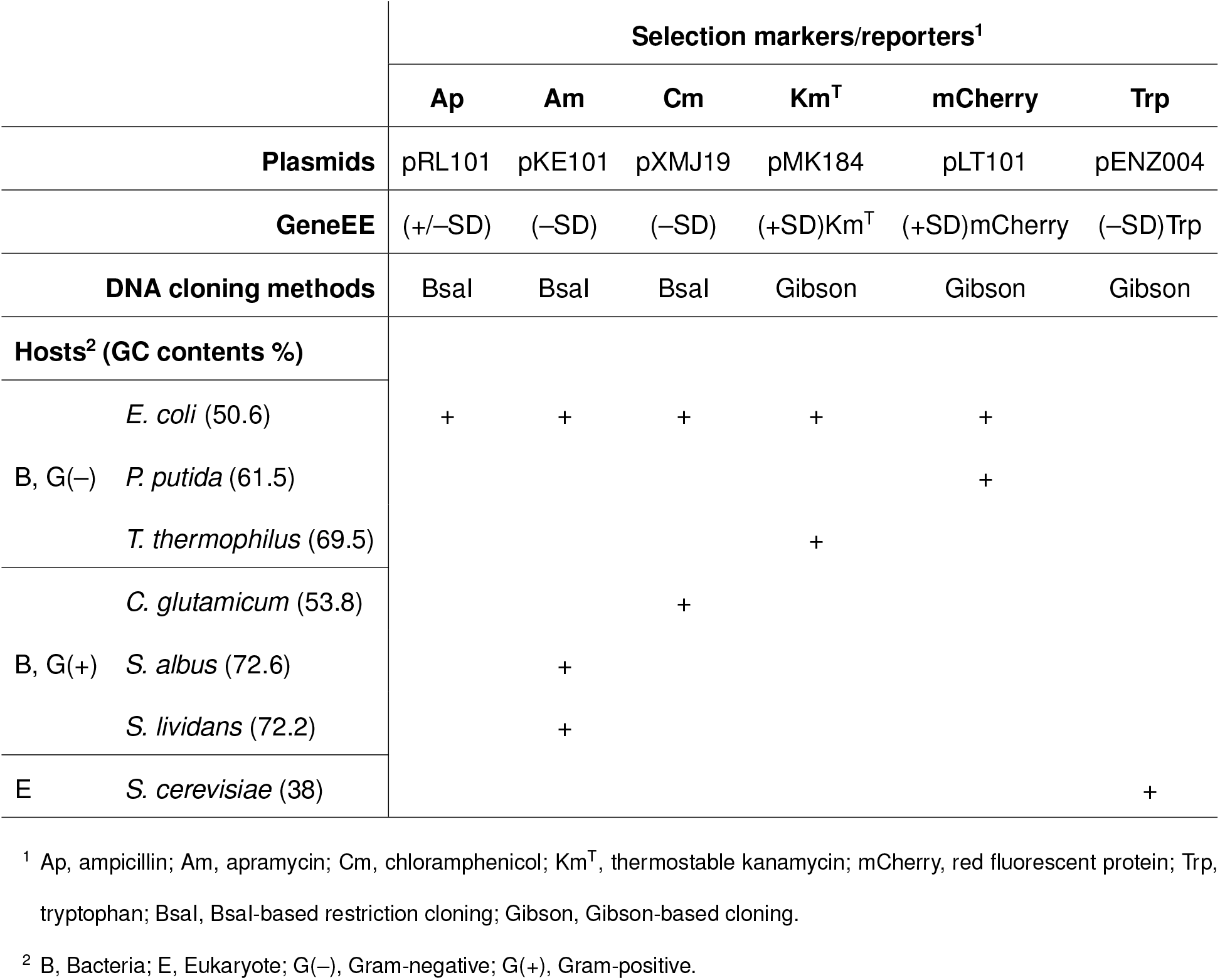
The hosts, GC contents, selection markers/reporters, plasmids, GeneEE segments (inserts) and the cloning methods used for the identification of ArtPromU sequences.

To gain further insight into the recruitment of the native transcription machinery by GeneEE segments, we constructed plasmid DNA libraries by placing the GeneEE(+SD) segment upstream of a DNA segment that encodes a red fluorescent protein (mCherry). In order to ensure that promoter sequences were followed with functional 5′ UTRs, in agreement with widely accepted notion of bacterial translation initiation, we used the GeneEE(+SD) segment as an insert for this part of the study. From a library consisting of ~5,000 clones, 192 *E. coli* clones expressing mCherry at various levels were picked from the agar plates, and the ArtPromU DNA sequences [SOM Table 6, SOM section 1.4.4, parse motifs.pl], and the transcription start sites (TSS)[SOM Table 7, SOM section 1.4.5, Supplementary spreadsheet 1, parse starts.pl] in each artificial promoter were experimentally determined by DNA and RNA sequencing, respectively. The mCherry protein production levels, resulting from each clone, were also measured [SOM Table 6]. The analysis of DNA sequencing data reveals the random nucleotide composition with high-sequence variation of the GeneEE(+SD) segment without any conserved positions or scars [Figure 2a]. A large variation in the numbers of *mCherry* transcripts [Figure 2b] and mCherry fluorescence intensities [Figure 2c] is observed. However, there is no correlation between the measured levels of mCherry fluorescence intensities and *mCherry* transcript abundance for the different clones. The majority of the ArtPromU sequences contain multiple TSS (up to 11) [Figure 2d], indicated by groups of transcripts with different mRNA lengths [SOM Table 7], and the TSS are located mostly closer to the translational start of the mCherry coding sequence [Figure 2e]. To identify potential promoter motifs within the artificial promoter sequences, the regions spanning from +1 to –50 were analysed and led to the identification of unique motifs [SOM section 1.4.6]. Similarly, motif analysis was also performed for the region downstream of the identified TSS, spanning from +1 to +25 (also known as the initially transcribed region within the 5′ UTR). This analysis also let to the identification of a unique motif [SOM section 1.4.7, SOM Table 8]. The identification of these previously unknown motifs demonstrates the power of the GeneEE platform as an exploration tool. In all artificial promoter sequences multiple σ-factor motifs could be detected, and the relative presence of conserved nucleotide positions positively correlate with the amount of produced *mCherry* transcripts [Figure 2f-k]. With these experimental findings we demonstrate that GeneEE segments can be used in generation of constitutive artificial gene expression systems in *E. coli*.

**Figure 2:**
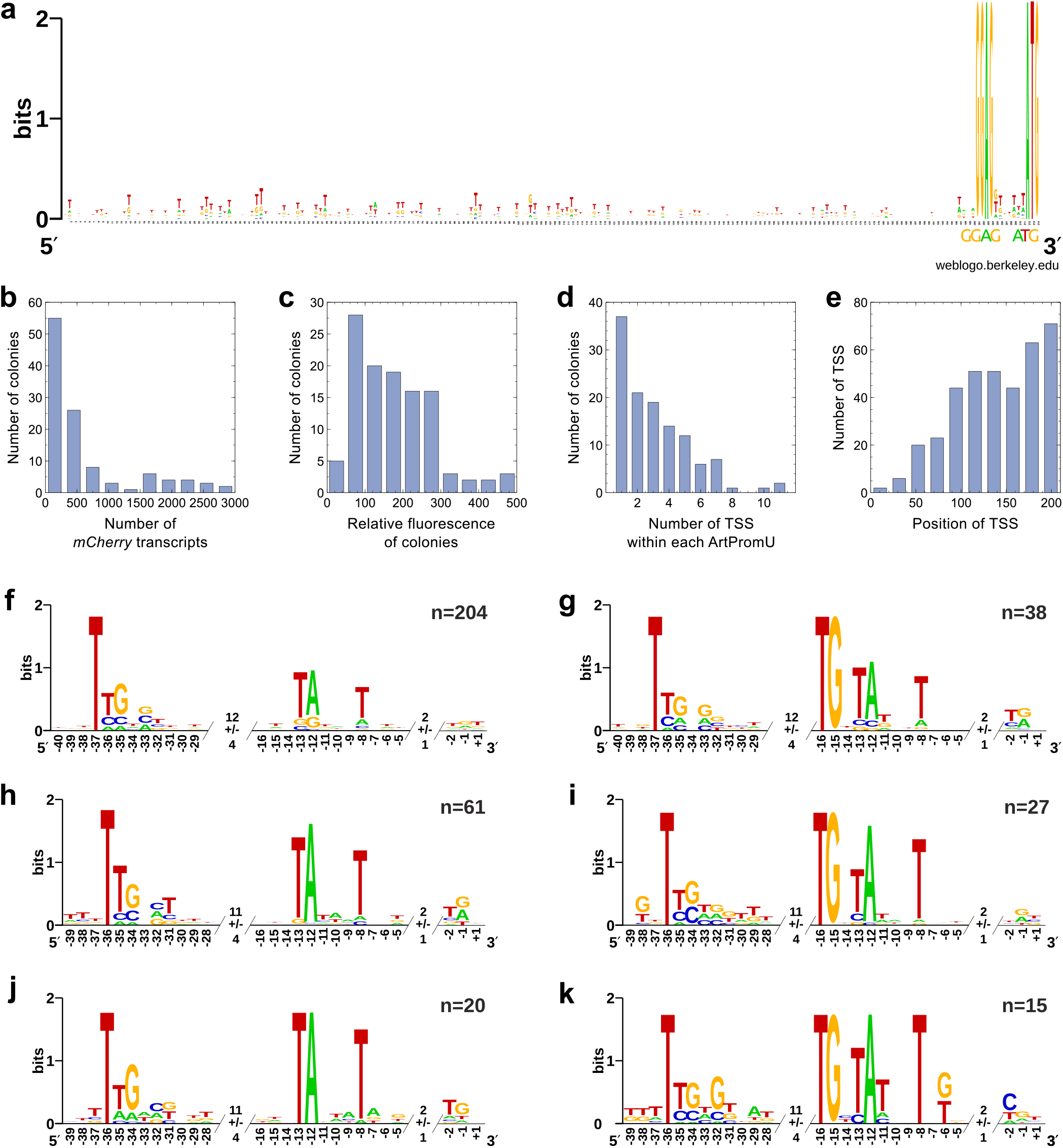
Multiple alignment of ArtPromU sequences, the analysis of transcription start sites (TSS) and promoter motif analysis. A plasmid DNA library was constructed using the *mCherry* gene, encoding for a red fluorescence protein, as a reporter for identification of clones that lead to GeneEE-mediated transcription and translation in *E. coli*. WebLogo^12^ image depicting the random nucleotide distribution of ArtPromU sequences expressing mCherry at varying levels (a). The defined SD sequence (GGAG), and the start codon of *mCherry* (ATG) are depicted in larger font size below the alignment. The total number of *mCherry* transcripts (b), relative fluorescence levels from each library colonies (c), the number (d) and position (e) of TSS identified within each ArtPromU sequences. Sequences grouped based on *mCherry* transcript abundance show distinct consensus sequences (motifs) for interaction with σ-factors (f-k). The sequences were realigned based on the identified motifs, cut down to the core motifs including 3 nt up- and downstream, and binned based on relative “transcription strength”, i.e. the number of mapped reads starting at each TSS as well as the presence or absence of the −10 extended motif TGn. For each bin, the median distance of the motifs was also calculated, the resulting numbers are given in the spacers. The numbers on the x-axis are based on these median distances. Motifs of “weak” promoters (10 to 99 mapped reads) without (f) and with (g) −10 extended motif. Motifs of “medium” promoters (100 to 999 mapped reads) without (h) and with (i) −10 extended motif. Motifs of “strong” promoters (1000 mapped reads and more) without (j) and with (k) −10 extended motif. The number of sequences used in each alignment are depicted with n above each panel.

Next, we investigated whether the GeneEE platform can also be used in recruiting other transcripton-relevant factors enabling inducible gene expression. For this study, we used the XylS/*Pm* system. In this system, XylS is a positive regulator protein that is a member of the AraC/XylS transcription factor family. Binding of the inducer (*m*-toluic acid), induces the dimerisation of XylS and the resulting XylS dimers bind to two DNA regions within the *Pm* promoter^6^, one of which overlaps with the −35 box σ-factor binding site by two base pairs. This interaction, in combination with σ^32^ or σ^38^, is thought to be the key in initiating transcription from *Pm*. In order to generate artificial *Pm* sequences, we constructed a plasmid that contained the *xylS* gene (coding sequence with its native constitutive promoter) and the β-lactamase coding sequence (excluding its native promoter). Using this plasmid, we constructed plasmid DNA libraries by cloning GeneEE(+SD) upstream of the β-lactamase coding sequence [SOM section 1.4.9]. Upon transforming plasmid DNAs to competent *E. coli* cells, a library consisting of ~30,000 clones was obtained. Using replica plating [SOM section 1.4.10], these transformants were either inoculated onto LA plates containing ampicillin and *m*-toluic acid, or only ampicillin. This screening led to the identification of 27 clones with inducible phenotype. DNA sequencing of the 27 clones revealed seven unique ArtPromU sequences conferring an inducible phenotype in *E. coli* [SOM Table 11]. In order to ensure that induction is XylS-dependent, we deleted the *xylS* gene from the seven plasmids and characterised the phenotype of the clones harbouring the plasmids without XylS. Among the seven clones, three clones had lost their inducible phenotype, indicating the reliance of induction on the presence of XylS. However, the remaining four clones still exhibited an inducible phenotype despite the absence of XylS, indicating the presence of an alternative mechanisms that responds to *m*-toluic acid in *E. coli*. The analysis of ArtPromU sequences that mediate XylS-dependent induction do not reveal the known binding sites for XylS. However, conserved thymidine nucleotides were present in all ArtPromU sequences that led to inducible phenotype [SOM Figure 4].

To demonstrate that the GeneEE platform can be applied to generate gene expression systems in different bacteria, we chose two Gram-negative *Pseudomonas putida*, *Thermus thermophilus*; and three Gram-positive bacteria, *Corynebacterium glutamicum, Streptomyces albus* and *S. lividans*, with a varying GC content [Table 1]. For these studies GeneEE segments, with and without defined SD sequences, were used to create five additional plasmid DNA libraries linked to a variety of selection markers/reporter genes [SOM sections 1.5-1.8]. Furthermore, we also applied GeneEE to the yeast model organism, Baker’s yeast *Saccharomyces cerevisiae* [SOM section 1.9]. Here, the GeneEE(–SD) was cloned upstream of a coding sequence conferring tryptophan-autotrophy into the *S. cerevisiae* strain with an impaired TRP1 gene. The application of GeneEE results in the generation of artificial gene expression systems in all seven tested microorganisms using six markers/reporters [Table 1]. The DNA sequences of all identified ArtPromUs are presented in the SOM Table 12 to 17.

## Conclusions

In this report, we demonstrate that a DNA segment with random nucleotide composition can be used to generate artificial gene expression systems in seven different microorganisms. This conclusion is supported by the detailed analysis of the artificial promoter sequences identified in *E. coli*, based on the experimental determination of TSS, demonstrating that RNA polymerase holoenzyme can recognise a wide variety of DNA sequences for transcription initiation. Furthermore, the GeneEE segments can also provide 5′ UTRs that lead to modulation of protein expression, following non-canonical translation initiation in *E. coli* ^4^. Alternative mechanisms to translation initiation are also known from viruses, such as the T7-type translation in bacteria^7^ and internal ribosome entry site-type translation in eukaryotes^8^. We see the potential that the large scale application of the GeneEE platform will yield hitherto unknown motifs and mechanisms for translation initiation in bacteria.

Gene and protein expression by DNA segments with random nucleotide composition is not only of interest for biotechnology, but also for the evolution by acquisition of new traits via horizontal gene transfer. The stable inheritance of the new trait to an offspring is likely to happen if the gene confers a selective advantage to the recipient organism. Our study indicates that the probability for initiation of transcription and translation is not seriously restricted when an organism receives a foreign gene. Apart form these evolutionary implications, our XylS/*Pm* study also demonstrates that inducible gene expression systems can be generated by recruiting and identifying native transcription regulation systems. The entire repertoire of native transcription machinery, including transcription regulators/factors and small regulatory RNAs, can be recruited by the GeneEE segments. In addition, the GeneEE-derived 5′ UTR can form riboswitches that modulate protein expression through triggers, such as pH and temperature shifts and interactions with metabolites^9^. The identification of hitherto unknown transcription regulators that respond to a specific stimulus, opens completely new possibilities for the development of biosensor applications.

Our study demonstrates the potential of the GeneEE platform, using easily available low-throughput methods for assessing selection markers and reporter proteins in combination with replica plating. The GeneEE platform, in combination with high-throughput approaches, such as fluorescence activated cell sorting^10^and microfluidics^11^ will constitute a fast and versatile method for gene expression and protein production engineering in a plethora of microorganisms in future biotechnology and synthetic biology applications.

## Supporting information

perl script: motifs

perl script: starts

## Additional information

### Acknowledgements

We thank Dr. José Berenguer (Universidad Autnoma de Madrid) for providing the pMK184 plasmid and the *T. thermophilus* strain; Sara Castaño Cerezo and Gilles Truan (Institut National des Sciences Appliquées de Toulouse, INSA Toulouse, Biosystems and Process Engineering Laboratory) for providing the pENZ004 plasmid and the *S. cerevisiae* strain; Niels-Ulrik Frigaard (University of Copenhagen, Department of Biology) for the pUC19-BBa-Km plasmid; Jochen Schmid for his valuable feed-back to the manuscript. This research was supported in part by the NTNU-Discovery program; NTNU-Biotechnology, Enabling Technologies program; EU-H2020 MetaFluidics project with grant agreement number 685474; The Research Council of Norway grant number 244278.

### Competing Interests

R.L. and M.F.H-M are co-inventors on a pending patent (PCT/EP2018/058227) which has been submitted by the Norwegian University of Science and Technology, describing the developed platform in this study.

### Author Contributions

R.L. and M.F.H-M conceived and designed the study. L.T. involved the *C. glutamicum* and performed the yeast experiments; J.N. performed the inducible phenotype experiments in *E. coli*; I.O. involved in creation of libraries in *E. coli* and *C. glutamicum* experiments; K.E. performed the *Streptomyces* experiments; C.F.A.W performed the *T. thermophilus* experiments; M.A and N.H. involved in the establishment of the method in *E. coli*; J.K. and C.R. performed the DNA sequencing and TSS determination experiments; and R.L. and M.F.H-M wrote the manuscript with feedback from all co-authors.

## Notes

#### Summary of Updates

The acknowledgments section has been amended: added: "Jochen Schmid for his valuable feedback to the manuscript" corrected spelling: "Gilles Truan"

