## Supplementary Online Material for "A universal method for gene expression engineering"

Lale et al., 2019

**References** **56**

### 1 Materials & Methods

#### 1.1 The composition of the single-stranded random nucleotide oligo

For the construction of GeneEE DNA libraries two different versions of synthetic single-stranded random nucleotide oligos (ss-RaNuqO) were used, each purchased from IDT (Belgium) as a four nmole ultramer: (A) containing the random stretch of 200 nt, $N_{200}$, named ss-RaNuqO(–SD) (Figure 1A); (B) containing the defined GGAG sequence in between the two stretches of random nucleotides, $N_{200}$ and $N_7$, named ss-RaNuqO(+SD) (Figure 1B). In the last mentioned version, the defined GGAG sequence serves as the minimal bacterial consensus Shine-Dalgarno (SD) sequence, and the ±SD naming indicates whether the oligo harbours the SD sequence (+SD) or not (–SD).

The ss-RaNuqOs have adapters on both ends. The Adapter1 contains the BioBrick Prefix [1], and the Adapter2 contains the BioBrick Suffix [1] sequences (Figure 1). Each of the adapters also contain a type IIS restriction enzyme recognition sequence for BsaI (5′-GGTCTC(N1)/(N5)-3′), with overhang sequences ACGG in Adapter1 and NTAC in Adapter2. These adapters also serve two additional functions: firstly, they are used for the functional immobilisation of the oligo during the chemical synthesis; secondly, they are used to generate the complementary DNA strand by PCR leading to double-stranded random nucelotide DNA (GeneEE segments). The BsaI overhang sequence in the Adapter2 has been designed to contain an N nucleotide in the 3′ overhang sequence, NTAC, to ensure that all possible four nucleotides will be presented in this position, hence leaving no scar upon cloning.

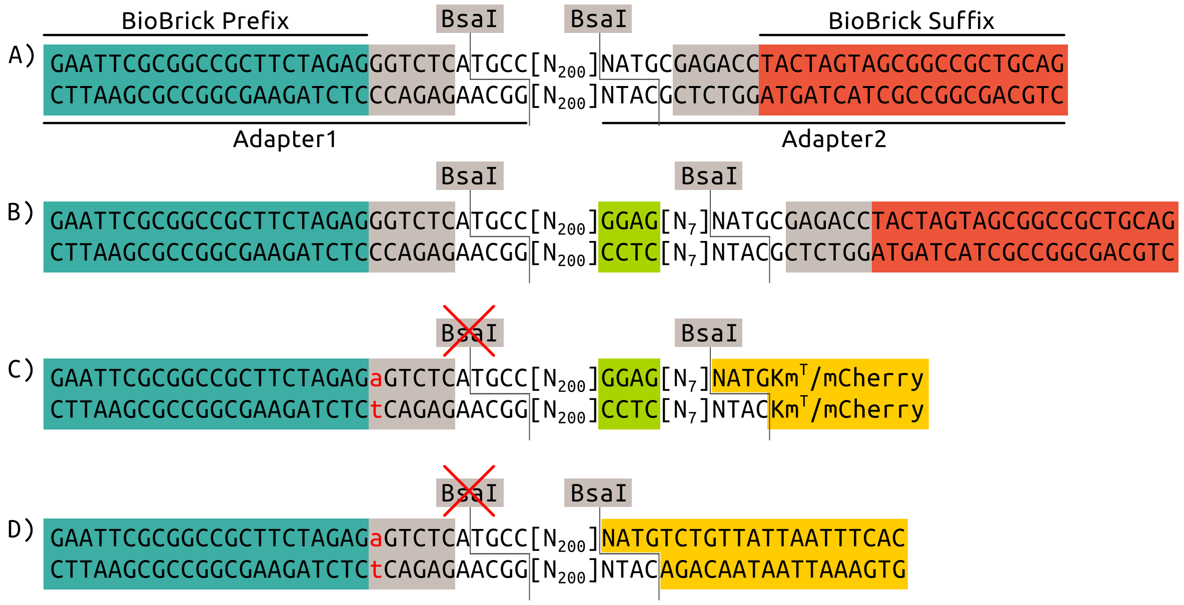

Figure 1: The nucleotide sequence composition of four different versions of GeneEE: A and B used for BsaI-based restriction cloning; C and D used for Gibson cloning. A) GeneEE(–SD), ss-RaNuqO in the upper strand and the complementary lower strand generated by PCR with primers targeting BioBrick Prefix and Suffix, containing the random stretch of 200 nt, $N_{200}$. The Adapter1 consists of the BioBrick Prefix (coloured in dark green) and BsaI recognition sequence (coloured in grey) and cut sites (marked with single lines); the Adapter2 is consisting of BsaI recognition sequence and cut site and BioBrick Suffix sequence (coloured in red). B) GeneEE(+SD), containing the defined GGAG sequence in between the two stretches of random nucleotides, $N_{200}$ and $N_7$. The Shine-Dalgarno sequence has been highlighted with light green colour. C) GeneEE(+SD)(+linkers for $Km^T$ or mCherry) that has been adapted for Gibson cloning. The single nucleotide marked with red colour is the single nucleotide mutation (G to a) introduced by PCR eliminating the BsaI recognition sequence. The dark yellow coloured box indicates the gene specific linkers for $Km^T$ and mCherry (Table 2). D) GeneEE(–SD)Trp, the GeneEE(–SD) segment that carries the linker sequence for the *tryptophan* gene.

#### 1.2 Generation of GeneEE segment for BsaI-based restriction cloning

The double-stranded GeneEE segment was generated by using either of the ss-RaNuqO(±SD) as the DNA template, and the BBa-Prefix-F and BBa-Suffix-R as primers (Table 1) in 10 cycles of PCR leading to GeneEE(–SD)BsaI (Figure 1A) and GeneEE(+SD)BsaI (Figure 1B). The low number of cycles was chosen to prevent a possible bias during the PCR amplification that can lead to reduced sequence diversity.

Table 1: The primers used in this study.

| Name | DNA Sequence (5′→3′) |
|---|---|
| BBa-Prefix-F | GAATTCGCGGCCGCTTCTAGAG |
| BBa-Suffix-R | CTGCAGCGGCCGCTACTAGTA |
| BBa-Prefix-delBsaI-F | GAATTCGCGGCCGCTTCTAGAGAGTCTCATGCC |
| delBsaI-F | AATGATACCGCGGGACCCACGCTCACCGGC |
| delBsaI-R | GGGTCCCGCGGTATCATTGCAGCACTGGGG |
| BsaI-Bla-F | GGCATGAGACCGTGCCACCTGACGTCTAAGAAA |
| BsaI-Bla-R | CTCGGTCTCCNATGAGTATTCAACATTTCCGTGTC |
| pRL101-Bla-Seq | CTTTCACCAGCGTTTCTGGGTG |
| mCherry-NdeI-F | GCTGCATATGGTTTCTAAAGGTGAAGAAG |
| mCherry-NdeI-R | GCTCGGATCCTTATCATTTATACAGTTCGTCCATAC |
| G-mCherry-F | ATGGTTTCTAAAGGTGAAGAAGACA |
| G-mCherry-R | CTCTAGAAGCGGCCGCGAATTCGCTGACGCCGTTGGATACAC |
| pLT101-mCherry-Seq | GATGTCAGCCGGGTGTTTAAC |
| $Km^T$-F | ATGAATGGACCAATAATAATGAC |
| $Km^T$-R | CTCTAGAAGCGGCCGCGAATTCGAGGCTAAGCCTTCCCTCAG |
| pMK184-$Km^T$-Seq | ATGGTAGAAATGTTGTCGGTCC |
| BsaI-Chl-F | CTCGGTCTCCNATGGAGAAAAAAATCACTGGATATACCAC |
| BsaI-Chl-R | CTGGCATGAGACCGCCCCGAAGAACGTTTTCCAAT |
| pXMJ19-Chl-Seq | TGAGCAACTGACTGAAATGCCT |
| Km-F | ACTGGTCCACCTACAACAAAGCTCTCATCAACCGTGGCTCTGATCATCTCGCGCAG CTCG |
| Km-R | GTGCAATGTAACATCAGAGATTTTGAGACACAACGTGGCTTGCCCTCATCTGTTAC GCCG |
| BsaI-Am5-F | AGTTCGGTCTCCNATGTCATCAGCGGTGGAGTGC |
| BsaI-oriV3-R | GGCATGAGACCTCGTTCCACTGAGCGTCAGACC |
| 5511-F | GCAGAGCGAGGTATGTAGGC |
| 234-R | TCGTCGTCCAGACCTGACCA |
| 5654-F-Seq | AAACAAACCACCGCTGGTAG |
| Am-5c-Seq | ACGCTACGGAAGGAGCTGTG |
| Trp-F | ATGTCTGTTATTAATTTCACAG |
| Trp-R | CTCTAGAAGCGGCCGCGAATTCTACCCATCTATGCTGAAG |
| Hom-F | CGTCTATGAGGAGACTGTTAG |
| Hom-R | CTCGCCAAGGCATTACC |
| pENZ004-Trp-Seq | CACTACTCTCTTCCTAGTCG |
| XylS-EcoRI-F | TAGAATTCCAGACCGCTTCTGCGTTCTG |
| XylS-XbaI-F | TATCTAGAAGGAACCCGACCTGCATTGG |
| XylS-delBsaI-F | CGCCTGGATTAGAAACCTGTTATCATCTGC |
| XylS-delBsaI-R | AACAGGTTTCTAATCCAGGCGAGATTACCC |

Table 1: continued from the previous page.

| | |
|---|---|
| Illumina-fwd | CACGACGCTCTTCCGATCTnnnnGGTGTATCCAACGGCGTCAG |
| Illumina-rev | CAGACGTGTGCTCTTCCGATCATGTTGTCTTCTTCACCTTTAGAAACC |
| Illumina-TSS | AGACGTGTGCTCTTCCGATCTGACCGTTCACAGAACCTTCC |

#### 1.3 Generation of GeneEE segment for Gibson cloning

The GenEE(±SD) segments (Figure 1C and D) were generated by using either of the ss-RaNuqO(±SD) as the DNA template and the BBa-Prefix-delBsaI-F and BBa-Suffix-R as primers (Table 1) in 10 cycles of PCR. The BBa-Prefix-delBsaI-F carries a single nucleotide mismatch eliminating the BsaI recognition sequence upon PCR amplification (Figure 1C and D). The resulting PCR product was digested by BsaI removing the Adapter2 and leaving the 3′-NTAC overhang. In this study, three linkers were ligated to generate three versions of the GeneEE(±SD) segments each carrying a specific linker for the *mCherry*, $Km^T$ or *trp* genes.

**The ligation of gene specific linkers.** Three complementary oligo pairs for each gene were ordered from Sigma-Aldrich (Germany) as unphosphorylated oligos. The oligos were designed so that upon annealing they would carry the required 5′-NATG overhang (Table 2). First, each oligo was phosphorylated by T4 Polynucleotide Kinase (NEB) and afterwards annealed to each other. The resulting double-stranded linkers with sticky-ends were ligated to the BsaI digested GeneEE±SD segments by T4 Quick DNA Ligase (NEB) leading to: GeneEE(+SD)mCherry; GeneEE(+SD)$Km^T$ (Figure 1C), and GeneEE(–SD)Trp (Figure 1D). And finally, all three GeneEE segments were PCR amplified using primer pair BBa-Prefix-F and the corresponding UpS oligo of the gene of interest.

Table 2: The gene specific linker sequences used in this study.

| **Name**[1] | **Sequence (5′→3′)** |
|---|---|
| $Km^T$-UpS | NATGAATGGACCAATAATAATGACTAG |
| $Km^T$-LoS | CTAGTCATTATTATTGGTCCATT |
| mCherry-UpS | NATGGTTTCTAAAGGTGAAGAAGACA |
| mCherry-LoS | TGTCTTCTTCACCTTTAGAAAC |
| Trp-UpS | NATGGTTTCTAAAGGTGAAGAAGACA |
| Trp-LoS | TGTCTTCTTCACCTTTAGAAAC |

[1] $Km^T$, thermostable kanamycin used in *T. thermophilus*; mCherry, used in *E. coli* and *P. putida*; Trp, *tryptophan* gene used in *S. cerevisiae*; UpS, upper DNA strand; LoS, lower DNA strand.

In this study, seven microorganisms were used: G(–) bacteria, *Escherichia coli, Pseudomonas putida, Thermus thermophilus*; G(+) bacteria, *Corynebacterium glutamicum, Streptomyces albus, Streptomyces lividans* and a yeast, *Saccharomyces cerevisiae.* For the details of the strains used, please refer to Table 3.

#### 1.4 *Escherichia coli*

##### 1.4.1 Growth conditions.

*E. coli* cells were grown in Lysogeny Broth (LB, 10 g/L tryptone, 5 g/L yeast extract, 5 g/L NaCl) or Lysogeny Agar (LA, LB + 15 g/L agar) at 37 °C, supplemented with 50 µg/mL kanamycin or varying levels of ampicillin as specified in the text.

Table 3: Microorganisms used in this study.

| Microorganisms | Description | Reference |
|---|---|---|
| **G(−) bacteria** | | |
| *E. coli* DH5α | Standard cloning strain | NEB |
| *E. coli* S17-1 | A conjugative strain used in transfer of DNA to *Streptomyces* spp. | 2 |
| *E. coli* C2984 | High efficiency turbo competent cells used for library construction. | NEB |
| *P. putida* KT2440 | A metabolically versatile, and well characterised bacterium with high tolerance for toxic chemicals. | 3 |
| *T. thermophilus* HB27Δago | A knockout strain of *T. thermophilus* HB27 with high transformation efficiency. | 4 |
| **G(+) bacteria** | | |
| *C. glutamicum* ATCC 13032 | A commonly used industrial strain in the production of amino acids. | 5 |
| *S. albus* J1074 | Derivative of *S. albus* DSMZ 40313, isoleucine and valine auxotrophic, deficient of SalGI-based restriction modification system. Widely used for heterologous protein/antibiotic production. | 6 |
| *S. lividans* TK24 | Plasmid-free derivative strain of *S. lividans* 66. Model *Streptomyces* strain routinely used for heterologous protein production. | 7 |
| **Eukaryot** | | |
| *S. cerevesia* CEN.PK2-1C | A tryptophan auxotroph *S. cerevisiae* CEN.PK strain commonly used as a yeast model organism. | 8 |

##### 1.4.2 The construction of GeneEE plasmid DNA libraries in *E. coli*.

For the functional screening of GeneEE segments in *E. coli*, two different genes were used: an antibiotic selection marker, *β-lactamase*, conferring ampicillin resistance; and a fluorescent reporter, *mCherry*, a red fluorescent protein variant.

For the construction of GeneEE plasmid DNA libraries with the *β-lactamase* gene pUC19-BBa-Km was used (Table 4). This plasmid carries a functional kanamycin resistance marker (native kanamycin promoter and coding sequence) cloned into the multiple cloning site of pUC19. The β-lactamase coding sequence carries a BsaI recognition sequence and in order to eliminate this site the entire plasmid was amplified by the primer pair delBsaI-F and delBsaI-R (Table 1), by following the Overlap Extension PCR cloning method[9], and the resulting linear PCR product was transformed to *E. coli* via chemical transformation and grown on LA plates supplemented with 50 µg/mL ampicillin. The primers carry a single nucleotide point mutation hence eliminating the BsaI recognition sequence. The resulting plasmid, pRL101, was then amplified by the primer pair BsaI-Bla-F and BsaI-Bla-R amplifying the entire plasmid excluding the native *β-lactamase* promoter. Both the PCR product and the GeneEE(+SD)BsaI was digested with BsaI and used in cloning as a vector and insert, respectively. The ligation mix was made using 1:5 molar vector:insert ratio in a total volume of 20 µL using the T4 Quick Ligase (NEB). The mixture was then transferred into the cloning host *E. coli* via chemical transformation.

For the construction of GeneEE plasmid DNA libraries with the *mCherry* gene a mini-RK2-based broad host range plasmid pHH100 (Table 4) was used. An *E. coli* codon-optimised variant of the *mCherry* gene (a gift from

Table 4: Plasmids used in this study.

| Plasmids | Description | Reference |
|---|---|---|
| pMK184 | An *E. coli*-*T. thermophilus* shuttle vector harbouring a thermostable kanamycin resistance gene; $Km^R$. | [10] |
| pKC1218 | Conjugative *E. coli*-*Streptomyces* shuttle vector. SCP2* *Streptomyces* replicon; pMB1 *E. coli* replicon; aac(3)IV gene conferring apramycin resistance; $Am^R$. | [11] |
| pKE101 | pKC1218-derivative with inserted aph(3′) cassette between oriT and SCP2* conferring kanamycin resistance; $Km^R$. | This study |
| pHH100 | Mini-RK2-based broad-host range low-copy number replicon; $Km^R$. | [12] |
| pLT101 | pHH100-derivative with the *mCherry* gene cloned in between the NdeI-BamHI restriction sites. | This study |
| pUC19 | Replicative *E. coli* vector with high-copy number pMB1 replicon; $Ap^R$. | [13] |
| pUC19-BBa-Km | pUC19-derivative with inserted *aph(3′)* cassette conferring kanamycin resistance; $Ap^R$; $Km^R$. | This study |
| pRL101 | pUC19-BBa-Km-derivative with BsaI recognition sequence eliminated from the *β-lactamase* coding sequence. | This study |
| pJN101 | pRL101-derivative with inserted *xylS* in between the XbaI-EcoRI restriction sites. | This study |
| pXMJ19 | *E. coli*-*C. glutamicum* shuttle vector; pMB1 *E. coli* replicon; $Cm^R$. | [14] |
| pENZ004 | *E. coli* replicative, *S. cerevisiae* chromosomal integrative plasmid. | This study |

Yanina R. Sevastsyanovich, University of Birmingham) was PCR-amplified using primer pair mCherry-NdeI-F and mCherry-BamHI-R and digested with NdeI and BamHI. The digested fragment was then used to replace the *luciferase* gene in pHH100 resulting in pLT101. pLT101 replicates both in *E. coli* and *P. putida*, and also carries a functional kanamycin antibiotic marker and the *mCherry* gene. For the construction of pLT101-based GeneEE plasmid DNA libraries Gibson cloning method was used[15]. The pLT101 was amplified by the G-mCherry-F and G-mCherry-R that leads to amplification of the plasmid excluding the native *mCherry* promoter. The reverse primer mCherry-R carries the BioBrick Prefix[1], as a flanking sequence, that serves as a homologous sequence in Gibson cloning. This amplicon was used as the backbone and the GenEE(+SD)mCherry, carrying the *mCherry* linker, as the insert. The Gibson ligation mixture was made using equimolar amounts of purified vector and insert (totalling 100 ng DNA) in a total volume of 5 µL, added to a volume of 15 µL of Gibson assembly mix. The mixture was then transferred into the cloning host *E. coli* via chemical transformation and grown on LA plates supplemented with 50 µg/mL kanamycin. The resulting library was consisting of ~5,000 transformants.

##### 1.4.3 The functional screening of GeneEE plasmid DNA libraries in *E. coli.*

**Ampicillin screening.** For the functional screening of GeneEE plasmid DNA libraries for clones that confer ampicillin resistance, the library was plated out both on LA plates supplemented with 50 µg/mL kanamycin, and LA plates supplemented with 10, 30, 50, 75 and 100 µg/mL ampicillin. Among the obtained clones, 20 clones were randomly selected—8 colonies growing on 50 µg/mL ampicillin, 8 colonies growing on 100 µg/mL ampicillin, and 4 colonies growing on 50 µg/mL kanamycin—and plasmids were isolated with QIAprep Spin Miniprep Kit (Qiagen) and were Sanger sequenced (GATC Biotech, Germany) using the primer pRL101-Bla-Seq (Table 1). With these sequencing results, all the 20 constructs were confirmed to carry the cloned GeneEE segment (Table 5).

Table 5: The initially identified ArtPromU sequences in *E. coli*.

| Clone | AC[1] | DNA Sequence (5′→3′)[2] |
|---|---|---|
| Ec01 | Ap (100) | GTACTAGACTTCTCGTATCTACTGTGGCAAAGCGTGATGCAGGATGAGGCAATATAGGTTTGT<br>GTGGACTGCATCGGAATGCTACGGTATAGTGTGTTGGTACCGGCCTTCTATGGGTGGAATACG<br>ATCTGTCTTAAGTATAGCTGATAATGGAGGGGTAAGAGCCCACAGGCATCGTCGGAGTCGTAT<br>GATG |
| Ec02 | Ap (100) | CAAGAACAAAAAAGGCAATTATTGTCGCGCGCCATCAGTTTTAATATATATCAGCGTGGCTCG<br>TAATTGATGAGTCGTAAAGTTAGGCTCTGAACGTTCTGTCGGATTTGATGACTGTTGGTAAAA<br>TCCAGCTTCAGATAGTGCAGAGTTGTATGCTCAACTAGCTATTTAATTCGTAAGTATATCGAC<br>TAAGGAGTCGTGGCATGAGTATTCAACATTTCCGTGGTCGCCCTTATCCCNTTTGCGGCA |
| Ec03 | Ap (100) | CTCATGCCGACGGTTTTCACATTGCTTATGTATCCATTTTGTGGCCAGGAGTTTTCCGTTATC<br>ATTTTTTTACGAAATATCTCCCGGTACATGTTGTCTCGGGCTTGAAAGAAACTGAAACTGTAC<br>ACCTGTGTGGGGTGGATCTTAATAGGCTGGATGTGGTTGTGGTCTAGGAAGAGTAATCCCATA<br>AAGTTTTCTCGTGGAGTGTGGGGATG |
| Ec04 | Ap (100) | GTGATACCTAGTATCCTTATAGGTAGAGTTGGCTCTTGGAAAGCATGGGGCACTGGCGCGACG<br>TATAGGGTTTCATATACAATCCAAATAATCTGCCTTGCAGGAACTCCCATCTGGGAATTACCT<br>CTGGTGAGAAGTTCAGGGAAGGAAGTTGGGTCGGTGGAGATGGATGTGGCAAATGGCGTAATT<br>AATTGGAGGTCCGGGATG |
| Ec05 | Ap (100) | TGTTCATTCCGAGTGTGTGATGATTTTGTGATGGTTTGCAAATATGCTTCGTGTATTTTAGTT<br>GCAGATTTACTTGCTGCAGACTTAAGAGGCCATATAATTCTATAAAGAGCGAGGTGGCCTGGT<br>AGCCGGTATCTGGGTCTTGGCTGTAATTTTTAGAAGGTGGTGCCTAGTGGCACGACGGTACAT<br>GGGAGGAGGGGGGCTATG |
| Ec06 | Ap (100) | AGTTCTCTCACTTTCTCAGAGGTACTGTTGATTCTAGTTACACCAGGGTGCCTATTCTCCTTA<br>TCGTCCTAGCTCTTTAATTTTTTGTCAGTTAGTGTATTGGTCTTGGTTAAGGCATTATATTAA<br>TTGTTATCCAGGTGCCAACAAGATGAATACTTTGGTACAACCGGAATTGTTCCCTCGTGAGAC<br>TATAGGAGCATAGGGATG |
| Ec07 | Ap (100) | CTTTTTATTTTAACAGAAAGAAGACGTATCAATGTTGAGAGCCAGGTAACTGTTAATATGGGT<br>GTGTCTGCTTCATCGAGCCAAGGCTAGCATTGGTTTGTGTTTTTTGGTGGCAAATGCTGGAAA<br>GGATAGTAAGATCATCAAAGGAGGTAACGTGCAAATAACGATATGCGTTAAGGTTCTATTCTG<br>GGACGGAGCAGAGGGATG |
| Ec08 | Ap (100) | GTGGCACGGTCTCATGCCTATAGCATTTTTCATTGGATATTATATACTTATTATGAAACTTCC<br>CCACGTTGGATGAATTCGTTGATTTCGCTACGAGACTTTTTGTTCGAGGTGCTTAAGATAACG<br>GTGACAATATGGTTGTGGTGCACAGGATTAGAATTTCGGACTCTGATCCTCTTAGGGTAGTGT<br>GGGCAAAATGCTTCACAGTAAGGGAGCGTGGTCTCTCGTGATG |
| Ec09 | Ap (50) | CTCTCCGACCTGTTCTACGCAGGTGCATGCATATCTTGATTTTAAATAATACATGAAAGATCA<br>TTAGTGATTTTGAATCTATAAGATTGCATGCAACTACACTATCGGACTACGTAGTCATTGCGG<br>ATGGTCGTCAATGTACCCAGATGTACGTCGATCTGGGTTCGCGCACAAGCTTATATTTCTGAG<br>TCCAGGAGTTCTGTGATGAGTATTCAACATTTCCGTGTCGCCCTTATCCCTTTTGCGGCATNC<br>T |
| Ec10 | Ap (50) | ACCTCGCCAATGCGGATGAATGATTTACGACATTACGGACGCTTGTACCATTGTACTAAGCTT<br>GTGCTATCATGATTGTCGGATGATTTCTTTGCTGGCGAGGGGAACCAACAGGCGGCTATAGAA<br>GGGGTTGAAGAATTGGCACCAGTTGGTTAGTCGGCAGGAGTTTGGACCGCGCGAATCAAGCAT<br>GGAGGTGGCGGATG |

Table 5: continued from the previous page.

| Clone | AC[1] | DNA Sequence (5′→3′)[2] |
|---|---|---|
| Ec11 | Ap (50) | GATTTTCAGACCGTAGTCAATTGACGGTGGTTTGAATAAGGTATCCGTGATGTTTCTTGCAAG TATGTTCCCACCCAAAAGGAACTTTTTGATCGTACAATTCGTTGTAAGTTCATAATTGCTAAA CATTTAGTCTAAGGGCTCGGTATATATATTAGCGGACCTTGTCATTGAGTTCATGATTTTTGT TCAAGGAGTATGTGCATG |
| Ec12 | Ap (50) | CCGGCGTACTGACTAAAGATATTCTCATTCCATAAACGTTAACTGCCTAAACATGCATTTGCC TCTGTTATAGAGATGGAACGCGCTGATGTGGACATTAAACTTTGTGTTCTCGCCTTCGGTGGT TTTAAAGGGTATGCGTGTGTGATTGTGATCAAGTACAATTAACGAACCATGTTCGACAATCTT TCTAGGAGGAAGGGGATG |
| Ec13 | Ap (50) | GATGTTATAAGATGTTGCTTATTTGGTATCATCCTATATGTACACCAACCGGATACGATTTTA TGGAAAACTGGGGTTAGAGTGTGCTTTAACCCATTGTGAGCTGGCTGCACGGAGGTTGCCTAT TTTGATACCCGTACAGCCGTGCTCGTGTGGACTCGTATTTTAATGATCGGAGCCTACCGCTCG GCCATG |
| Ec14 | Ap (50) | GCAAGAGAATTTCCTGCGAAGTTCACGTTTTACATTAGGGTTTAGTGACTTTTTTGGCACAGC ACCTTATTTGTGCTACGTTGGTAAAGGAGCTCAGTTTTTCGTAGCAGAGTAGTTCGAAGGTGG GCATGTTCATACATACTTAACTCCGTGTTCCGGAGGTATTCTACGCTGATTCTCCATAAATGG AGTGGAGCGCGATGATG |
| Ec15 | Ap (50) | CAGCCTACTATTCGCTAACTAAGGCATAGTGATACCGATTAGCGAATTGTTGTTGATTATTCC GTTTCGTCAAATATAATTTTGCTTTACAAATTGGAAATCGTGGATCGCTTTTCATTAAGGGAA GAACCAATATGCTTCGGAGGGGGATGATCTGATAGGTGGTGGGTTAGGGGACTAGTGTGAGTT TGGGGGAGGCAGGGGATG |
| Ec16 | Ap (50) | GCACGGCAACTCCCCGTTAGAAATGAGTTAGCATACCCTCATCCATCCAGAGACTTGGTCTAG TTTCTTCCCAGGTTGTAATGTAACGTGCTATTAAAACGAGGTTGTAAAATTATGGGGAGGTAG GTTATGTCAGGAGATGTCAATGGGTTACGCAGGGAGGTCAAGCCGGTTATACGCGGTGATGCC GGGAGTAATCGTATG |
| Ec17 | Km (50) | GTGGCACGGTCTCATGCCTCTAACGGAGTGCGATATCGGGTCATTTCATCCTCGGCCAGAGTG GTGCGTATGGGTGTCAGTATAGGGCGTACCGAAATGTATCATTTTTTAAGCGAGCAAACGGTA CGGATTGACAGGTCCCGTGAAAGGGAGCGACGGCGTCGATGTGACGGGCTCGGGAGGTCTGAA TGCTGGCGGGTCGCCCGGTCTAGGAGAGCGGTGATG |
| Ec18 | Km (50) | CTCATGCCATTAAAAATAATTAGTGTACGGTCGTGGCATTGGCTCATTTTGTATATTGAACAT GTACCATGAAATCGTTCTTGCGGATGTTGGAGGCGGTCTCGTCGGATG |
| Ec19 | Km (50) | AAAACTGTGCGGGATTTGTTGGAGCATTCTTACAGAACTTATGACATGGATGGATTGTAGATT ATAAGAATTGGGCTTCTATCTTGCCTGTAACTCGAGCGGCTGTTTGCCTGTTTGGGTTCATGG ATGTGAGTTTGTGTGGACTTTGAGACATCGGTCTGCCAGGGCGTTGGAAGGGCGAGAGACTTG TCAAGGAGGGGGGTGATG |
| Ec20 | Km (50) | CTGTTGTCTTTCTCTATGCACTTTTTTGTATGTATCAAAAAACTCTAACAAATGTGATCTTAC CTTAATTATAAGCTTTCTGCGCGTCACTTATGGGAGGGTGGGGATG |

[1] The antibiotic concentrations used for the selection (µg/mL). Km, kanamycin; Ap, ampicillin.

[2] The ATG sequence at the 3′-end of each ArtPromU sequence is the start codon of the gene (*β-lactamase* in Ap, *kanamycin* in Km constructs).

**mCherry screening.** From the ∼5000 transformants, three colonies were randomly picked and plasmids were isolated with QIAprep Spin Miniprep Kit (Qiagen) and were Sanger sequenced (GATC Biotech, Germany) using the the sequencing primer pLT101-mCherry-Seq (Table 1). The sequencing confirmed the presence of the correct GeneEE segment within these clones. Upon this confirmation, 192 clones were picked that were appearing visibly red, with a varying degree of intensity to the naked eye, and inoculated into 2x 96-well plates containing LB supplemented with 50 µg/mL kanamycin. The plates were then incubated overnight under constant agitation, 800 rpm. The fluorescent measurements were made (excitation and emission wave length of 584 nm and 620 nm, respectively) using an Infinite 200 PRO fluorescence microplate reader (Tecan). The 192 colonies from the 96-well plates were also replica plated into 14 cm agar plates containing LA supplemented with 50 µg/mL kanamycin and were high-throughput sequenced (see below). The identified artificial promoter and 5′ UTR (ArtPromU) sequences with their corresponding phenotypes (mCherry fluorescence intensity measurements) are listed in Table 6. Based on this data a WebLogo[16] image was generated (due to the multiple alignment parameters only the ArtPromU with 211 nt long sequences were used) indicating the high-sequence variation within GeneEE DNA libraries (Figure 2).

###### 1.4.4 High-throughput sequencing of the GeneEE library.

To determine the ArtPromU sequences resulting from the GeneEE segments, a multiplexing strategy was applied. Two 96-well plates were arranged as one 192-well plate with 16 rows (A-P) and 12 columns (1-12). For each row and each column, one pool of cells was established, resulting in a total of 28 pools. For each pool, a sequencing library was created by two rounds of PCR. In the first PCR round, the GeneEE segment was PCR amplified using the primers Illumina-fwd and Illumina-rev (Table 1) in 15 cycles. In the second round of PCR, indexed sequencing libraries were constructed using the standard Illumina TruSeq primers. The indexed libraries were quantified using a BioAnalyzer (Agilent Technologies, Germany), pooled in equimolar amounts and sequenced in a 2x 300 nt run on an Illumina MiSeq sequencer. The reads from each library were joined with FLASh[17] using the following parameters: –max-overlap 450, –min-overlap 200, –max-mismatch-density 0.65, –allow-outies. From the 57,672 to 156,052 joined reads per library, the ArtPromU sequences were identified via search for the flanking regions. By counting the number of occurrences of each sequence in each library, the most abundant sequence found for a row/column pair was assigned as a ArtPromU sequence in the corresponding clone (Figure 2).

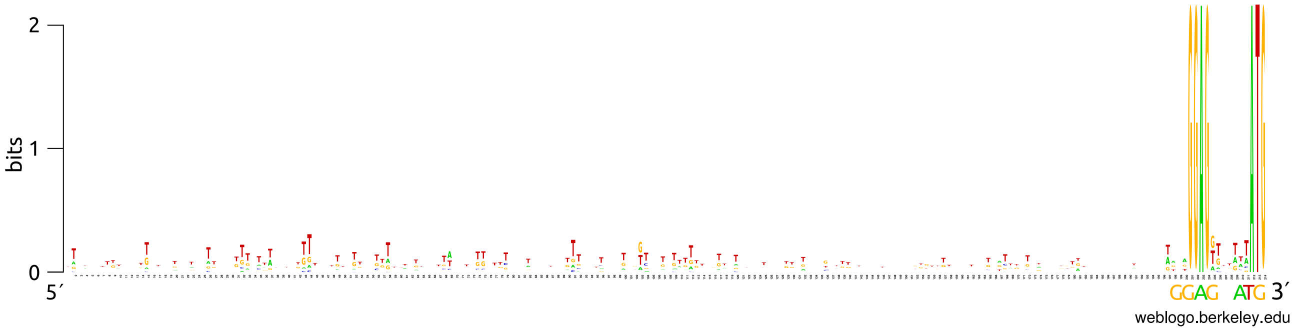

Figure 2: The WebLogo image of ArtPromU sequences identified in *E. coli* cells expressing mCherry at varying levels. In the alignment, only the ArtPromU sequences with 211 nt in length were used. For clarity, the defined defined Shine-Dalgarno sequence (GGAG), and the start codon of *mCherry*, ATG are depicted in larger font size below the alignment.

Table 6: The ArtPromU sequences identified in *E. coli* expressing mCherry.

| **Clone** | **FIM**[1] | **DNA Sequence (5′→3′)**[2] |
|---|---|---|
| EcE06 | 2246 | TCTCCTCATTTCTATTGAGTCATCCTAGCATTAGAGTTATTTGTCAGAGAGTACAGTATTAAGC GATTTTATGGTGTCCTAGGAGGTAGGGGTATGTACACAGTTCGTGCTTATAAAATGTGGCCTTT TCCTAGTGTTTACGTGCCGTTTTTTCTGCGACGGTGTCCCCAGATGCATTGTTTTGAGCGGTTG TGCTCGAGGGAGTTGCATAATG |

Table 6: continued from the previous page.

| Clone | FIM[1] | DNA Sequence (5′→3′)[2] |
|---|---|---|
| EcE05 | 2478 | ATTGTGTGGTTCTTATAATAATCCGTGATTTTAGAAGGTTAGATAGATATTTTAGTTCTTCTGA<br>TCAGGTACTGAACTGCGCGAGTAGCATATGTGTATTTGGGATTCCCCGGCACACCAATTGTGGC<br>GTGATTGGGGACGGGACTGCATCTATGTGGGGGACGTGCCGTCTGTACCATGCTGGGGATCGTC<br>CTCGGCGGGGAGGGCGGATATG |
| EcF04 | 3878 | CTTAGCGTGTTCGCGTGTCGAGTAGTGTAATGATAAATATTTTTAAAGTTAAAGCTGGTTTTAT<br>ATGTCACTGAGATTTTTATGTTCCTAAGTCAATGATGCACTCGGTTTGGATAGAGAGTCTATTA<br>TCCCGCAGATGATTGTCTGGAACTGCCTTTTGGGTTACTTCTAGGTGTTAGGCCTTGTCATATC<br>GATCGGAAGGAGGTGGTTTATG |
| EcB01 | 4202 | TTAATGGGGTTTTAAGTATTCAGCATTTTTTGCAGTGCATTATTCTTGTAAGAGAGTTCTGTTC<br>TTGGACGTAGTCGGGTGCTTTTGCTTAGATATTGTAGTTAGCTAGTTAGTGTTTGGTGTAGTCG<br>TAAATTATTTTTAGATTCTCTCAAATGACTTTTCGCATTCACAGTACTTGCTATCGGAGGTTAG<br>GTATG |
| EcA04 | 4854 | ATTTCTGCGATCCGGTTAGGAAATAACTTTCCTGAATGTGAAGGTGTTTGCCAAATGTTGTTGG<br>TTATTCCTCCGCATCGCACCACTTTTTTTGAGGTCGGGAAGGTCTTGTGGTTATCGCGCGCATA<br>CTGTCGTCTCTGGCATGTACCTGGTCGACGGTGGTGTCTTTGGTCGTAATGTGACAGTCTATTA<br>TGCAATGTGGAGCGTCGTAATG |
| EcM05 | 4925 | TTAGCGATATTTTCTGTTGTCATGTGTATGTTTTGCTAGATTCTTCTATTTCTAATTGGGTAAG<br>TCGAAGTTTTTATTCAGTAGGTTGGATTTGTTTGCAGATGTCTGGATGCTCTGATATTGGTTTT<br>TGCGTGATTCCTTGAGGCGCAATTCTGTTTCTGGAGATATCAGGTGCTTCGACATCGGGATGTT<br>CCTCTGTGGAGCGAGATAATG |
| EcG02 | 5191 | TTGACCCTATCTTGGTCTATTAAAGTGCAGGCACGTACATGTTTCTTGGAATTTGTTTATCGGC<br>GGTTTGGGATTTGTTGGGAACGGCTTTGCGGTTATAGGGTGTTGATATACTCTAATCGGTTACT<br>TTCCGGTCTAGGCTGATCTGAATATGGTTTCTCGGTTTATGTATGGCGTCCTGAGTGGGGCTTT<br>TACTGTTGGGAGGCGATCTATG |
| EcD05 | 5203 | GTGTCCTTTAACCAGATCGGCGTCATGTGCGTGTGAACGGAGTTCTGACTCGTATCTTCAAGGT<br>AACGGTTTTTAGACTATCCTTAGAGGGTGGGTTCGAGATCCCCAGTATGACTCGGGTAGCTGAA<br>ACGAGGCACGGAGTTTTTTGTTGCTTAGTGCGGTGGGGATTGGGTAGCAGGTCGTGGCAGTTCG<br>ACTTGCGTGGAGAGCGTCGATG |
| EcH02 | 5238 | TTGACCCTATCTTGGTCTATTAAAGTGCAGGCACGTACATGTTTCTTGGAATTTGTTTATCGGC<br>GGTTTGGGATTTGTTGGGAACGGCTTTGCGGTTATAGGGTGTTGATATACTCTAATCGGTTACT<br>TTCCGGTCTAGGCTGATCTGAATATGGTTTCTCGGTTTATGTATGGCGTCCTGAGTGGGGCTTT<br>TACTGTTGGGAGGCGATCTATG |
| EcE01 | 5511 | GCGAGCTTTTTATAGTTGTAAACTAAACATCTTGGATCTTGCGTCGTTCTGGTAGCTTCAGTTT<br>TACCTTGGATGTTTGGGTCTATTCTACGTAATCCGGTTGGCGGCCTTATTCCGGCGAGTAGTCG<br>GCTGCTACGTGCGGAGGTAAGTTATG |
| EcN08 | 5538 | TGGGAGTGTAAGTATG |
| EcH08 | 5590 | GTTAGTTTTTTTAGTGCATGTTGAATCGCTTTGGGAAGTCCTATTCTTTTTTGAGTGTAAGATC<br>TAGCGTTCTTTCCTCGTGGGGTCACTGCTGTCGGAGCTCGTTTCTATTTGATCTAACACGACCC<br>GTCTGCTCTAGTGCAGGCTCACGGAGTGGCTCGATGTCCTGTTAATATAAGCTTTATGTAGCGG<br>AAACTATGGGAGGGCAAAAATG |
| EcG04 | 5709 | GTATTGGTTTTATTTATGGCTAGAGCACAATTATGCTTATTGTCAGTTGTCCGAGATCGTAATG<br>TAAATTTTGTAACGGTTGGTTGGAGGTATAGGATG |

Table 6: continued from the previous page.

| Clone | FIM[1] | DNA Sequence (5′→3′)[2] |
|---|---|---|
| EcD06 | 5819 | TGTAGGTTGATTCTATTGAGTTATTGCGAGAGTGTGTGCAGTGTGGCTGGGCGGGATACGCACG<br>AATCTCTATTGACTCTGTAGCTAGTGAGTTATGTAGCGCTTTTTGTTTAAGACAACACGCCTTT<br>TCGGTTTGGGTAGTTATTTTTACAGTAATTGGTCTGGAGTTGGCTACGTGGTGGCGTTTAGCTG<br>CAACTCTGGGAGTGAAGAAATG |
| EcG08 | 5853 | TTTTTGTTTTTTACTGTCTTTCTGGGCAATGTGCGATCTTCTGTATATAGATTGTTCGCACTGG<br>TGTGTCGGAGGGTGTAAGGCAAGCGGGAGCACTTGTTGGTTTGTGCTCTGTAGTATATGGTTCG<br>GTAAATAGACCGCACGGTGGGAAGTCACTGTGGTTGTGGTGGCGAGGAGAACGTATGTCGATTT<br>TTTAAGTGGGAGGGCTGATATG |
| EcF07 | 5957 | AGTTCGCAACGCCCGCCGAGAAATTTCGATTGAGGTCTGGGCTCGGTATACCTGGTCATATCTA<br>AGTTTGTGGGGTGAGGCCTATGATCATTAAGTGAAGTAGTACAGGGGGCTCATTCTTTGTGTGA<br>GGTAAACCGAGGACAATTTAATCCGCCTGACGCTTACGGTTTCCTAAGCATTCAGGATATGTGG<br>GTGTGGAAGGAGATCATTGATG |
| EcM01 | 5978 | TGTGTACCCCACTTGGCCCTAGGCGTGTACACTGACGTTCAGATCTTCGAATCTATGCGCATAG<br>CAATATTTATTTCGAGTTAGGTACTTGCGGCGAAACATTAGTACTTGTCGGCGTATCTTCAGTG<br>CAATTTTCAATGTGCACTTGTATAGGGTACGTTATTGATTACGTCGTGGATGTCTCAGTCACGA<br>CCTTATGTGGAGTTATTTTATG |
| EcA05 | 6109 | CCGTTCAGTTTTGCGAGTACTGGGGTTTCAGGACGGATTAGGAGCCTTGTTTACCTGGAGCTGA<br>TAAACGTTTATCCGTAGGTATCGGACTCAGCTAATTTCACAGTGAATTTTCCGTTGCGAGTACG<br>TTATCGGTCATGTACGGAAAGAGCTTTTACATTCTTTAATTTGAGCAGCATGGTCTACTCTAGG<br>TCAAAGTCGGAGTGTTTATATG |
| EcD08 | 6289 | TATTAGTTTAGATTATTAGGCCTGTCTAGTTTCTCGTTTTGTTTTCACTTCTGTTTCATTTCCG<br>TTATTCTTCTTACGGTCGCTTGCGTCTGTGTCGTGATCATAAAGGAACTTCTGCGAGTTACCTT<br>TTCACCGCATTTCTGGACCTTAATTGTTGTTCTTTTTATTATTCTGGTATCAAGGCAATGAGTG<br>CTATAATTGGAGTCTGTAGATG |
| EcJ03 | 6685 | ATGATACGGTCCATTGCAGACGGTTTATACGAGGTTTTATGATTTCATTTTGTTACTTGAGTAT<br>CCGAAATGCTCTACGGCCTTGGTCTTCTTTCAAGTTCGTCACTCTTGTTTGGGTGCGAATTTGG<br>TGATGGGCGTTAAATTGCATCGGTGTTGTTGAACGTTAATTGTTTGAGATCTTTCTTTTTTCAT<br>TAGGTAAGGGAGGGGATTAATG |
| EcE03 | 6761 | CGGCATATGATTATTGGCAGATTCTTGTGTCTGTCCCTTTTTGTTGTGTGAATGTTTTTATCCG<br>CATTAGTTGAATGGCGCTTTCTCCTGTGTGTTATCTGTGCTATGTTCGTTACGAAGTCAACCTG<br>GCCATAATTCGGTTGGAGTGTGTTAACCGCGGTTAATGGGTGCCCGTATTGTGTCCTTCTTTAT<br>TAACTACAGGAGGGGCCTCATG |
| EcH07 | 7031 | TTGGATACGGTCATGGTACTGCTTGTGAAGTTTGTATACCAGTTAGGCTTAGGACCATACAATT<br>ATGTACGAGGGTTTGATATGGGATGATCAGTCAACTAGTAATTCTAGGTAAGTGCTTCCGCATG<br>GTATGACGCAGTGAGAAACCCCACGCTGTGGGGCGCTTGCAGCAGATCATCAGATACGTTTTTA<br>GTGCAGGGGGAGGTGTGACATG |
| EcB04 | 7129 | CGTTCTTGGTGGCATATGCATGTGTGACTGGTGTTTCCACCCTGGAGAGTCTACAGTTAATTTA<br>AAAGTAAGGTTCTGTCTTAAATTTTCGTTCTTTAATACGTTCCCTTTGGCTCGGGTTGAACCGA<br>AAATTGTGCTTTGTTGTACTCTGTATGGTTTCCTCTTATTTTGTCAAGCTTGATAACCCAGGTG<br>TACTGGTTGGAGGTTCTCTATG |
| EcC04 | 7494 | GGTAGTGGTAGATTCTCGTTTTTGAATCTGGATATGAGTGTTCGATATTGTTTGCTGGGTGGCT<br>CTGGATAGGTTTTTGTTTCAAGTTTTTCTCTTTAGTTGTTAACTTGCACCTTGTTTGACTTTTC<br>GTCGACATTCATGGGAGAGGATCTGGCAGATGTTTGGGTTTTAGGATTTTTCATTCAGTGCAAC<br>TTGTAGTGGGAGATCTACTATG |

Table 6: continued from the previous page.

| Clone | FIM[1] | DNA Sequence (5′→3′)[2] |
|---|---|---|
| EcN04 | 7510 | TATTGGACTCCGTATTGGCCTGGGCATCGCTTGAGTTCGTATCTTCGCACATATCTTAGCCGTT<br>TTATCTTAAGCAGTATTTGACGGATGTGCTTATGAATTGGCTCCTGGGCCTTTCGATTATCCTC<br>TGTTCTCATTATAATTGTCTCTCCGGTATTTTAAATTACTTGGGCTTCTGCCTTAGTATTTGTC<br>CTCGTAGGAGTAACATTATG |
| EcD04 | 7710 | AAGTGGTTGGTTCGTCAGATGTTGTATAATGTTGAATGTATTTTTATACTGTTACTCATCATCT<br>TTGCGGCCTTCACAATGAAGTACTTGTTATATTGAGTTGTTCGTTTCTTTCTTCCGTTTGTGCA<br>GGGACTGAGTGTCTTTATGTGTTTTTTATTTTCGAGTGCTGAGGCGTCAAGTGTGGCATTTTTG<br>GGTTGAGAGGAGGTTTTGAATG |
| EcF03 | 8034 | TGAATGAAGCAATTGATTTAGGTCCCGCATGGATAGGTAATGCTCCTCTTGCCATGTTTTTGCC<br>TTCTTGACCAAGTGGGTCTATACTTTTCGTTGTAATAATTGCATCTCTCTGGACGAACTTGTCG<br>GGCTATTGTAGGTTTTTTTCCGTGTTTGCTGTGCCTTCTTTTTCGGAGGACTAAGTGCCATACA<br>GGTCCTTGGAGCAGCAACATG |
| EcC02 | 8052 | TTCTTTAATTCGTCTCTACTGTCGCTTATTTTTTTTAAGTTTTTGCGGGAGATGTTGCTTTATT<br>TTGGGATTCTTGCCACACTTTTTTTCTTGCGGCTGCAGGTTGAATGTTGTGGTTCTAATGATTG<br>GGTGTGACATTATTGTCTGGTTTTCTCTTCCAACGGTATTTTGTATTTACTCTATTCGGCTATA<br>ACATAGTGGGAGGGTCACTATG |
| EcA06 | 8201 | TTGGATACGGTCATGGTACTGCTTGTGAAGTTTGTATACCAGTTAGGCTTAGGACCATACAATT<br>ATGTACGAGGGTTTGATATGGGATGATCAGTCAACTAGTAATTCTAGGTAAGTGCTTCCGCATG<br>GTATGACGCAGTGAGAAACCCCACGCTGTGGGGCGCTTGCAGCAGATCATCAGATACGTTTTTA<br>GTGCAGGGGGAGGTGTGACATG |
| EcF06 | 8469 | CTGGATTTTGGCGGAAGACAGTAGAAGGTGGTTCTAACGTCACCAAAGTATCACATTTATTTCA<br>TCATACGATTGCAGTGTGGCTACGTAAAAAGTGTAATGCATCGTTTCAGTTGCCCCCCGCCGGT<br>GGGATCAGTTAGGCTCTTATGCGGTTGCAGTCGCTATTGTCTCGGGAAAAGTCGTCTGATGCTC<br>AGCAAAGAGGAGCAAGAGTATG |
| EcL10 | 8479 | TTAAATCTTAAGGTTCTAGACTTGCTTCGACGTTTTTTTCGTGCGCCTCTTCAATGTACTTGTC<br>GAGTATTATTCGATTTATTGGTGTGTGTCCTTGGCTCTGGCGCCTTTTTGGCGGGATTCTTCGG<br>GTCTACCGACGGCCCGTGTCCTGGTTTAGTTCGTCGGAATTGTGGTATATTTATAATCAAGAGA<br>ATTTCTATGGAGAGTGTTAATG |
| EcA07 | 8597 | TTGGATACGGTCATGGTACTGCTTGTGAAGTTTGTATACCAGTTAGGCTTAGGACCATACAATT<br>ATGTACGAGGGTTTGATATGGGATGATCAGTCAACTAGTAATTCTAGGTAAGTGCTTCCGCATG<br>GTATGACGCAGTGAGAAACCCCACGCTGTGGGGCGCTTGCAGCAGATCATCAGATACGTTTTTA<br>GTGCAGGGGGAGGTGTGACATG |
| EcA02 | 8950 | TGGTGATAATTTGATGGCGATGTTGGGTTTTCTATGTCCTCCATGATCTGTATAAATCATGCGA<br>TCTGTGCTCTCTTATTGGTAACAATTTTACTATCGCTAAATGGGCTCTTTGAATGAACGTACAA<br>GAGGAGATACCCGATG |
| EcK01 | 8969 | TTTATATTCGGTTATTCCATCGAAAACCTCATTATAAATGTTCTTATAGGTAAGGCTTTAACAT<br>CACTAGTTTGTGCTTCACGGATTTTTCATGCAATGAGGTCGCTACCTTCTCAACTCCGGGGTGT<br>CATTCACGGCAGCGGGGTATGATGGGGTTCAAGGGTTAGAATTGGTTGGGGGTTGCGAACCTAC<br>CTAGTAGTGGAGGAAAATTATG |
| EcJ01 | 9285 | TCCCCGGATATTAACTAGTATCGGAATTCACTTGGCCGCCCCTCGGAGTGATTACATG |
| EcA01 | 9352 | ATAACGGACAGGATTAGGATTGATCAAAGTGGTACCGTGATTCGTAAGGTTGATGACTTACATA<br>CCAAAGTGCTTTCGCTTTTTGCATTATATGAGTCCATTTTAGGGAGTATCTCCATG |

Table 6: continued from the previous page.

| Clone | FIM[1] | DNA Sequence (5′→3′)[2] |
|---|---|---|
| EcB07 | 9580 | TGAGTCATCTATACTACCCATGGAGTGTATTTGCGGAGCGTTGCTAATTTGGCTGGCAGTTCTG |
| | | CTTGACTTGGTATCTCCGCCCTTAGTCGTCGACTATACTATGGAGATTCATGCATTACAGTATC |
| | | GCTAGAGTTGAGAGGTGACCTGCTCTGGCCCCGGATCTGGTTAGAAGCAGGCTATACCAGACGT |
| | | CGTATAGTGGAGGGCTATTATG |
| EcD01 | 9585 | GATGTTTTATTGAATCATTCCAAGGGCATATTTGGGTCCCGATTACTCGTGGACATACCAAAAC |
| | | GATATGTTGAGTCATGCACGATGTTACATCATTCGTCAGTCGATATACGTTTTTAATGTTTACG |
| | | GTCATCGCCGGGCTTAATTTTTTGTCTGTTCGACAATGTGATAAGAGGATAATGTGAATAGTGG |
| | | TTTAGATGGAGTCGTGGTATG |
| EcF01 | 9825 | TTTCACATGATCCGTAACGTCTACTGTCGTCGTGTGGTTAGGGGACGTAAGAGTGCTTAGCGTA |
| | | TACATGGCCGCTGGACTGGGAGTCAAAATGCTCAGAGTATGTTCGTTCTCCTTTTGGATTCGGC |
| | | TACGCCTTGAATCCTGTTCAGTAGCCATCGCAACATTATTTGGTCACATGTGAACATGTGTTAG |
| | | GTATGTAGGGAGCATGAACATG |
| EcE12 | 9955 | AATGGGACTTTGCGCTCTTTGTTTGGATCTATGCTCAACCGAGTACCTCTATAGTTCGGGAGGT |
| | | GGTTAACTATGAGAACTCGTATTCTTAGAATTTTATGACTCTACGCTTTTTTATTCAGTGACTT |
| | | CACGTGACACAGTACTTTAGAGTTGAGGTGACGTCTGTCTGGCATGGGGTATGAATTTGATATT |
| | | GGGTTTAGGAGAGTAAGAATG |
| EcB08 | 10069 | GAAACTTTTTGACAACGTTAGGTTTTTTAGTTCCTAGTGTTGAGTGTTATGCTTTTTCATAGGC |
| | | TTTGGTCTTAGTGGCGGCACGTCTGTGTTCCATCATTTCCTTATGCGTGTTTTTCGGGGTTTGC |
| | | TTCTCTGGAGTGTGTACCAGGGTGGTCTGCCTACTATGAACGCTCGGTACTCTCATGTACTGGC |
| | | TAGTAGAGGAGTGAGTATATG |
| EcO02 | 10312 | GCTTTGGTTCGGATTACGATGTATCATTCAGATTAGGACTAGGCTGTAATGGTTCTTATATGTA |
| | | TGGCCTTAGGGTGTCAAATCTACTATCTACTTATCTCGGTGCTAATTATTATTACAGTTTGTGT |
| | | TTGTCACAGTATCTACGTCCTCAATATCGTTTGAGGCTTCTATCATGAGATTTTTAGTTATGGA |
| | | AATCGGAAGGAGTGTTTTGATG |
| EcM09 | 11068 | GACGGCATCACAATGTTATAAAACCATGTCTCGTATGCAATGGATTGGTTCTAGCTTATGTCTA |
| | | ATTCAGCGGGTCCAGGGTTGAAGTGACCAGTGCCTTTGTTCTACGATTCGTTTTATGTTGCGAG |
| | | GATGCATTTGACGATATGCTTAAGGCGTATCCTAAGTAAAATTCCGTTTGTCATTGTGCTCGGT |
| | | CCGAAAAAGGAGGGGTAGGATG |
| EcD09 | 11204 | CGTATATCTGAGTATTGATCCTGGTTTATTTGACTTGTTTTTGGTGTTTGTAGTAAATTATGTC |
| | | TGCGATGTTGCGACTTTTATTAAAATCGGTTTCGAGGGCGAGTTATCGTTATTTATTTTGTGTG |
| | | GATGTTGGAACATGTCTAGGCCAGTTCTAGAGGAATGGCGTTCTCGCCTCCTCATAGAAACTGG |
| | | GAGGCGTTTTATG |
| EcF12 | 11322 | TATCGTTCCAAAGGTCTTGATGTATTCATTACAATTTTTAGGTATTCTTGGTGACGGTTCGATG |
| | | GGTTAGGCGATCGGTGCATGTAGTCTAGGTCTCTTGCTGTATTTCTTCGAAGTAGTGTGCTTTC |
| | | TCGTGTGTGTTATTAAGTTTACGTGTTGGCGTGCTATCTTCCTCATCAGCTCGTCATAGTTCAA |
| | | GTTTGGGAGGAGTAGTGTTATG |
| EcH06 | 11837 | TTGGATACGGTCATGGTACTGCTTGTGAAGTTTGTATACCAGTTAGGCTTAGGACCATACAATT |
| | | ATGTACGAGGGTTTGATATGGGATGATCAGTCAACTAGTAATTCTAGGTAAGTGCTTCCGCATG |
| | | GTATGACGCAGTGAGAAACCCCACGCTGTGGGGCGCTTGCAGCAGATCATCAGATACGTTTTTA |
| | | GTGCAGGGGGAGGTGTGACATG |
| EcN10 | 11922 | CTTCTCGATTCCTTATGTTGGGTCCGTCGCTTTAAGGCGTCTCTTTCTATTGGCCCTTTGATTA |
| | | CATCAATCATGGGTAAAATTTCATATAAGTCTGCTAGTGTAAATTTTGTAGGTTGTTGTGAAAG |
| | | ATGTTATTTTCTGTGTTCACATCGAAGGTATTCAAGAGTTAGTTCTTAGACGGTAATAGTTCAA |
| | | AGGTTACGGGAGATTGAGTATG |

Table 6: continued from the previous page.

| Clone | FIM[1] | DNA Sequence (5′→3′)[2] |
|---|---|---|
| EcI10 | 12196 | GTGATAGAGCCATAACAGCTCGTCTTATCGATGGGGGCTGTACTACTAGCAGGAGGGACCTGCC<br>TCGTTGATACTGTGAAATGAGATATTTCTGCTTGGTCCGGCGAATTGCAGTCGTCGGGTCTGTT<br>TTTGTAATGGAGTTTGAGAATG |
| EcH05 | 12237 | CTTCTTAGGGTGAAGTTTATTATCTACTCATCTGCGTTGGGGGTCTCAGGTATATGGGTTTTGA<br>GGCTATATGTTGGGTTATAGGCATCATTTCTCCTCTTCGCGTGTTACTTTGAAAATACGTTCTA<br>CGATATCACCTCGAACGGCATCGGGGTCTTGGCTCACCGCAAGGATACGGCAACCTATGTTAAC<br>TAGATTTTGGAGACGGATTATG |
| EcA08 | 12555 | CATATTTGGGAGTGTTAATATG |
| EcN06 | 13303 | GATCCCGCAGTGTGTGCGTTCTCAGTTTTCTGTAAGAGAAGGAGGAAAGTGATG |
| EcG07 | 13397 | GTGAGGCCATCCCTGATGTGTATATTGTCCTGTGCTTATTTGGTACGTAAGGACCGTCTCGGTT<br>AAATATTCTAACCTGGATTTGTATCCTTCCGGTGTTTGGCTAATGTTAGTATGATGTGAATGCT<br>AGAGTTTCTGTTCAGTATTGGTTAGTTGGTATAATCCAAGGTTGAACGGTAGTTAGTGGGCTCT<br>GAAATTTGGAGGGGATCAATG |
| EcC08 | 13622 | GTGGTAATGGCGTGATTATTAGTTATGTGACGGGTTGTACTATTCGGGTATCTAACTTTTGTGT<br>GAATCGTTCTTGTGTGTAACGTTTGGATATATTATTGAGATTGGATAGAGTTCCGTTAGTGTGT<br>TGAGGGAGGTTGTTACATTGCTTGGTTGTATTCTGGTGAGTATGAGTTTCTTGTATATCTGAAT<br>CGATTACTGGAGGCAACTTATG |
| EcO08 | 13745 | GCGTTTGGTTGGCCGTGAAGACAGATTGACTAATACTGTGCACCAGTATTGATGCTAGCTGGAT<br>GCTGAGCACTCTACCGGATCACTTGCGCAAATAAGTCAGAGGCCTTTAGTCTGCCTTAGGTATT<br>TGTTTTTTTAACACGCCACTCTATTGCTTATACTGACTGTCACTGACTCAGATTCGGGTTATCC<br>GCTCTACGGGAGGAATATTATG |
| EcO04 | 13770 | AACACCGCGGCCGGTGCGGTTTTGGTCTCTGTCGCGTATCCGTTTTTTTATTCGACGTACACTG<br>CTTTACAATCGTTCTTGTAACAAGTCGTCGTTTGTGTCTTGGCGATTTTCGGACTTTTGTTGAG<br>GTTTACCTGAGTTTACGACAAGGGCCTTATGATTCCGTTAACTTGTGAATGATGGGGGTTGGTT<br>GGCTTTATGGAGGTACGATATG |
| EcF08 | 14096 | GAGGGGTACGCGGGTTACATTAATTCTCTATCCGTCCTTTGTTTCACAGTTGCCTATATTTAGT<br>ACATTTTTATTATAATTAAGTATGGCTTCCTCGAGTGGTTCGGATTTGCTTCAACTGTACTTTA<br>ATTGTGGTAATGGTATTTTTTAACAAACTTAGGGAGCTTACCTGATGTCTGTAATCCATACGTA<br>TATCGAGAGGAGGTATCTAATG |
| EcM03 | 14281 | TTGATTTATCATAGGGTATGTATGTATGTTTTAACGTTTATGGAGTACCAGAGCATTTTCTTTG<br>TCTTCTGGAGTTATTCGCGATTATCTTATTCTTTAGGTGCGGTTCTATGTGTACGAATCGTGGT<br>TGTATTATTATGCGATTCATTACGCGGGGTGCGGCTACTCTGATGCGTGCATTTTGTATTTTAT<br>CATGATCTGGAGGTATTAAATG |
| EcN05 | 14339 | ATAAGTTTTAAAGGGAGGCGCAGTCAGAGGTTTCTCTTTTTCGTCGGATCGACTTTTCGTGTAA<br>TGTTTTCTACCGGGGTGTAATATAAACGGAGCTGAGTTCGTGCGTACAGTTTACAATGAGATAT<br>CTCTGGTTTGTTGGACTGAGCTAGAGCTGGTTGCAAGGTCAGCGTGGTGCTGATTTGCCGGAGC<br>GAGGACGGGGAGTGCTGTTATG |
| EcG12 | 14434 | TACCACGTGTGGTCCGTCGTGGGTGGGTGCTATAGTTCAACGCAAGCAATTGCCTGAATATACT<br>GGTTTGTTGGGTTCTATACAAAAAATGTTTAATGTGATGAACGGATGTTTCTTAGGGGATGCCT<br>GACATCTTTCACCAGCATTGGTCGAAGGCAATTTGTATAGTGGTTGCGTGCTTCGTAGGAGTCT<br>TAGTTTGAGGAGTATCTCGATG |
| EcD07 | 14526 | ATTACATTTTCGTTTGGTTTAGGCACGTGAATTTCTACAGAGGTTGGAGTGGGTATATGTAATG<br>CTTTAGATAGTATTGCATAGCGGCTGTAGCACTTCTTCGGCGTGACCTGTAGAGTTTGGAGTGG<br>GCGGGTCAATCAAAGTCAAACAGGGGTTGCCTTATCTGAGTTGTATCCCGCCAACCCGTCCATG<br>CAAATAGAGGAGTTATAAAATG |

Table 6: continued from the previous page.

| Clone | FIM[1] | DNA Sequence (5′→3′)[2] |
|---|---|---|
| EcE04 | 14650 | GTGGAAACGACGGCCTGTTTGGACTCTCATTTGTTGAGTGTGATATGTTTCCGTTATGACTATG GGTGTCGTTTTGAGGTGCTCATGCATTGAATGAATGTGATGGGTGTCATGATTTTTGATTCTGA GGTTGCCCCTTTCAATGTTCGGTTCTCGGCCACTTTTGTACTGTTATTATTTCGAAATACTTGT ACTCTGAAGGAGACGATTGATG |
| EcD02 | 14690 | AGCTGCGATGGGGCGAGGGAGACTGCTTGTTGTGCAACCCCGTGCTGGGTCTAATTTTGTTAAT AAGTTACACGTGTTCTGCTGCGGAGCTTGTACGGTGGTATCAAAGGGGTTAGGAGAGTAATCCT AGATATTCGATTATCGACTGGGCGTGAGCTGTGAAGGTGCGGTGCACGGTCAAGGGCGCTCGCC AAATGTAAGGAGCAAGTGTATG |
| EcG11 | 14786 | ATTTCAGGTCCAGGTTTCGATGGCGGGAGGTTATTATATTGTGCTCTTCTATCCATGTGTTGGA CTGCTTCCTCGACTTGACTGTCGCCGTTTTAAGGTCTTGTGGGCTCATGGCGCAGTCGTTTTTC TAGGTTTACCTTTACCTTGCTAGGTTAAACTCCTTAGTATTGTTTTCATTGGCATTTGGCTCAC ACTCATATGGAGATATTTCATG |
| EcG01 | 14881 | CTGCGGTTCGGTGCCTCGCTTAGTGAGTCTCTGTTTTCTCCGTTGGAAGTGGGAGACAAATGTT AGTTTATTCTATTGTGTGTGTAGGCAAATAATTTCTCAGGCGTTTGAATCTTTTATCCGTGGGA CGGGTTAGTGCGTATTATTTAGATTTAATCTCGTCAAGGATGTTGCTTGGTCCATGGATGAGTT ATTATAGGAGTGTATGTATG |
| EcB05 | 14998 | TGGGAGTGTAAGTATG |
| EcK11 | 15406 | TAAGAAGTCTTGGTGAGGATCTCATTAGACTTGCCCCGTGCTAGTATACCTCAGTAACGTTCCT GTTTGTGGAGTGATATGATG |
| EcH03 | 15512 | AGGTGAGTTATTTTGATCATGCGCATTTGTAGTATATTTTCGCGTTTAAGTTTGTGTTCCTTAT TGCGCCCAGTGATCTTCTTATGTCAGTGTACATCACAATCACATTTTGGCTGATATTCGTGGTT AGGTTGTTGTTTTTTCTGTAAACAATAAACTAGTTAACTAGAGTTATGTGGTTTTATAACTCGA TGGAGTAAGGAGTCAGTAAATG |
| EcJ07 | 15530 | ACAATTTCATTGGGGAAAGAAGTACTCTAAGGCGCATATATATTTGTTTTCGGTCTATAGTCTA GAGGCCCCTATGATATGTGTCTAATCTGGGGTCCTTGGACTTTACGTTAAGGCTATGAGTACAG TTCAAGAGAACGTCCAATGTTTTTGGTACTTCTACTGGAGGAATTTGTTGGAACACTTTGTGTT ACGTACTCGGAGGTATTGCATG |
| EcL06 | 15943 | AGTTGTTGACAGAGATTTCTAGCGTTTCCTAACGATGACGTAGTCAAATACGAATAAATCCAGG AGTCACAATATG |
| EcM08 | 16115 | GCCCTGTTTCGTTAGGATCCTAGGTGGAGATTATCTACATTGAATCTAGATGGAACTTATATAG CTCGACCTTTTCGTTTAAGTGCTAGTGGTTAGTATATGTTACCGAATTGTTAGTAAAAGTTACT CGAAGGTCCAGGCTCTAGTATTCCGATTTCAGTAATCTTAGTTAAACGCAGCGCTTTGCGTGCT AATCAGGAGGCATAATATG |
| EcL08 | 16228 | GCCCTGTTTCGTTAGGATCCTAGGTGGAGATTATCTACATTGAATCTAGATGGAACTTATATAG CTCGACCTTTTCGTTTAAGTGCTAGTGGTTAGTATATGTTACCGAATTGTTAGTAAAAGTTACT CGAAGGTCCAGGCTCTAGTATTCCGATTTCAGTAATCTTAGTTAAACGCAGCGCTTTGCGTGCT AATCAGGAGGCATAATATG |
| EcF05 | 16326 | TTTTCAGTACGCACAATCTAGGGGGAGAATTTTAATG |
| EcE09 | 16349 | CACAGTTAGTCAGTTATTATTTAAATTGCGTCCCCGGTAGTTTGGCACGGCCTGATTCATATTC AGAGCACTGCTCTTCATTTACTGTCTTGCTTACCCGCCGTTGCGTGCTATCCTTGAGTTTATGA AAGGCGGGTTACTTACGCTTATCGGCGGCGAGCTGCACGGGCGCTTGGGGGTCCCGAACTCATT GGCTTATGGGAGGTGGTTTATG |

Table 6: continued from the previous page.

| Clone | FIM[1] | DNA Sequence (5′→3′)[2] |
|---|---|---|
| EcN12 | 16752 | CATAATTTTTATAAGTTTAGTATTGTGGTTGATCAGTGGTGAATAAATGGGCGCACCTACTGTA<br>CCCTATTTTATAAGTTGCGAAGCTATAATGGACGGGGCATGGTTCTGTTTCGGATTGGCTCGCA<br>GTGTGCGGGCGACAACAAGAATTAGTGGTGCGGGATGAGTACGTCTTAGGTTGGTTTTAACGTA<br>TCCTCAAAGGAGTTTCGTTATG |
| EcL02 | 16808 | TCTGGTTGCAGTTGAGACTTAGTCTAGCTTGTTCTGCCAAGTGGTTACAGATATTTGGTCATGG<br>GTACTTAGTGTGCGTGACATGTCATGTCCCAATTAGGTGTTTGACCTGTAAGTTGACTTATGTA<br>CTGGCTACGGCGGCGATTTATTTTAGTCGGCACGCGGAGTCATTGCAATAATGGGACCAGCTAT<br>GGTCATGGGGAGTTTTTATATG |
| EcL04 | 16810 | AGCGTGATAATGTTCGATGCAGGTACTACAATTACAATGATCTGGTTTTCAACACCTGATTTTT<br>ATAATGTAATTGTATAGTTCGTTCAGTTTATGTTCTCAATTTTGGTCTTTGGTTGTTTTGTGGG<br>TTACAACTTCAGGGTAGGATTTCGTGCTTAGTGTAATTGAGGATGTCTGTTTTACTAAAATGAG<br>GATTTGGTGGAGAACGGTTATG |
| EcC11 | 17143 | GTCGAATTTGTTCATGCTAAATAGGTTTTCAGGGTTGGCTGGGTGCATTCATTTCGTACGTTGT<br>CGCCCTCAACGCATTATATTTAGTCTATCTGTAAGGGAGCACTTTTTAGGGCTGGTTATCGCCT<br>TGTGTGGGCGTGTATGACCGAGCGAATGTGGTCCGTAAGCACTCGGTAAATGAGAGAGCAACGC<br>TATAAAGGGGAGTTCAACAATG |
| EcA10 | 17359 | TTACTATGAGTGTGAATCGCGAATGGTCGGTTATGAACTGGAAGAGTGGGCTTAGTGTTGATAA<br>CGCTATTTATGATATAACCAGACAATTTTGCTTGTTGTTGGGCCACTTCGAAGTTTATCTTATA<br>TGTACCGGTAAGTGTTTCTTTGGCCAGACTCCATTGCTATATAGGTGATGTCATGTCACGGTGG<br>GTTAATTGGGAGATTTGATATG |
| EcH11 | 17695 | GTTTATAACTTGCATTGTGCTCCCTCTTAGTGGTATTTGTTGTTATCACTTTTAATGGGTCTTT<br>TCTTCTGGGAAACGTATCTTTTTTTATTCCTTGAACGTTTTTAAGGTGTTCAGGGGCTCAGCGA<br>TCGTGCTTCAGCTCACGTTTCAACGGCCCTGATCAATGAAATCTGATGTTTTGATAACTCGCCA<br>GGCAAGTAGGAGATGAAAAATG |
| EcC05 | 17734 | CATTTGACAGTTTTTGTATGAGATGAATGGTGGAGTTATACTCTAATGCTGTAAGTGTGTAGGC<br>CCTAGTTTTTGCTTCGTTCTATGTACCAGCGAGTTTCTTTAACATTCTATTTCCATAACAAAAT<br>TTGTTGTTCTTGTTCTATAGGGGGCATCCTTTGGGTTTGTAATTAGTCGTATAGTCATCAAGTA<br>CTATCTTATGGAGGTAACTAATG |
| EcC06 | 17933 | GGCAGAGCGTTCTTCGCTGGAGCTAGAGACGTACTTATGGATAATGAACTCATCGTCTGGATAT<br>GTTTGATTTTCTACTAGGATGATCTGCTGCTTTCAGAGGCGATTTGTAACTATAAGGTACGTCT<br>TGGATAGGTTTTAGAAGTGTGGTCATATTTTCGGGACCGGCGTTAGGTTTTATGAGCCTACAGA<br>TGTTATCCGGAGGTTTTTTATG |
| EcF10 | 18030 | ATCTGGTTGAACGTTGTGATTTAAAATACTGTTTTAAGGTTTTGACGTTCATTCGCATGGAAGG<br>GATCATGTTTGGGGCTGTGGGAGTTTTTATCAGTGGGAATGTACTAGACTCGGTGATGTACCAT<br>GTGACTAGGTTCAAGTTTGAAATTTCTTGTGCATGGGTGCAGACGCCTGGGTTAACAGCTATGT<br>AAAAGACGGGAGAGTAGTGATG |
| EcP10 | 18079 | ATTACATTTTCGTTTGGTTTAGGCACGTGAATTTCTACAGAGGTTGGAGTGGGTATATGTAATG<br>CTTTAGATAGTATTGCATAGCGGCTGTAGCACTTCTTCGGCGTGACCTGTAGAGTTTGGAGTGG<br>GCGGGTCAATCAAAGTCAAACAGGGGTTGCCTTATCTGAGTTGTATCCCGCCAACCCGTCCATG<br>CAAATAGAGGAGTTATAAAATG |
| EcN11 | 18082 | CTTTCCGACATCGTTTGCTTGGGAATTATGGATCAATTTATTGAGACCTGTTTAATAAATGCTG<br>TGTAAAACATTATCATTGAGTGTATATTAGTTGTTACTTTCATCCTAGTCTCGATTTGCTTTAA<br>TGGCGCGCTTTCTTTTGGGATGCATGTATTGGTTTTTTATACGCTTATCCGCTAATCCGTTTCG<br>TACGTTGAGGAGTTTCTCGATG |

Table 6: continued from the previous page.

| Clone | FIM[1] | DNA Sequence (5′→3′)[2] |
|---|---|---|
| EcI02 | 18092 | TTGGTTGGTTTGGCACGCCTGGGTGCGTATATGCGTATAGGCTCTTTGACGATTAAATATTTGA<br>GGAGGTGACGGTGTTATATCATGCGTCTATCAGTTCACGCTGCCTTGCGTCTATATTAGTGCGC<br>ACGGGTTATGTTATTTGGTTAACACGGGCTTATGTGTTTTAGCTCGATCTTTTAAAAGTGATTG<br>TGCTCGTGGAGGTTTCTAATG |
| EcK02 | 18161 | GATATCGGTACTGTTGTGAATGTTTTCACCTTACGTTGTCTAGGTAAGTATTAAGGGTATTTTG<br>ATGTTGAGGGGTTATAACAAGCGAGGCATTTGATTTTCGTCGAAGCCGATCTATTTCTGTGTAT<br>CTTGTAACATCTTTTGGTCAGTAAGTTGTAAGGCAGGAGTATTGTGAAATCGATGTGTGGCGTA<br>GAACTTGAGGAGAGAGGATATG |
| EcF02 | 18299 | TAAGTCTATTGTGGGTCATCATTGGCTGGCCGCTATATACTTTTTATGAGTAGCCTTTCAATGT<br>TTTTTTGTTTTAGGCTCAGGGTTTTGGATATAATAGCTTGTAGTCTATGATGTAGATATGGACT<br>TGTGAGCCTAATGTATAGAGTGTTCTTTGTTTGGGTATATGGGTGCTCAGTGTGCAAATTGGGC<br>CAATTGCTGGAGGATGTGTATG |
| EcE08 | 18320 | ATGCTAATATAGGAGTCCTTTTATG |
| EcL03 | 18471 | GGCCCTGATTTATATTCGCTGATGACTATGGGTTAATAATTCGATTGTTTACTTAGTTTACTGT<br>ACTCATACGTATCATAAGTTAGATTGTTTAATCGATTTTCTGGAAAGAGTTGTGCTACACTGTC<br>CGGGCTTGAGGATATTTAGCTCTCAACTGTAGTTCATTTTTCATATCCCTGTTTACTTCTCCGC<br>ATGTTGATGGAGAGGGTCTATG |
| EcO10 | 18586 | TATTCAACGTGTCTGGATTGGGAGAACTTTAATG |
| EcN01 | 18642 | TGCTTTATGCGTGGTCGTTCACGGTGAGTTGTCTTGCTGTGTGGGCATATTGTTAAGGTCTGAC<br>TCTATCGTTGTATTTATGCTGGAACCAGTACTGAAAGAGCAAGTATGGTGTTTTTCAATACTAT<br>TGTGTTGTAGATTTGGATTCACTGTGTAACTCAATTGTGGTGGGGAGTGGGGTTTCCCGCCCCT<br>GGCGACGTGGAGGTTCTCTATG |
| EcB11 | 19087 | GTCGAATTTGTTCATGCTAAATAGGTTTTCAGGGTTGGCTGGGTGCATTCATTTCGTACGTTGT<br>CGCCCTCAACGCATTATATTTAGTCTATCTGTAAGGGAGCACTTTTTAGGGCTGGTTATCGCCT<br>TGTGTGGGCGTGTATGACCGAGCGAATGTGGTCCGTAAGCACTCGGTAAATGAGAGAGCAACGC<br>TATAAAGGGGAGTTCAACAATG |
| EcC09 | 19162 | TAGGCATTGAATCTTCTTCTACGAAATTTGGTACTTAAATCATTTAGTTTTTTGTGTTGGATTT<br>CGACTCTTGCTCTTCTTTGCTGTACGTTTCTCTTCTGTATCGTCTAATTGTCGGGTGGGTTTGA<br>GTTTTCCTATTTTGTTTGAGTCATTGGTGACTTTTGCTATAGTGGTTAACCGTAAACGAAGTAG<br>GTCAATCCGGAGTATCGCTATG |
| EcO06 | 19187 | GGCAGAGCGTTCTTCGCTGGAGCTAGAGACGTACTTATGGATAATGAACTCATCGTCTGGATAT<br>GTTTGATTTTCTACTAGGATGATCTGCTGCTTTCAGAGGCGATTTGTAACTATAAGGTACGTCT<br>TGGATAGGTTTTAGAAGTGTGGTCATATTTTCGGGACCGGCGTTAGGTTTTATGAGCCTACAGA<br>TGTTATCCGGAGGTTTTTTATG |
| EcK10 | 19390 | CGTGCTTGATCTTTCAGCATACTTCAGAATGTTCAATTTATCGTATATTACTCGTCTCGGTCGT<br>TAAGATGGTTAGGTTTTGTTCTAAGAAGTATTTGATATACAACGGGGAGTTTGATTATG |
| EcK08 | 19413 | ATAGATCCTTTGTAAGGATTCGTTGGACTAGTTGTTTTTCTCTAATTATATTGACCCATTAAGC<br>CCATCTTCTATGTCTCAGGCTCTGATACCCATAGCTACTCTTAATAGAGAGGTAGCGTTGGTAT<br>CCGCTACGTGTCAGCCGTAGCGGGAGGTGAAAACAACTTGGTTTAAATGGGTATTGTGTATAGT<br>ATTGAACAGGAGTGTTTGTATG |
| EcI05 | 19623 | GTGTATGTGGGTCTTCGCCATACGGTCTCTAGCGCATCAGGATCACGCGATGGTTGGTTTACGG<br>TTTCCAAAAGGGGAGGGTGTATATG |
| EcL11 | 19871 | TAGATGGGAGAATGATAATG |
| EcP04 | 20219 | TGCTTTTTTATAGCGGGAGACTCAATATG |

Table 6: continued from the previous page.

| Clone | FIM[1] | DNA Sequence (5′→3′)[2] |
|---|---|---|
| EcH12 | 20439 | ACTAATGTGAATAGAACTCGCTCTTTGCGACCTCTAGATGATCTAAGTTTATTTAGTTAATCGA |
| | | GTATACTAGGTTCGACTTGAATGGTATCGTTTCGTCGTTTACGACGAATTAGAAAACGTGGTGA |
| | | CTTGCGACGCTTGTTGTAAAGGTTTTTTGTCGATCGAACTTGAGACGCCTTTGTGGGCTCCCA |
| | | TAGCGATGGAGATATTTTATG |
| EcI09 | 20611 | AATGCGTGTATGTGGGGAGGTTTAGGATG |
| EcO09 | 21000 | GGCAGAGCGTTCTTCGCTGGAGCTAGAGACGTACTTATGGATAATGAACTCATCGTCTGGATAT |
| | | GTTTGATTTTCTACTAGGATGATCTGCTGCTTTCAGAGGCGATTTGTAACTATAAGGTACGTCT |
| | | TGGATAGGTTTTAGAAGTGTGGTCATATTTTCGGGACCGGCGTTAGGTTTTATGAGCCTACAGA |
| | | TGTTATCCGGAGGTTTTTTATG |
| EcI07 | 21067 | AATTAGGTGTGCATGGCAATCATTAGAGCTCTTTTTAATTCTTTTTGTACTTGAGTTGTCCGGA |
| | | TGCAAGAAACGGTGTGAATTTATGGATCTTTGTGTAGGATGGGTTGACGCGAATAGAGGGACAG |
| | | TGTTCGATATCTGGTCGAAATTATTGGTATGTATCGTACCGTGGTTGCAATCAACGCTTAACAA |
| | | TGAGATCCGGAGGTGCAATATG |
| EcM10 | 21241 | TACGTAGGGAATTTGTATGAGGTTAGTGCATCTCGTGCCGTCTGAGTCGGAGAAGGGAAGAGCC |
| | | ATTGAACCTCCTTACGGATGGATACGCTTAATCCTTTGTATTGCCGTTGAGATATTGTTTTAGG |
| | | GTAGATTTTTGTTACTGGACACTGTGCTATGGTTCCTCGTTCGCGGGATGAATCCATGTACATT |
| | | ATAGCGCTGGAGGTTTTTTATG |
| EcM11 | 21312 | CTTTCTTGCGTTCCATGGGAAACGATATGTGTGTAGTAGCTGTTGTGGATTACACTTTTATATG |
| | | TCATCTTCATTAGTGTACGGTTTCCTCGGGTAAGATAGTGTGAACTCATGATAAGCTCGCTTGT |
| | | GGTATAAACTGGGGTTACTTCTTGAAGCTTTGAAGTGTTTAAGCGCGGTTTTTGCGTTTTTGTT |
| | | GGATTATAACCAGGAGGTACAGTATG |
| EcP05 | 21458 | TGATATTGTGTCCGTTCATCTACTCTCACTTTACACGTGATTCTTGTGATCAGCGTTTTCGTAA |
| | | TGAGGTGATCTTACGGGGTCCTGTTTTCTCTACTCTTCAATGAATATTTTACTTTCCAAGGTTC |
| | | GCCTTTTTTAGAGTTCTATTTTTTGTTATTGTATGCTGCGTCTAAATTTCGATTGTCGATCTAA |
| | | AAGGTAAGGAGTCACTCTATG |
| EcK07 | 21613 | GCATTAAGGTGTGCAGTCGCTGCTTCTGTACTTCCAAGTTGTCATCCATTGTTAGGTCTTGTTC |
| | | TGAACGGTGGTTACGCAAGACGGAATTTGTACGTTGCCGTATCCGTGTATGCTAAGGTGTTTTC |
| | | GCCTAGCTGGTGTTTAGGGTATGGCGTAAGTTCCCTTATTTCGTCATGAGCTGTTTAGGCACAA |
| | | TATGCAGGAGGGGTTATATG |
| EcE10 | 22181 | AAATTGTCTTGTTAATATTACGGTTGATTGTGTTCTTACTTCTCTTCTACACACTGTGTGGCCG |
| | | AATTATTTGTATGGTGTTCACCGTTACTCCTTTATGTTAGATTTGCTAAGGGCGTCTAGTAGGC |
| | | CTCTTCCGCTTTGTAGGATTGCTTAACACTATTTATTGGTGTATCAAAGTGTTTCAGAACGGTT |
| | | CTGCTAACGGAGGTTTCGAATG |
| EcN09 | 22454 | CATACAATTGATTCCGTCCCACCGGTTCTGTTACTTAGAATTGTTTACGAGAGGGTTCTGTGTG |
| | | CGTTTATTTTCAATATTTATAGTGCAACTCTGCTAGTGATTCGCGGAGCTGCGTTATG |
| EcF09 | 22633 | TTGAATCATTGATCGACGCTTTTGATCGGACGTGCTTATAGTCTTGATATGGTTGTAATCTTAT |
| | | TTATGCATAATGTTGCTCACATTGACTTTGGGATTCGGCTCCTTATATTAAGGTGACTAGGGAT |
| | | GTGCTAGCTTGAAAGGTGCAGGTCCCCCGATGCGCGTTTTTTGAGTGTCGTTTGCTATAGCAAG |
| | | AGGATGCGGAGTGCTGTTATG |
| EcF11 | 22645 | ATGTCAGTAACCTGCTTTTATGGAGTGTATGAATG |
| EcB10 | 22800 | ACTTGTGTTTACGATTGCTCAGGAGTTAATAGATG |
| EcG09 | 22977 | CTGACGCTACCGGACAGTGGGTCGGTAGGAGTTCAGTATGGTTACTTTTGTCGGTTGTCTGACA |
| | | TTACAAGTTTGGTTCCGTTATACGTGTTATAACTCGTTGTACTTTTCAAACGTGTATCGGCACA |
| | | AGGTAATGGAGGACCTACACATGGGGGCGCTTGTCGTCAATAGTAGTAGACTGTCTGCAGCAAC |
| | | AGAGTTCCGGAGGGTTATAATG |

Table 6: continued from the previous page.

| Clone | FIM[1] | DNA Sequence (5′→3′)[2] |
|---|---|---|
| EcG10 | 23426 | TAATCTTTGACGTCTCCGTCGCAACTAGCTCGGTCTTTCTCCTTCTCTGCATAAAACTCTTTCC<br>TTTGTACATTGTCGTGCACTTAACCTTTGCTAATCTGGGTAGTCGCGCCCTCCTGTTTAATATC<br>TTTAGAGCTATTATTTATTTTTCATTGTTGTGTTATCGTGTCATTCGATGATGAACGTAGATCG<br>TTGACTGAGGAGGTTGTTAATG |
| EcE07 | 23519 | CTCTAGTGGGATATCTCAGAATTCATATTGATTATCCAGCTGGAGTTTGCGTGTTTGTCGGATC<br>GTGATTTTTGGTATAGTTTCCTGTTAGAGTATCCTGTGTTAATTTGGGACATACCACTATCGCC<br>CTAGCACGGTGGTAGAAGTTGAAGTCGTGCTCTGTTAGCTCTGGTAGTGTTGGACATTGGATAT<br>CCACGTGGAGGTCGAAGATG |
| EcL09 | 23763 | GAGGTTTTGCCATGTGGTTTTAGGCGCGAGAGGGTGGCCGTATTTAGCCAAGTGATAGTTCTAA<br>CGGGATGCAGCCCTTCTTGCGGTAATTCGTGGTTGGTGGTATTGTATTGCTTGGCAAGGTTGTC<br>CTGGTAAAGGGTTGAGGTAGCGGACTATACTGTTTATTACTGTATGCGGGAGTTTTGCGTCGAT<br>CCGTGAGTGGAGTTGGGGTATG |
| EcC07 | 24397 | ATCAGGAAGTTCCTCTATGATGATAAGTTGTTTACTTGCGTGGATGTATTAGCTTTCATGTAGT<br>GGGTGAATTCGTTAGATGGCTGAGGTTTGACTTATAGATCAGGATTGTTGTTTTTTAAGGTCAA<br>GTGTCTTGACGTATACAGCCTGCCTGAATACTGTAATGCTATCTTAGGAATTAATTTTTGTAGC<br>ATAATGTAGGAGCTCGATCATG |
| EcJ09 | 24864 | GTGATTCACCAGTTGTTTTTAGCTTGCGTCTCTCTGGTTTCAGGTGTACTTATGTATCCGTGGT<br>TCTTGTAAGTTTTTGTAAATGCGTTCTTGCTTTTGTTATGAAAACGTAAATGATTGAGTATTTA<br>CCAGTACTGTTACAAGTGTCATACTTTGCTAGCACGTGTACTAAGGGGCGACCGGTACAACACG<br>CATTTCGGGAGATGTTAAATG |
| EcJ10 | 25316 | AATAAAGATTATAGTGTTTGTGCCTGTCAATCGATGTGTGGGGAGTTCGGAAATG |
| EcP01 | 25338 | GCCGTGAAATTTTACGTTGAGAAATTTCACCACGAAAGGATCAATTTGTCTTACTTGCGTCATA<br>ACAGGGGTTAAGTAATGATTTAGTCTAACTGTTTTCGTTAAAGTGTATGTAGTGAGCTCTCAGT<br>CGCGTGGGGAGAGCTTGGTCCTGTTGAATTGCCACGGCCTGTGGAAGTAGGGGGGTTTAATGTG<br>GGAGTCGAGGAGTAGTTTTATG |
| EcO12 | 25470 | TTTGGATTAAGCGCTAGCTGTTCATTTTACATCATTGGATGGGGGTAATACGGATACTTTCTGT<br>TTGAATTAAGACCACAGTACTATATGGGGCTGACTCAGGGTTACTTTTGTTGTTGCTGCTCAGA<br>CTAGCGTGGTTGGTTGTTTCACCTTGTGTTTTTTGAGTAGCGTTTATTTAAGGGCCTTTGTAAC<br>TTAATATAGGAGTCGAATTATG |
| EcK03 | 25770 | GACGGGAGGGATGTGGGTTGATTACCGGTATTTTTCGTTTCCATGACTCATGTATTGTCCTAAG<br>GGGCACGTCTACTGTATCCTGAGACTGTCTTCAATTTCTCATGGGAATTACGTTCGGGTGTTTC<br>CCTGTGGTGGGTGGAGAGGATCTCAGTGCGAAGTTGGTAAGTAATCTTTTGTAGAAAATACCCT<br>TCGTTCTTGGAGTATTGTTATG |
| EcP12 | 26046 | CGCAGGATTTTTCTTTTTGTATATTCTCCGGAATAGTCTATGGTATGCCGAGTTCTTGCTAATC<br>TCGGGGGCTAACGTTAATGGGTCTTGTTTAGATCTTTTATCGTGGGTAAGATCGAGTTGTACAG<br>GGGGGGCAACAATAATCAAGGCATTTTATACGATATGGACGTTGAGTAGGGATTCTCGTATCAG<br>TGAAGGGGAGATGTTACATG |
| EcO07 | 26371 | TGGTAGATTTGTTAGCTACTGTTTTTTGAATCGCTTGCTGTTTTTCTTAACTGTGGTCTCGATT<br>ATGTACCCAGTAGATTTTGCCTTGAATTGGTTTGTTTTTTATACTGTCTCTGTTTTCGTAAGTG<br>GTGTTGTGTATCGTGTGTTAACGGGATCCTCTTTTCGTGTTTAGTTAAGAGCGTTGGTCTAGTT<br>GTGGTTTAGGAGTTATAAAATG |
| EcL01 | 26844 | GTTGCCTAATGTTTATCTTAATCGTATATTGGTTTTTCCTATGACCATAGTTATAACATTCGAC<br>AAGTCCTCTTGTTGTGGCATCCGCTTATGGTAAATTCGTCAAATGGGTTATGAAACATTTGACT<br>AATATTAGTAACTGGATGTCCTCTATTGGTTTAAGGGTACTATTCTAGAGCTATTTGTAGGCAA<br>GGATAGTGGGAGTGAGTTCATG |

Table 6: continued from the previous page.

| **Clone** | **FIM**[1] | **DNA Sequence (5′→3′)**[2] |
|---|---|---|
| EcK12 | 26871 | CAGATTGAGTAATCTTTGTAGCTTATTCGTTTCCGACTAAATGATCTTTTAGTCAATCGTGTCA<br>TCCGGGGGCTGTAAAATTTGCAGTTTCGGATGTTCGTTTCGGTTATCATGGTTTGACGCAAAGG<br>AGTTATCATATG |
| EcA11 | 27507 | TGGTACATATATCTGAATCAGTGAGAGATTGTGCTTATATAAGTCCTGTATGGGTCTGAGCGCT<br>GAGAACTACGAATTAGGAGGGTTCTTATG |
| EcP07 | 28130 | TGGTAGATTTGTTAGCTACTGTTTTTTGAATCGCTTGCTGTTTTTCTTAACTGTGGTCTCGATT<br>ATGTACCCAGTAGATTTTGCCTTGAATTGGTTTGTTTTTTATACTGTCTCTGTTTTCGTAAGTG<br>GTGTTGTGTATCGTGTGTTAACGGGATCCTCTTTTCGTGTTTAGTTAAGAGCGTTGGTCTAGTT<br>GTGGTTTAGGAGTTATAAAATG |
| EcJ02 | 28163 | TTGTCTAGATGGCATACTTGAATAGTTTTTCTGATTACGTTTGTTAGACCGGTGTTCGGTAGTG<br>TAATGTGGAAGGATTCACTCTTAGGTTGGAGTCGATTGCTCGTTGGTTCTTCGTGCATAGGTTT<br>TGATAACCGTTTCCTGATCAGATGTTGCATCGTGGCCATGAGCTCGCGCTTTGTATTCGTTACT<br>GACAAATAGGAGGTAGTAGATG |
| EcC10 | 28530 | CTTATGTTTACTTGTGGTTGTACATAGCTGTTGTCGTTCCATTTTACGGTAGGGAGTTCGTCGT<br>CTTAACAAACAGTGTCTCTTTTTATGTTTCCGAGGAAGTTTATTTATTTATCTTTCTCACTTAC<br>ACTGTGAAATTCGAGGTGATGTTGCGATTGAAGCTCAAATGTCGTCGCTCATATGTGTCGGATT<br>TATTTGGAGGAGTGTTGAAATG |
| EcP08 | 28557 | TTTGCTAGAAGAGTTATGGTTCTGTGATCTTCGAGTGCGTCGATTTGTTCAAGTTACGTGTTGT<br>CCAGGTTGTGTGTACCCTTTATTCATTGTCTTCTTCATACGGGTATTTCTCGACTAAACTAGAT<br>TCTTTTTCACGAGATCGATTTCTTTCAACTTCACTGTTCGCTACAATGATATACTCATTGGGCG<br>TGATGAGGAGGGGTATAATG |
| EcD10 | 28586 | ATTACATTTTCGTTTGGTTTAGGCACGTGAATTTCTACAGAGGTTGGAGTGGGTATATGTAATG<br>CTTTAGATAGTATTGCATAGCGGCTGTAGCACTTCTTCGGCGTGACCTGTAGAGTTTGGAGTGG<br>GCGGGTCAATCAAAGTCAAACAGGGGTTGCCTTATCTGAGTTGTATCCCGCCAACCCGTCCATG<br>CAAATAGAGGAGTTATAAAATG |
| EcP03 | 28618 | TCTTTTATACGGGTGTGTCCTTTTTTTTCGTCGCTTCTTGTTACGGTGTCTTGATAAGTAGCTT<br>TGTTTTTGTTGTATGTATACAGGCTTCAATGGGAGTTCGTTAGAGGTTATGATATAGTGGTGTT<br>TAGGGTGGGCATTGTACTTGAGTAATTATGATGATTGGGTTAGCCCGATTCCATCTCTATTTTA<br>GCTTGTTTGGAGTGTTCATATG |
| EcH09 | 28696 | GTTTTGTGCTCCAGTGTAGTATCCGAGGTTAATGTATGTTATAGTTTTAGGGGGTTCTTTTTTC<br>TCGGAGGGTCTAAATG |
| EcO11 | 28745 | GGCTACTGTGGAGGTATATAATG |
| EcN07 | 28796 | GTGTATCCTGAAAGTCTGTTGCTGCGCCTGTGTACCTTCCACTCTCCGACTTATGGTCGGAGTC<br>GATATGTTGGGTGAAGATTTGCGAATGGTAACCTTTTTTTGTTGTAGCGTTGGGAAGTGTTGGT<br>GTGATTCGTTCGTTTTTAGACCTCTCTATGGCAAATCTTTGTTACACCTCTTCGGAGGCTTTTC<br>TTGAATTAGGAGTGATTAAATG |
| EcI08 | 29345 | GTTAGGAGGAGTTATATTATG |
| EcO05 | 29416 | TTGTGTGTTTGTTTGTTAGGTGTTTAATAACGAGGAGTGCCGGTATG |
| EcP11 | 29751 | ATGTTATGTGGCGTGTTCGAAGCGGATGTGACCCTTGCAGCTTTTCGTGTATGTGTTGGTCTGG<br>GCTCCGCTTCGTCTGGATTTTTTCCACTTTGTTGACTTATGAGCATTCAATATTGTGTAAGGTG<br>CGTTTTGTGGGAGTGTCAGGTGTGATACAGTAGGGTCGTAAAGGTGTTCTGTTGTAGGAAATCC<br>GCTAAAGGGAGGGTCGTTATG |

Table 6: continued from the previous page.

| Clone | FIM[1] | DNA Sequence (5′→3′)[2] |
|---|---|---|
| EcJ04 | 30254 | TTAATTTTCCAATGGGTTGCTTCATTTACATTATTTATCAACCCGTGGTAATGTCGTTCTACTT<br>TCATCTATAGTGTGTAATGTTATTGTTTTCTTTCTCTGTTGGTGGCTTTTTAGGGGGTTGTACT<br>GTACAGCGTTCTCTATTATGCGTTAAACTTTGAATTCATATAGGATGAATTTATACAAAAAGTC<br>ACATTTAAGGAGTGTTACGATG |
| EcJ05 | 33454 | GGATTTGCTATTTTTTTACCCGCTGATTTTCGTGTGAATCGCCTTGGTAGCTCTTTGGATCGCT<br>CACGACGGATAAAGAAATTTTGTGCGTTCTGTGTGTCTTGGGTTACCAGAAGTCGTCTTATCCT<br>TGTTGATCGGGCGAAGGGTGTGGGGCACGTTGCGGCGTCCGGAATGTGTATTAGTACCTGTGTA<br>ATGGTATAGGAGTTTTGTTATG |
| EcJ11 | 33792 | GGCATGCAAATATTTCGGCGCGGGCTATACGTTATCAGGAGCTTCGGCTCGTTGGCGTGGAATC<br>ACAGTGTACGTTATGCGATCGTAGTTTTTTAGGCGTTTGATCGGTCGGAGTTGCTTACAACAGG<br>TCCTAACCAGTCCTACAAATGGGTTTATCGGCATTTTAAGTAGTCTTGGAGGGCCTGGCGGTAT<br>CGGCTGATGGAGGTTAAATATG |
| EcL07 | 34130 | TTAGCTGCCAGATAACTGACAACTTCATTGTGCCGGAGGTTCACTCGGCCAATTAGGTATTGCG<br>TTTTGATCTTGTCGTCGTATTTCGAGTGTGCTGGCGTTTGCCTAGACATTAGGTGGTGGTAAGA<br>TGGGGGATATATCTGCGTGGGGTATCGGTAGTGATATCGAAATGACAAGCAGGGTGGAGGTTGA<br>TTATG |
| EcI12 | 35589 | GCTATAAGTATTGTTCCCTATTTGGTGTAACGCTTATAGTAGGGTTTATGGTTGAGGCCATCTC<br>TTAAGGGTAGGCTACTGATTCGTTGGTTGATTTGATAGCGCAGGCAGACATGTGTGAATATAGT<br>ACGCGGGTCGATTTACATGCGAACGCCTTCGGATTAGTAGGGCGAAGTGTCGTCAGTGGTGTTA<br>ACTTAATTGGAGTTGATTTATG |
| EcN03 | 38132 | ATGATACGGTCCATTGCAGACGGTTTATACGAGGTTTTATGATTTCATTTTGTTACTTGAGTAT<br>CCGAAATGCTCTACGGCCTTGGTCTTCTTTCAAGTTCGTCACTCTTGTTTGGGTGCGAATTTGG<br>TGATGGGCGTTAAATTGCATCGGTGTTGTTGAACGTTAATTGTTTGAGATCTTTCTTTTTTCAT<br>TAGGTAAGGGAGGGGATTAATG |
| EcJ08 | 38328 | GTTAGGAGGAGTTATATTATG |
| EcI03 | 38972 | CCTAGTTAGCGCGCCATTCATATCTGTGGCTTCTCACTGGCTGATCTTGAGGCGTGTGGTTGGT<br>ACAGTCGTTTGGCAGTGGTATTTTTGTATCACGGTAGATTGTGCGTGACCTATTGCGTATGATT<br>GTAGGTAGGAGCGGTCGTGCATAAGCCGCGTGTTCGCCATTCAGGCATTTGTTTAACTTGGGGA<br>GTGATTGGATG |
| EcI11 | 43241 | TCCTTTGTTAGCCGCATGCTAACTAGTACGGTGTGATTTTATCAGTTAAGATTGGGAGGTTGGG<br>TGATTTCGGGAGTTTGTGGGCCTCGCTGAAGTCGTTAGTTTTAGTTCTGTAACTCTAGGCATTA<br>GCCTTTTGAGGCGCCGACATGGGAGATTATTCATG |
| EcK04 | 44388 | AAATCCTTACCTCTTACCGTCGCATTTCATGTTTATATTCATTTTTGCGCGGTAGACTTATCGG<br>TTTGATTCGTGTATTTGTTCAACTTATAGGTCGTTACATGTGCATCTTGAGTGTGCTTGGAAGT<br>GTGTAGTGTGTCTGTTGTTGTAGTGTCTGGTCATGCGGTGTAATCGTGGGTGACTGCGGCGTCG<br>ATGCTCGTGGAGTTGCGACATG |
| EcM06 | 45741 | CCTGTGTTGTCACATAAAATGTTTAACTTGGCTTACTTGGGAGTATCCATTGCGGATCATACTA<br>CAACGGACAAATTATATAACTTGGTATTAATTTGGTGTAGTTTTTGACGGTTTAGTCTATGATC<br>TATGTTGAAGGTTCATCTAGTTTCGCCGCAGGGGCATGAGGTCATGTGGCGGTCGTCAAAGTAG<br>ATTGAAGGGAGTTGTGGAATG |
| EcP06 | 48422 | CTCGGAACTGTCGGATAGTGTATCACCATCGTGGCTTTGTTTTGATGAGGCGTCTGGAGTATAG<br>CGTTTTTTGTCTGTCGCTAACGTGCTTCGGTTTCAATGCGATGCCAATTTATTAGGTATCTTGT<br>CGACATTTATAGTAGAATCTTATGAGTAAGATCAATGTATCTTATTGTGTAATGGGGGCGGTTT<br>ACAGTTAAGGAGGTGTTTTATG |

Table 6: continued from the previous page.

| Clone | FIM[1] | DNA Sequence (5′→3′)[2] |
|---|---|---|
| EcK05 | 49336 | TTCTTCCGATGATCCTGTAGTTAGGTTATATATATCTCGGAATTCTGCTTTAAGGGGTGCGCGG GTCGTCACACTTTTTCTTCGTTTTCCTTGTGTCATCGTTTTATTTTTCGGGAGAATGGTAAGTA TACTGGAAAAGTTGGTCGTCAGTTAGTGTGGCTGCCTCTTCTTGTACTAGGGGCACTATATTGT GGTATGACGGAGTTAGATTATG |
| EcA12 | 50598 | TTAGGTGCGCGTTTTATGTTAATAGTGAACGTATACAGGTGTTTGTGTGCAGTTTCTGTTTTAG CGTATGCGATTGATCAGGTAAGTCATGTCTAGCGTTGAGGGATGGTGGCGCAGTACCGGCAACT CTGATGTGGATGTTGTCGCTAAGTTGAGTATGTATTATTATAGGTGAGTGACGGGTAAGTAGAG TGGGTAAAGGAGTGAGTGTATG |
| EcI04 | 51110 | TATGTTACATTTGCCTATCCGACGTTAAGTGATTGATTTTTGGGTTTAAATATATTTTATAACC GTTAGATCGTTTTAAGTATCTGATTTTCCGATTTCTGTTTAAGGATGATTTGATACCTGCCTCT ATGTTTCTTCCTATTTTTTGAGCGTGATTCGAATTAAAGGGCTATAATACTTTCTCTGTCGGTT CGCTGTTGGAGGGGTATAATG |
| EcK06 | 53717 | GCATAAATCAAGGGTACCTCGAGCTTAATAGATTCGCTTTTGCTTCGTCGATAGAGTTTTTATA GTATTAGTATTGTCCTATACACAGTTAATTGTTTGCGTGCTATAGTTGGTTTCTTGGTTCTGCA CCAAGTTTTTGTGTATAAACAGTAGGCCAAGTATCACGCTTTGTTTATCGAATGCTAGTTGTAA GATCGATAGGAGTCTGTATATG |
| EcO03 | 59961 | GCCAAATGATTCCTAATATGTCTCTTATCACGGGTCGCGCAACGGTTACTGTCGAGGTTCGCAT ATCGTTGCACCTGTGGGTTAAGAGCGGTACGTGTGAGAGTAGCACTTAATCTATTCCACCTTAG TACAGTGGGTATCACTTGGCTTTGTCGGGTTAGTTCGATCGTGTGAGTTAGATCGCTAAAGTTA GTGTGGTAGGAGTATATCTATG |
| EcJ12 | 61496 | ATTGGTCGGGCAATGACCCTTTTCGGTCAGCTTACGGACATAGGGCGTATGTTGTAGGGTACTT TGGTCCCAGCTTTATGTTATGATATCGTCCGTTGTACTCTAAACGGTTTGCATCTTGGCAACAT CTGCCTCTGGCCATAGCACGTTGAGTCCGTTAGAGTATGTCCCTGGAACAGTGTAATATGAGTA TGTTGGGGGAGGATGGGTATG |
| EcK09 | 84949 | TTGGGGTCTAAGTGTCGACTTGTGAGTTGAACTTTTTCTACTATTTTTAAGTTGGATTATCGAG AAGGGTTGCTAGGTGTTGTAGTATGTCCCGTGGGTATGTGTTAATGGTTTCCAGTGGAGGTTGA GTATG |

[1] Normalised fluorescent intensity measurement read outs.

[2] The ATG sequence at the 3′-end of each ArtPromU sequence is the start codon of the *mCherry* gene.

##### 1.4.5 Determination of transcription start sites.

To obtain RNA suitable for sequencing, all cell materials from 192 *E. coli* GeneEE clones were combined. With this pool, a 10 mL overnight pre-culture in LB was inoculated. From this culture, the main culture was inoculated with a starting $OD_{600}$ nm of 0.2 which was then grown until mid-log. For RNA isolation, samples of this culture were harvested by quick centrifugation, removal of the supernatant, flash-freezing in liquid nitrogen, and were stored at –80 °C. RNA was isolated and a transcription start site (TSS) library was prepared as described previously[18], with one modification: Instead of the unspecific loop adapter, the Illumina-TSS oligo (Table 1) was used for reverse transcription. Sequencing was done in a 75 nt run on Illumina MiSeq sequencing machine, resulting in 6,085,833 reads. Mapping of the reads was done using bowtie2[19] with standard parameters, using the ArtPromU sequences, each extended by the mCherry coding sequence between the ATG and the Illumina-TSS oligo, as references. From the 309,794 mapped reads (122,680 unique, 187,114 multiple), counting of read starts, identification of potential TSS and extraction of the region uptream of identified TSS was performed with a custom Perl script (see supplementary material, TSS-Perl). To determine the threshold for a potential TSS, a simple statistical analysis was performed by randomly distributing a number of hits in an array with the length of all reference sequences combined. The average number of reads per position as well as the standard deviation was calculated and the threshold number of reads for a "true" TSS was set to this average + 3 * standard deviation. Using this approach, a total number of 376 TSS was detected in 120 out of 157 ArtPromU sequences (Table 7).

Table 7: The list of all the motifs identified within the artificial promoters identified in *E. coli* with their TSS.

| Clone[1] | Pos.[2] | # Starts[3] | Shifted[4] | DNA Sequence Region, –50 to +1 (5′→3′)[5] |
|---|---|---|---|---|
| EcA01 | 98 | 95 | Yes | TTGATGAC**TTACAT**ACCAAAGTGCTTTCGCTTTTTG**CATTAT**ATGAGTCC |
| EcA01 | 93 | 15 | No | TAAGGTTGATGAC**TTACAT**ACCAAAGTGCTTTCGCTTTTTGCATTATATG |
| EcA02 | 121 | 17 | No | CTC**TTATTG**GTAACAATTTTACTATCGCTAAATGG**GCTCTT**TGAATGAAC |
| EcA02 | 117 | 29 | No | TGCTCTC**TTATTG**GTAACAATTTTACTATCGCTAAA**TGGGCT**CTTTGAAT |
| EcA02 | 84 | 23 | No | GTCCTCCATGAT**CTGTAT**AAATCATGCGATCTGTGC**TCTCTT**ATTGGTAA |
| EcA02 | 63 | 68 | No | TGGCGATG**TTGGGT**TTTCTATGTCCTCCATGATCTG**TATAAA**TCATGCGA |
| EcA04 | 196 | 188 | No | CCTGG**TCGACG**GTGGTGTCTTTGGTCGTAATGTGA**CAGTCT**ATTATGCAA |
| EcA04 | 194 | 15 | No | TACCTGG**TCGACG**GTGGTGTCTTTGGTCGTAATGT**GACAGT**CTATTATGC |
| EcA05 | 197 | 57 | Yes | GAGCTT**TTACAT**TCTTTAATTTGAGCAGCATGGTC**TACTCT**AGGTCAAAG |
| EcA05 | 194 | 52 | Yes | AAAGAGCTT**TTACAT**TCTTTAATTTGAGCAGCATGG**TCTACT**CTAGGTCA |
| EcA05 | 185 | 19 | No | CATGTACGGAAAGAGCTT**TTACAT**TCTTTAATTTGAG**CAGCAT**GGTCTAC |
| EcA05 | 168 | 55 | Yes | GCGAGTACG**TTATCG**GTCATGTACGGAAAGAGCTTT**TACATT**CTTTAATT |
| EcA06 | 71 | 55 | No | TTGTGAAGT**TTGTAT**ACCAGTTAGGCTTAGGACCA**TACAAT**TATGTACGA |
| EcA07 | 71 | 49 | No | TTGTGAAGT**TTGTAT**ACCAGTTAGGCTTAGGACCA**TACAAT**TATGTACGA |
| EcA10 | 203 | 63 | Yes | GACTCCATTG**CTATAT**AGGTGATGTCATGTCACGG**TGGGTT**AATTGGGAG |
| EcA10 | 178 | 22 | No | GTACCGGTAAGTGTTTCT**TTGGCC**AGACTCCATTGC**TATATA**GGTGATGT |
| EcA10 | 132 | 739 | No | AGACAATT**TTGCTT**GTTGTTGGGCCACTTCGAAGTT**TATCTT**ATATGTAC |
| EcA11 | 48 | 50 | No | GTGGTACATATAT**CTGAAT**CAGTGAGAGATTGTGCT**TATATA**AGTCCTGT |
| EcA12 | 192 | 25 | No | TCGCTAAG**TTGAGT**ATGTATTATTATAGGTGAGTGACGGGTAAGTAGAGT |
| EcA12 | 189 | 19 | No | TTGTCGCTAAG**TTGAGT**ATGTATTATTATAGGTGAG**TGACGG**GTAAGTAG |
| EcA12 | 184 | 24 | No | GGATG**TTGTCG**CTAAGTTGAGTATGTATTATTATAGGTGAGTGACGGGTA |
| EcA12 | 178 | 3320 | Yes | TGATGTGGATG**TTGTCG**CTAAGTTGAGTATGTAT**TATTAT**AGGTGAGTGA |
| EcA12 | 176 | 7647 | No | TCTGATGTGGATG**TTGTCG**CTAAGTTGAGTATGTAT**TATTAT**AGGTGAGT |
| EcA12 | 174 | 199 | No | ACTCTGATGTGGATG**TTGTCG**CTAAGTTGAGTATG**TATTAT**TATAGGTGA |
| EcA12 | 171 | 38 | Yes | GCAACTCTGATG**TGGATG**TTGTCGCTAAGTTGAGTA**TGTATT**ATTATAGG |
| EcB01 | 165 | 31 | No | TTGGTG**TAGTCG**TAAATTATTTTTAGATTCTCTCAAA**TGACTT**TTCGCAT |
| EcB01 | 146 | 17 | No | GTAGTTAGCTAG**TTAGTG**TTTGGTGTAGTCGTAAAT**TATTTT**TAGATTCT |
| EcB01 | 115 | 122 | No | GGACGTAG**TCGGGT**GCTTTTGCTTAGATATTGTAGT**TAGCTA**GTTAGTGT |
| EcB01 | 104 | 16 | No | GTTCTGTTCT**TGGACG**TAGTCGGGTGCTTTTGCTTA**GATATT**GTAGTTAG |
| EcB01 | 101 | 307 | No | AGAGTTCTGTTCT**TGGACG**TAGTCGGGTGCTTTTGCT**TAGATA**TTGTAGT |
| EcB04 | 203 | 75 | Yes | GGTTTCCTC**TTATTT**TGTCAAGCTTGATAACCCAGG**TGTACT**GGTTGGAG |
| EcB04 | 192 | 458 | No | GTACTCTGTA**TGGTTT**CCTCTTATTTTGTCAAGCTT**GATAAC**CCAGGTGT |
| EcB04 | 190 | 49 | No | TTGTACT**CTGTAT**GGTTTCCTCTTATTTTGTCAAGCT**TGATAA**CCCAGGT |
| EcB04 | 180 | 26 | No | TTGTGCTTTG**TTGTAC**TCTGTATGGTTTCCTCT**TATTTT**GTCAAGCTTGA |
| EcB04 | 178 | 81 | No | AATTGTGCT**TTGTTG**TACTCTGTATGGTTTCCTCT**TATTTT**GTCAAGCTT |
| EcB04 | 171 | 13 | No | AACCGAAAA**TTGTGC**TTTGTTGTACTCTGTATGGTT**TCCTCT**TATTTTGT |
| EcB04 | 140 | 66 | Yes | TTCTTTAATACGTTCCCT**TTGGCT**CGGGTTGAACC**GAAAAT**TGTGCTTTG |
| EcB07 | 135 | 83 | Yes | TAGTCG**TCGACT**ATACTATGGAGATTCATGCATTA**CAGTAT**CGCTAGAGT |
| EcB07 | 110 | 21 | No | CTGC**TTGACT**TGGTATCTCCGCCCTTAGTCGTCGAC**TATACT**ATGGAGAT |
| EcB07 | 22 | 27 | No | TCAGCGAAT**TCGCGG**CCGCTTCTAGAGTGAGTCATC**TATACT**ACCCATGG |

Table 7: continued from the previous page.

| Clone[1] | Pos.[2] | # Starts[3] | Shifted[4] | DNA Sequence Region, –50 to +1 (5′→3′)[5] |
|---|---|---|---|---|
| EcB08 | 61 | 221 | No | CAACGTTAGGTTTT**TTAGTT**CCTAGTGTTGAGTGT**TATGCT**TTTTCATAG |
| EcB09 | 14 | 16 | No | CAACGGCG**TCAGCG**AATTCGCGGCCGCTTCTAGA**GATTCT**GTATCTTGAG |
| EcB11 | 167 | 85 | No | GTTATCGCC**TTGTGT**GGGCGTGTATGACCGAGCGAA**TGTGGT**CCGTAAGC |
| EcB11 | 165 | 56 | No | TGGTTA**TCGCCT**TGTGTGGGCGTGTATGACCGAGC**GAATGT**GGTCCGTAA |
| EcB11 | 163 | 32 | No | GCTGGTTA**TCGCCT**TGTGTGGGCGTGTATGACCGAGC**GAATGT**GGTCCGT |
| EcB11 | 101 | 24 | Yes | TTCGTACGTTG**TCGCCC**TCAACGCATTATATTTAGTC**TATCTG**TAAGGGA |
| EcB11 | 97 | 18 | No | TCATTTCGTACG**TTGTCG**CCCTCAACGCATTATATT**TAGTCT**ATCTGTAA |
| EcB11 | 91 | 21 | No | GTGCATTCATT**TCGTAC**GTTGTCGCCCTCAACGCAT**TATATT**TAGTCTAT |
| EcC02 | 203 | 1408 | Yes | CTTCCAACGGTATT**TTGTAT**TTACTCTATTCGGCTA**TAACAT**AGTGGGAG |
| EcC02 | 200 | 221 | Yes | TCTCTT**CCAACG**GTATTTTGTATTTACTCTATTCGGC**TATAAC**ATAGTGG |
| EcC02 | 197 | 56 | Yes | TTTTCTCTT**CCAACG**GTATTTTGTATTTACTCTAT**TCGGCT**ATAACATAG |
| EcC02 | 195 | 20 | No | GGTTTTCTCTT**CCAACG**GTATTTTGTATTTACTCTAT**TCGGCT**ATAACAT |
| EcC02 | 190 | 30 | No | TGTCTGGTTTTCTCTT**CCAACG**GTATTTTGTATT**TACTCT**ATTCGGCTAT |
| EcC02 | 184 | 21 | No | CATTA**TTGTCT**GGTTTTCTCTTCCAACGGTATTTTG**TATTTA**CTCTATTC |
| EcC02 | 182 | 13 | No | GACATTA**TTGTCT**GGTTTTCTCTTCCAACGGTATTT**TGTATT**TACTCTAT |
| EcC02 | 177 | 67 | No | GGTGTGACATTA**TTGTCT**GGTTTTCTCTTCCAACGG**TATTTT**GTATTTAC |
| EcC02 | 148 | 124 | No | CAGGTTGAATG**TTGTGG**TTCTAATGATTGGGTGTGA**CATTAT**TGTCTGGT |
| EcC02 | 146 | 123 | No | TGCAGG**TTGAAT**GTTGTGGTTCTAATGATTGGGTGT**GACATT**ATTGTCTG |
| EcC02 | 90 | 1581 | Yes | TTTGCGGGAGATG**TTGCTT**TATTTTGGGATTCTTGC**CACACT**TTTTTTCT |
| EcC04 | 204 | 54 | No | AGATGT**TTGGGT**TTTAGGATTTTTCATTCAGTGCAACT**TGTAGT**GGGAGA |
| EcC04 | 192 | 42 | Yes | AGAGGATC**TGGCAG**ATGTTTGGGTTTTAGGATTTTT**CATTCA**GTGCAACT |
| EcC04 | 174 | 24 | Yes | TTCGTCGACAT**TCATGG**GAGAGGATCTGGCAGATGTT**TGGGTT**TTAGGAT |
| EcC04 | 144 | 21 | No | TTAGTTGTTAAC**TTGCAC**CTTGTTTGACTTTTCGTC**GACATT**CATGGGAG |
| EcC04 | 117 | 204 | No | ATAGGTTT**TTGTTT**CAAGTTTTTCTCTTTAGTTGT**TAACTT**GCACCTTGT |
| EcC04 | 115 | 33 | No | GGATAGGTTT**TTGTTT**CAAGTTTTTCTCTTTAGTTGT**TAACTT**GCACCTT |
| EcC04 | 112 | 62 | Yes | TCTGGATAGGTTT**TTGTTT**CAAGTTTTTCTCTTTAGT**TGTTAA**CTTGCAC |
| EcC05 | 190 | 65 | No | TCTATAGGGGGCATCCT**TTGGGT**TTGTAATTAGTCG**TATAGT**CATCAAGT |
| EcC05 | 51 | 212 | No | TTTGACAGTTT**TTGTAT**GAGATGAATGGTGGAGTTA**TACTCT**AATGCTGT |
| EcC07 | 204 | 43 | No | ATACTGTAATG**CTATCT**TAGGAATTAATTTTTGTAG**CATAAT**GTAGGAGC |
| EcC07 | 201 | 53 | Yes | TGAATACTGTAATG**CTATCT**TAGGAATTAATTTTTG**TAGCAT**AATGTAGG |
| EcC07 | 193 | 20 | No | AGCCTGC**CTGAAT**ACTGTAATGCTATCTTAGGAAT**TAATTT**TTGTAGCAT |
| EcC07 | 189 | 17 | No | ATACAGC**CTGCCT**GAATACTGTAATGCTATCTTAG**GAATTA**ATTTTTGTA |
| EcC07 | 187 | 15 | No | GTATACAGC**CTGCCT**GAATACTGTAATGCTATCT**TAGGAA**TTAATTTTG |
| EcC07 | 184 | 66 | Yes | GACGTATACAGC**CTGCCT**GAATACTGTAATGCTATCT**TAGGAA**TTAATTT |
| EcC07 | 174 | 21 | No | CAAGTGTC**TTGACG**TATACAGCCTGCCTGAATACTG**TAATGC**TATCTTAG |
| EcC07 | 169 | 1667 | No | AAGGTCAAGTGTC**TTGACG**TATACAGCCTGCCTGAA**TACTGT**AATGCTAT |
| EcC07 | 65 | 64 | Yes | ATGATGATAAG**TTGTTT**ACTTGCGTGGATGTAT**TAGCTT**TCATGTAGTGG |
| EcC07 | 29 | 44 | Yes | ATTCGCGG**CCGCTT**CTAGAGATCAGGAAGTTCCTC**TATGAT**GATAAGTTG |
| EcC08 | 194 | 18 | No | TTGCTTGG**TTGTAT**TCTGGTGAGTATGAGTTTCTTG**TATATC**TGAATCGA |
| EcC08 | 170 | 15 | No | AGTGTG**TTGAGG**GAGGTTGTTACATTGCTTGGTTG**TATTCT**GGTGAGTAT |
| EcC08 | 133 | 23 | Yes | GTTTGGATATA**TTATTG**AGATTGGATAGAGTTCCGT**TAGTGT**GTTGAGGG |
| EcC08 | 122 | 82 | Yes | TTGTGTGTAACGT**TTGGAT**ATATTATTGAGATTGGA**TAGAGT**TCCGTTAG |
| EcC08 | 115 | 39 | No | ATCGTTC**TTGTGT**GTAACGTTTGGATATATTATT**GAGATT**GGATAGAGTT |
| EcC08 | 109 | 37 | No | GTGTGAATCGTTC**TTGTGT**GTAACGTTTGGATATAT**TATTGA**GATTGGAT |

Table 7: continued from the previous page.

| Clone[1] | Pos.[2] | # Starts[3] | Shifted[4] | DNA Sequence Region, –50 to +1 (5′→3′)[5] |
|---|---|---|---|---|
| EcC08 | 106 | 120 | Yes | TTTGTGTGAA**TCGTTC**TTGTGTGTAACGTTTGGATA**TATTAT**TGAGATTG |
| EcC09 | 196 | 42 | Yes | GTCATTGG**TGACTT**TTGCTATAGTGGTTAACCGTAAA**CGAAGT**AGGTCAA |
| EcC09 | 180 | 15242 | Yes | TTCCTATTTTGT**TTGAGT**CATTGGTGACTTTTGC**TATAGT**GGTTAACCGT |
| EcC09 | 177 | 3081 | Yes | GTTTTCCTATT**TTGTTT**GAGTCATTGGTGACTTTTGC**TATAGT**GGTTAAC |
| EcC09 | 174 | 34 | Yes | TGAGTTTTCCTATT**TTGTTT**GAGTCATTGGTGACTTT**TGCTAT**AGTGGTT |
| EcC09 | 101 | 59 | No | TGTGTTGGATT**TCGACT**CTTGCTCTTCTTTGCTG**TACGTT**TCTCTTCTGT |
| EcC10 | 139 | 1306 | Yes | TTTCCGAGGAAGT**TTATTT**ATTTATCTTTCTCACT**TACACT**GTGAAATTC |
| EcC10 | 137 | 176 | No | TGTTT**CCGAGG**AAGTTTATTTATTTATCTTTCTCACT**TACACT**GTGAAAT |
| EcC11 | 165 | 163 | Yes | TGGTTA**TCGCCT**TGTGTGGGCGTGTATGACCGAGC**GAATGT**GGTCCGTAA |
| EcC11 | 163 | 25 | No | GCTGGTTA**TCGCCT**TGTGTGGGCGTGTATGACCGAGC**GAATGT**GGTCCGT |
| EcC11 | 102 | 21 | No | TCGTACGTTG**TCGCCC**TCAACGCATTATATTTAGTC**TATCTG**TAAGGGAG |
| EcC11 | 97 | 25 | No | TCATTTCGTACG**TTGTCG**CCCTCAACGCATTATATT**TAGTCT**ATCTGTAA |
| EcC11 | 91 | 19 | No | GTGCATTCATT**TCGTAC**GTTGTCGCCCTCAACGCAT**TATATT**TAGTCTAT |
| EcD01 | 191 | 408 | No | TAATTTT**TTGTCT**GTTCGACAATGTGATAAGAGGA**TAATGT**GAATAGTGG |
| EcD01 | 188 | 86 | Yes | GCTTAATTTT**TTGTCT**GTTCGACAATGTGATAAGAG**GATAAT**GTGAATAG |
| EcD01 | 182 | 15 | No | CGCCGGGCTTAATTTT**TTGTCT**GTTCGACAATGTGA**TAAGAG**GATAATGT |
| EcD04 | 197 | 80 | No | GTTTTTTATTT**TCGAGT**GCTGAGGCGTCAAGTGTGG**CATTTT**TGGGTTGA |
| EcD04 | 194 | 25 | Yes | TGTGTTTTTTATTT**TCGAGT**GCTGAGGCGTCAAGTG**TGGCAT**TTTTGGGT |
| EcD04 | 190 | 26 | No | TTTATGTGTTTT**TTATTT**TCGAGTGCTGAGGCGTC**AAGTGT**GGCATTTTT |
| EcD04 | 103 | 26 | No | CTCATCATCT**TTGCGG**CCTTCACAATGAAGTACTTGT**TATATT**GAGTTGT |
| EcD04 | 58 | 161 | No | GTTCGTCAGATG**TTGTAT**AATGTTGAATGTATTTT**TATACT**GTTACTCAT |
| EcD04 | 55 | 15 | No | TTGGTTCGTCAGATG**TTGTAT**AATGTTGAATGTATTTT**TATACT**GTTACT |
| EcD05 | 91 | 132 | No | TTCTGAC**TCGTAT**CTTCAAGGTAACGGTTTTTAGAC**TATCCT**TAGAGGGT |
| EcD06 | 162 | 37 | No | AGACAACACGCCTTT**TCGGTT**TGGGTAGTTATTTT**TACAGT**AATTGGTCT |
| EcD07 | 127 | 67 | No | GCATAGCGG**CTGTAG**CACTTCTTCGGCGTGACCTG**TAGAGT**TTGGAGTGG |
| EcD08 | 207 | 1769 | Yes | TCTTTTTA**TTATTC**TGGTATCAAGGCAATGAGTGC**TATAAT**TGGAGTCTG |
| EcD08 | 177 | 109 | No | TTCACCGCATTTC**TGGACC**TTAATTGTTGTTCTTTT**TATTAT**TCTGGTAT |
| EcD08 | 78 | 961 | Yes | TTTCTCGTT**TTGTTT**TCACTTCTGTTTCATTTCCGT**TATTCT**TCTTACGG |
| EcD09 | 188 | 14 | No | ATGTCTAGG**CCAGTT**CTAGAGGAATGGCGTTCTCG**CCTCCT**CATAGAAAC |
| EcD09 | 68 | 226 | No | CCTGGTTTAT**TTGACT**TGTTTTTGGTGTTTGTAG**TAAATT**ATGTCTGCGA |
| EcD09 | 66 | 1869 | No | ATCCTGGTTTAT**TTGACT**TGTTTTTGGTGTTTGTAG**TAAATT**ATGTCTGC |
| EcE01 | 137 | 116 | Yes | TACGTAATCCGG**TTGGCG**GCCTTATTCCGGCGAGTAG**TCGGCT**GCTACGT |
| EcE01 | 114 | 32 | Yes | ACCTTGGATGT**TTGGGT**CTATTCTACGTAATCCGGT**TGGCGG**CCTTATTC |
| EcE01 | 101 | 26 | No | TAGCTTCAGTT**TTACCT**TGGATGTTTGGGTCTATTC**TACGTA**ATCCGGTT |
| EcE01 | 95 | 154 | Yes | TTCTGGTAGCT**TCAGTT**TTACCTTGGATGTTTGGGTC**TATTCT**ACGTAAT |
| EcE01 | 31 | 2096 | Yes | TCGCGG**CCGCTT**CTAGAGGCGAGCTTTTTATAGTTG**TAAACT**AAACATCT |
| EcE03 | 164 | 31 | Yes | CGAAGTCAAC**CTGGCC**ATAATTCGGTTGGAGTGTGT**TAACCG**CGGTTAAT |
| EcE03 | 155 | 43 | Yes | TGTTCGTTACGAAG**TCAACC**TGGCCATAATTCGGT**TGGAGT**GTGTTAACC |
| EcE03 | 117 | 2640 | Yes | AGTTGAA**TGGCGC**TTTCTCCTGTGTGTTATCTGTGC**TATGTT**CGTTACGA |
| EcE04 | 204 | 1493 | Yes | GGCCACTT**TTGTAC**TGTTATTATTTCGAAATACTTG**TACTCT**GAAGGAGA |
| EcE04 | 202 | 14 | No | TCGGCCACTT**TTGTAC**TGTTATTATTTCGAAATACT**TGTACT**CTGAAGGA |
| EcE04 | 188 | 13 | No | TCAATGTTCGGTTC**TCGGCC**ACTTTTGTACTGTTAT**TATTTC**GAAATACT |
| EcE04 | 185 | 53 | No | CTTTCAATGT**TCGGTT**CTCGGCCACTTTTGTACTGT**TATTAT**TTCGAAAT |
| EcE04 | 179 | 120 | Yes | TTGCCCCTT**TCAATG**TTCGGTTCTCGGCCACTTTTG**TACTGT**TATTATTT |

Table 7: continued from the previous page.

| Clone[1] | Pos.[2] | # Starts[3] | Shifted[4] | DNA Sequence Region, –50 to +1 (5′→3′)[5] |
|---|---|---|---|---|
| EcE04 | 176 | 34 | Yes | AGGTTGCCCCTT**TCAATG**TTCGGTTCTCGGCCACTTT**TGTACT**GTTATTA |
| EcE04 | 56 | 60 | No | CGACGGCCTGTT**TGGACT**CTCATTTGTTGAGTGTGA**TATGTT**TCCGTTAT |
| EcE06 | 88 | 40 | No | TTTG**TCAGAG**AGTACAGTATTAAGCGATTTTATGG**TGTCCT**AGGAGGTAG |
| EcE07 | 189 | 13 | No | TAGAAGTTGAAG**TCGTGC**TCTGTTAGCTCTGGTAGTGT**TGGACA**TTGGAT |
| EcE07 | 186 | 66 | Yes | TGGTAGAAG**TTGAAG**TCGTGCTCTGTTAGCTCTGG**TAGTGT**TGGACATTG |
| EcE07 | 177 | 15 | No | CTAGCACGG**TGGTAG**AAGTTGAAGTCGTGCTCTGT**TAGCTC**TGGTAGTGT |
| EcE07 | 169 | 13 | No | CTA**TCGCCC**TAGCACGGTGGTAGAAGTTGAAGTCG**TGCTCT**GTTAGCTCT |
| EcE07 | 117 | 3227 | No | TTTTTGGTA**TAGTTT**CCTGTTAGAGTATCCTGTGT**TAATTT**GGGACATAC |
| EcE07 | 115 | 1101 | No | GATTTT**TGGTAT**AGTTTCCTGTTAGAGTATCCTGTGT**TAATTT**GGGACAT |
| EcE07 | 102 | 826 | Yes | TTTG**TCGGAT**CGTGATTTTTGGTATAGTTTCCTGT**TAGAGT**ATCCTGTGT |
| EcE07 | 100 | 56 | No | TGTTTG**TCGGAT**CGTGATTTTTGGTATAGTTTCCTGT**TAGAGT**ATCCTGT |
| EcE07 | 90 | 1117 | Yes | GGAGTTTGCGTGT**TTGTCG**GATCGTGATTTTTGGTA**TAGTTT**CCTGTTAG |
| EcE07 | 87 | 30 | Yes | GCTGGAGT**TTGCGT**GTTTGTCGGATCGTGATTTTTGG**TATAGT**TTCCTGT |
| EcE09 | 195 | 61 | Yes | TTATCGGCGGCGAG**CTGCAC**GGGCGCTTGGGGGTCC**CGAACT**CATTGGCT |
| EcE09 | 149 | 44 | Yes | CCGTTGCGTGCTATCC**TTGAGT**TTATGAAAGGCGGGT**TACTTA**CGCTTAT |
| EcE09 | 127 | 52 | No | CATTTACTGTC**TTGCTT**ACCCGCCGTTGCGTGC**TATCCT**TGAGTTTATGA |
| EcE09 | 125 | 3126 | No | TTCATTTACTGTC**TTGCTT**ACCCGCCGTTGCGTGC**TATCCT**TGAGTTTAT |
| EcE10 | 191 | 26 | No | AGGA**TTGCTT**AACACTATTTATTGGTGTATCAAAG**TGTTTC**AGAACGGTT |
| EcE10 | 182 | 1010 | Yes | CCGCTTTGTAGGA**TTGCTT**AACACTATTTATTGGTG**TATCAA**AGTGTTTC |
| EcE10 | 113 | 469 | Yes | AATTAT**TTGTAT**GGTGTTCACCGTTACTCCTTTATGT**TAGATT**TGCTAAG |
| EcE10 | 109 | 26 | No | GCCGAATTAT**TTGTAT**GGTGTTCACCGTTACTCCTT**TATGTT**AGATTTGC |
| EcE12 | 158 | 164 | No | CTTTTTTAT**TCAGTG**ACTTCACGTGACACAGTACTT**TAGAGT**TGAGGTGA |
| EcE12 | 153 | 20 | No | CTACGCTTTT**TTATTC**AGTGACTTCACGTGACACAG**TACTTT**AGAGTTGA |
| EcF01 | 202 | 88 | Yes | CATCGCAACA**TTATTT**GGTCACATGTGAACATGTGT**TAGGTA**TGTAGGGA |
| EcF01 | 171 | 70 | No | TTCGGCTACGCC**TTGAAT**CCTGTTCAGTAGCCATCG**CAACAT**TATTTGGT |
| EcF01 | 166 | 20 | No | TTGGAT**TCGGCT**ACGCCTTGAATCCTGTTCAGTAGC**CATCGC**AACATTAT |
| EcF01 | 111 | 59 | No | TATACATGG**CCGCTG**GACTGGGAGTCAAAATGCTCA**GAGTAT**GTTCGTTC |
| EcF02 | 197 | 52 | Yes | TGTTCTTTGT**TTGGGT**ATATGGGTGCTCAGTGTG**CAAATT**GGGCCAATTG |
| EcF02 | 166 | 127 | Yes | AGATA**TGGACT**TGTGAGCCTAATGTATAGAGTGTTCTTTGTTTGGGTATA |
| EcF02 | 150 | 20 | No | TTGTAGTCTA**TGATGT**AGATATGGACTTGTGAGCC**TAATGT**ATAGAGTGT |
| EcF02 | 122 | 421 | No | TTAGGCTCAGGGTT**TTGGAT**ATAATAGCTTGTAGTC**TATGAT**GTAGATAT |
| EcF02 | 106 | 656 | Yes | TCAATGTTTTT**TTGTTT**TAGGCTCAGGGTTTTGGA**TATAAT**AGCTTGTAG |
| EcF02 | 104 | 1097 | No | TTTCAATGTTTTT**TTGTTT**TAGGCTCAGGGTTTTGGA**TATAAT**AGCTTGT |
| EcF03 | 152 | 15 | No | TGCATCTCTC**TGGACG**AACTTGTCGGGCTATTGTA**GGTTTT**TTTCCGTGT |
| EcF03 | 150 | 36 | No | ATTGCATCTCTC**TGGACG**AACTTGTCGGGCTATTG**TAGGTT**TTTTTCCGT |
| EcF03 | 113 | 23 | Yes | TTC**TTGACC**AAGTGGGTCTATACTTTTCGTTGTAA**TAATTG**CATCTCTCT |
| EcF03 | 96 | 1893 | No | CTTGCCATGTTT**TTGCCT**TCTTGACCAAGTGGGTC**TATACT**TTTCGTTGT |
| EcF03 | 93 | 17 | No | CCTCTTGCCATGTTT**TTGCCT**TCTTGACCAAGTGGGTC**TATACT**TTTCGT |
| EcF04 | 178 | 404 | No | CCCGCAGATGA**TTGTCT**GGAACTGCCTTTTGGGT**TACTTC**TAGGTGTTAG |
| EcF04 | 175 | 71 | Yes | TATCCCGCAGATGA**TTGTCT**GGAACTGCCTTTTGGGT**TACTTC**TAGGTGT |
| EcF07 | 189 | 31 | Yes | ACAATTTAAT**CCGCCT**GACGCTTACGGTTTCCTAAG**CATTCA**GGATATGT |
| EcF08 | 155 | 53 | No | GATTTGCT**TCAACT**GTACTTTAATTGTGGTAATGG**TATTTT**TTAACAAAC |
| EcF08 | 152 | 37 | No | TCGGATTTGCT**TCAACT**GTACTTTAATTGTGGTAA**TGGTAT**TTTTTAACA |
| EcF08 | 150 | 14 | No | GTTCGGAT**TTGCTT**CAACTGTACTTTAATTGTGGTAA**TGGTAT**TTTTTAA |

Table 7: continued from the previous page.

| Clone[1] | Pos.[2] | # Starts[3] | Shifted[4] | DNA Sequence Region, –50 to +1 (5′→3′)[5] |
|---|---|---|---|---|
| EcF08 | 88 | 81 | No | TGTTTCACAG**TTGCCT**ATATTTAGTACATTTTTAT**TATAAT**TAAGTATGG |
| EcF08 | 86 | 185 | No | TTTGTTTCACAG**TTGCCT**ATATTTAGTACATTTTTAT**TATAAT**TAAGTAT |
| EcF09 | 146 | 26 | No | TTCGGCTCC**TTATAT**TAAGGTGACTAGGGATGTGC**TAGCTT**GAAAGGTGC |
| EcF09 | 123 | 209 | No | TGTTGCTCACA**TTGACT**TTGGGATTCGGCTCCTTA**TATTAA**GGTGACTAG |
| EcF09 | 120 | 290 | Yes | TAATGTTGCTCACA**TTGACT**TTGGGATTCGGCTCCT**TATATT**AAGGTGAC |
| EcF10 | 198 | 13 | No | ATTTC**TTGTGC**ATGGGTGCAGACGCCTGGGTTAACAGC**TATGTA**AAAGAC |
| EcF10 | 122 | 89 | No | TGGGGCTG**TGGGAG**TTTTTATCAGTGGGAATGTAC**TAGACT**CGGTGATGT |
| EcF11 | 11 | 21 | No | ATCCAACGGCG**TCAGCG**AATTCGCGGCCGCTTCTA**GAGATG**TCAGTAACC |
| EcF12 | 176 | 13 | No | CTCGTGTGTGTTA**TTAAGT**TTACGTGTTGGCGTGC**TATCTT**CCTCATCAG |
| EcF12 | 153 | 14 | No | ATTTCT**TCGAAG**TAGTGTGCTTTCTCGTGTGTGTTAT**TAAGTT**TACGTGT |
| EcF12 | 56 | 168 | No | CCAAAGGTC**TTGATG**TATTCATTACAATTTTTAGG**TATTCT**TGGTGACGG |
| EcG01 | 204 | 6603 | Yes | ATCTCGTCAAGGATG**TTGCTT**GGTCCATGGATGAGT**TATTAT**AGGAGTGT |
| EcG01 | 195 | 37 | Yes | TTAGATTTAATCTCG**TCAAGG**ATGTTGCTTGGTC**CATGGA**TGAGTTATTA |
| EcG01 | 166 | 16 | No | TATCCG**TGGGAC**GGGTTAGTGCGTATTATTTAGATT**TAATCT**CGTCAAGG |
| EcG01 | 161 | 23 | No | TCTTTTAT**CCGTGG**GACGGGTTAGTGCGTATTATT**TAGATT**TAATCTCGT |
| EcG01 | 158 | 30 | No | GAATCTTTTAT**CCGTGG**GACGGGTTAGTGCGTATTATT**TAGATT**TAATCT |
| EcG02 | 195 | 24 | No | TGAATATGGTTTC**TCGGTT**TATGTATGGCGTCCTGAG**TGGGGC**TTTTACT |
| EcG02 | 125 | 81 | Yes | GTTGGGAACGGCT**TTGCGG**TTATAGGGTGTTGATA**TACTCT**AATCGGTTA |
| EcG02 | 122 | 65 | Yes | TTTGTTGGGAACGGCT**TTGCGG**TTATAGGGTGTTGA**TATACT**CTAATCGG |
| EcG07 | 171 | 405 | Yes | AATGCTAGAGTTT**CTGTTC**AGTATTGGTTAGTTGG**TATAAT**CCAAGGTTG |
| EcG07 | 126 | 2051 | Yes | TGGAT**TTGTAT**CCTTCCGGTGTTTGGCTAATGTTAG**TATGAT**GTGAATGC |
| EcG07 | 122 | 71 | Yes | AACCTGGAT**TTGTAT**CCTTCCGGTGTTTGGCTAATGT**TAGTAT**GATGTGA |
| EcG07 | 117 | 23 | No | ATTCTAACCTGGAT**TTGTAT**CCTTCCGGTGTTTGGC**TAATGT**TAGTATGA |
| EcG08 | 198 | 57 | Yes | AAGTCACTGTGG**TTGTGG**TGGCGAGGAGAACGTATG**TCGATT**TTTTAAGT |
| EcG08 | 145 | 13 | No | TTGTTGGT**TTGTGC**TCTGTAGTATATGGTTCGGTAAA**TAGACC**GCACGGT |
| EcG08 | 135 | 36 | No | GCGGGAGCAC**TTGTTG**GTTTGTGCTCTGTAGTA**TATGGT**TCGGTAAATAG |
| EcG08 | 133 | 195 | No | AAGCGGGAGCAC**TTGTTG**GTTTGTGCTCTGTAGTA**TATGGT**TCGGTAAAT |
| EcG08 | 131 | 35 | No | GCAAGCGGGAGCAC**TTGTTG**GTTTGTGCTCTGTAGTA**TATGGT**TCGGTAA |
| EcG10 | 178 | 17 | No | TTAGAGCTA**TTATTT**ATTTTTCATTGTTGTGTTATCG**TGTCAT**TCGATGA |
| EcG10 | 175 | 1480 | Yes | TCTTTAGAGCTA**TTATTT**ATTTTTCATTGTTGTGT**TATCGT**GTCATTCGA |
| EcG10 | 149 | 18 | Yes | GGGTAG**TCGCGC**CCTCCTGTTTAATATCTTTAGAGC**TATTAT**TTATTTT |
| EcG10 | 146 | 22 | No | TCTGGGTAG**TCGCGC**CCTCCTGTTTAATATCTTTA**GAGCTA**TTATTTATT |
| EcG11 | 176 | 38 | No | CTAGGTTTACCT**TTACCT**TGCTAGGTTAAACTCCT**TAGTAT**TGTTTTCAT |
| EcG12 | 176 | 144 | No | TGACATCTT**TCACCA**GCATTGGTCGAAGGCAATTTG**TATAGT**GGTTGCGT |
| EcG12 | 107 | 44 | No | TATACTGGT**TTGTTG**GGTTCTATACAAAAAATGTT**TAATGT**GATGAACGG |
| EcG12 | 44 | 24 | No | TAGAGTACCACGTG**TGGTCC**GTCGTGGGTGGGTGC**TATAGT**TCAACGCAA |
| EcH02 | 198 | 14 | No | ATATGGTTTC**TCGGTT**TATGTATGGCGTCCTGAGT**GGGGCT**TTTACTGTT |
| EcH02 | 195 | 20 | No | TGAATATGGTTTC**TCGGTT**TATGTATGGCGTCCTGAG**TGGGGC**TTTTACT |
| EcH02 | 130 | 14 | No | GAACGGCT**TTGCGG**TTATAGGGTGTTGATATACTC**TAATCG**GTTACTTTC |
| EcH02 | 125 | 71 | Yes | GTTGGGAACGGCT**TTGCGG**TTATAGGGTGTTGATA**TACTCT**AATCGGTTA |
| EcH02 | 122 | 64 | Yes | TTTGTTGGGAACGGCT**TTGCGG**TTATAGGGTGTTGA**TATACT**CTAATCGG |
| EcH02 | 63 | 19 | No | GTCTA**TTAAAG**TGCAGGCACGTACATGTTTCTTG**GAATTT**GTTTATCGGC |
| EcH03 | 168 | 402 | Yes | TTCGTGGTTAGG**TTGTTG**TTTTTTCTGTAAACAA**TAAACT**AGTTAACTAG |
| EcH03 | 160 | 26 | Yes | GGCTGATAT**TCGTGG**TTAGGTTGTTGTTTTTTCTG**TAAACA**ATAAACTAG |

Table 7: continued from the previous page.

| Clone[1] | Pos.[2] | # Starts[3] | Shifted[4] | DNA Sequence Region, −50 to +1 (5′→3′)[5] |
|---|---|---|---|---|
| EcH03 | 101 | 30 | No | TGTGTTCCTTA**TTGCGC**CCAGTGATCTTCTTATGT**CAGTGT**ACATCACAA |
| EcH03 | 99 | 22 | No | TTTGTGTTCCTTA**TTGCGC**CCAGTGATCTTCTTATGT**CAGTGT**ACATCAC |
| EcH05 | 199 | 41 | Yes | GGGGTCTTGGC**TCACCG**CAAGGATACGGCAACCTA**TGTTAA**CTAGATTTT |
| EcH05 | 197 | 13 | No | TCGGGGTC**TTGGCT**CACCGCAAGGATACGGCAACC**TATGTT**AACTAGATT |
| EcH05 | 173 | 20 | No | TCTACGATA**TCACCT**CGAACGGCATCGGGGTCTTGGCTCACCGCAAGGAT |
| EcH05 | 143 | 91 | Yes | TCCTCT**TCGCGT**GTTACTTTGAAAATACGTTCTACGA**TATCAC**CTCGAAC |
| EcH05 | 139 | 41 | No | TTTCTCCTCT**TCGCGT**GTTACTTTGAAAATACGTTC**TACGAT**ATCACCTC |
| EcH05 | 133 | 134 | Yes | GCATCATTTCTCCTCT**TCGCGT**GTTACTTTGAAAA**TACGTT**CTACGATAT |
| EcH05 | 128 | 21 | No | TATAGGCA**TCATTT**CTCCTCTTCGCGTGTTACTTT**GAAAAT**ACGTTCTAC |
| EcH05 | 121 | 76 | No | GTTGGG**TTATAG**GCATCATTTCTCCTCTTCGCGTGT**TACTTT**GAAAATAC |
| EcH06 | 71 | 53 | No | TTGTGAAGT**TTGTAT**ACCAGTTAGGCTTAGGACCA**TACAAT**TATGTACGA |
| EcH07 | 71 | 51 | No | TTGTGAAGT**TTGTAT**ACCAGTTAGGCTTAGGACCA**TACAAT**TATGTACGA |
| EcH08 | 204 | 19 | No | GGC**TCGATG**TCCTGTTAATATAAGCTTTATGTAGCG**GAAACT**ATGGGAGG |
| EcH08 | 198 | 15 | No | CGGAGTGGC**TCGATG**TCCTGTTAATATAAGCTTTA**TGTAGC**GGAAACTAT |
| EcH08 | 184 | 446 | No | CTAGTGCAGGC**TCACGG**AGTGGCTCGATGTCCTGT**TAATAT**AAGCTTTAT |
| EcH08 | 147 | 148 | No | AGCTCGTTTCTAT**TTGATC**TAACACGACCCGTCTGC**TCTAGT**GCAGGCTC |
| EcH08 | 71 | 62 | No | GAA**TCGCTT**TGGGAAGTCCTATTCTTTTTTGAGTG**TAAGAT**CTAGCGTTC |
| EcH08 | 69 | 49 | No | TTGAA**TCGCTT**TGGGAAGTCCTATTCTTTTTTGAGTG**TAAGAT**CTAGCGT |
| EcH09 | 51 | 585 | Yes | TTTGTGCT**CCAGTG**TAGTATCCGAGGTTAATGTATGT**TATAGT**TTTAGGG |
| EcH12 | 202 | 65 | Yes | TTTTGTCGA**TCGAAC**TTGAGACGCCTTTGTGGGCTCC**CATAGC**GATGGAG |
| EcH12 | 101 | 1287 | Yes | TTAGTTAA**TCGAGT**ATACTAGGTTCGACTTGAATGG**TATCGT**TTCGTCGT |
| EcH12 | 98 | 109 | No | TATTTAGTTAA**TCGAGT**ATACTAGGTTCGACTTGAA**TGGTAT**CGTTTCGT |
| EcH12 | 78 | 123 | No | ACCTCTAGA**TGATCT**AAGTTTATTTAGTTAATCGAG**TATACT**AGGTTCGA |
| EcH12 | 56 | 43 | No | TGAATAGAAC**TCGCTC**TTTGCGACCTCTAGATGATC**TAAGTT**TATTTAGT |
| EcI03 | 135 | 923 | No | TTTGTA**TCACGG**TAGATTGTGCGTGACCTATTGCG**TATGAT**TGTAGGTAG |
| EcI03 | 95 | 117 | No | TTGAGGCGTG**TGGTTG**GTACAGTCGTTTGGCAGTGG**TATTTT**TGTATCAC |
| EcI04 | 191 | 14 | No | TTTTTGAGCGTGAT**TCGAAT**TAAAGGGCTATAA**TACTTT**CTCTGTCGGTT |
| EcI04 | 183 | 2801 | Yes | CTTCCTATTTT**TTGAGC**GTGATTCGAATTAAAGGGC**TATAAT**ACTTTCTC |
| EcI04 | 181 | 284 | No | TTCTTCCTATTTT**TTGAGC**GTGATTCGAATTAAAGGGC**TATAAT**ACTTTC |
| EcI07 | 197 | 18 | No | TTATTGGTATGTA**TCGTAC**CGTGGTTGCAATCAACGCT**TAACAA**TGAGAT |
| EcI07 | 144 | 63 | Yes | GTGTAGGATGGG**TTGACG**CGAATAGAGGGACAGTGT**TCGATA**TCTGGTCG |
| EcI07 | 142 | 39 | No | TTGTGTAGGATGGG**TTGACG**CGAATAGAGGGACAG**TGTTCG**ATATCTGGT |
| EcI07 | 113 | 14 | No | TGCAAGAAA**CGGTGT**GAATTTATGGATCTTTGTG**TAGGAT**GGGTTGACGC |
| EcI07 | 110 | 325 | Yes | GGATGCAAGAAA**CGGTGT**GAATTTATGGATCTTTGTG**TAGGAT**GGGTTGA |
| EcI10 | 83 | 56 | No | GGGCTGTACTAC**TAGCAG**GAGGGACCTGCCTCGTT**GATACT**GTGAAATGA |
| EcI11 | 59 | 2138 | Yes | GCCGCATG**CTAACT**AGTACGGTGTGATTTTATCAGT**TAAGAT**TGGGAGGT |
| EcI12 | 203 | 68 | Yes | CTTCGGAT**TAGTAG**GGCGAAGTGTCGTCAGTGGTGT**TAACTT**AATTGGAG |
| EcI12 | 135 | 81 | No | TTGG**TTGATT**TGATAGCGCAGGCAGACATGTGTGAA**TATAGT**ACGCGGGT |
| EcI12 | 133 | 127 | No | CGTTGG**TTGATT**TGATAGCGCAGGCAGACATGTGTG**GAATAT**AGTACGCGG |
| EcI12 | 131 | 27 | No | TTCGTTGG**TTGATT**TGATAGCGCAGGCAGACATGTGTG**GAATAT**AGTACGC |
| EcI12 | 76 | 15 | No | TAACGC**TTATAG**TAGGGTTTATGGTTGAGGCCATC**TCTTAA**GGGTAGGCT |
| EcI12 | 47 | 135 | Yes | AGGCTATAAGTA**TTGTTC**CCTATTTGGTGTAACGCT**TATAGT**AGGGTTTA |
| EcJ02 | 208 | 791 | No | GTGGCCATGAGC**TCGCGC**TTTGTATTCGTTACTGA**CAAATA**GGAGGTAGT |
| EcJ02 | 77 | 55 | No | TTCTGA**TTACGT**TTGTTAGACCGGTGTTCGGTAGTG**TAATGT**GGAAGGAT |

Table 7: continued from the previous page.

| Clone[1] | Pos.[2] | # Starts[3] | Shifted[4] | DNA Sequence Region, –50 to +1 (5′→3′)[5] |
|---|---|---|---|---|
| EcJ04 | 164 | 221 | Yes | AGGGGG**TTGTAC**TGTACAGCGTTCTCTATTATGCGT**TAAACT**TTGAATTC |
| EcJ09 | 160 | 1006 | Yes | AAATGA**TTGAGT**ATTTACCAGTACTGTTACAAGTGT**CATACT**TTGCTAGC |
| EcJ09 | 126 | 13 | No | TGTAAATGCGTTC**TTGCTT**TTGTTATGAAAACGTA**AATGAT**TGAGTATTT |
| EcJ09 | 60 | 71 | No | GTTGTTTTTAGC**TTGCGT**CTCTCTGGTTTCAGGTG**TACTTA**TGTATCCGT |
| EcJ10 | 22 | 81 | No | TCAGCGAAT**TCGCGG**CCGCTTCTAGAGAATAAAGAT**TATAGT**GTTTGTGC |
| EcJ11 | 38 | 22 | Yes | CGCTTC**TAGAGG**GCATGCAAATATTTCGGCGCGGGC**TATACG**TTATCAGG |
| EcJ12 | 201 | 24 | Yes | GTCCG**TTAGAG**TATGTCCCTGGAACAGTGTAATAT**GAGTAT**GTTGGGGGA |
| EcJ12 | 194 | 1514 | Yes | ACGTTGAGTCCG**TTAGAG**TATGTCCCTGGAACAGTG**TAATAT**GAGTATGT |
| EcJ12 | 191 | 24 | Yes | AGCACG**TTGAGT**CCGTTAGAGTATGTCCCTGGAACAG**TGTAAT**ATGAGTA |
| EcJ12 | 172 | 1653 | Yes | AACAT**CTGCCT**CTGGCCATAGCACGTTGAGTCCGT**TAGAGT**ATGTCCCTG |
| EcJ12 | 100 | 113 | No | TTGTAGGGTACT**TTGGTC**CCAGCTTTATGTTATGA**TATCGT**CCGTTGTAC |
| EcJ12 | 95 | 3351 | Yes | GTATG**TTGTAG**GGTACTTTGGTCCCAGCTTTATGT**TATGAT**ATCGTCCGT |
| EcJ12 | 93 | 17 | No | GCGTATG**TTGTAG**GGTACTTTGGTCCCAGCTTTATGT**TATGAT**ATCGTCC |
| EcK01 | 120 | 23 | No | TTGTGCT**TCACGG**ATTTTTCATGCAATGAGGTCGC**TACCTT**CTCAACTCC |
| EcK03 | 91 | 334 | No | ATGACTCATGTA**TTGTCC**TAAGGGGCACGTCTACTG**TATCCT**GAGACTGT |
| EcK04 | 90 | 21 | No | TTTTTGCGCGG**TAGACT**TATCGGTTTGATTCGTG**TATTTG**TTCAACTTAT |
| EcK05 | 198 | 17 | No | GTTAGTGTGG**CTGCCT**CTTCTTGTACTAGGGGCAC**TATATT**GTGGTATGA |
| EcK05 | 139 | 212 | No | TTGTGTCA**TCGTTT**TATTTTTCGGGAGAATGGTAAG**TATACT**GGAAAAGT |
| EcK05 | 128 | 239 | No | CTTCGTTTTCC**TTGTGT**CATCGTTTTATTTTTCGG**GAGAAT**GGTAAGTAT |
| EcK05 | 126 | 29 | No | TTCTTCGTTTTCC**TTGTGT**CATCGTTTTATTTTTCGG**GAGAAT**GGTAAGT |
| EcK06 | 154 | 34 | No | ATAGTTGGTTTC**TTGGTT**CTGCACCAAGTTTTTGTG**TATAAA**CAGTAGGC |
| EcK06 | 117 | 183 | Yes | TAGTA**TTGTCC**TATACACAGTTAATTGTTTGCGTGC**TATAGT**TGGTTTCT |
| EcK06 | 76 | 39 | No | ATAGATTCGCTT**TTGCTT**CGTCGATAGAGTTTTTA**TAGTAT**TAGTATTGT |
| EcK07 | 194 | 14 | No | GGTA**TGGCGT**AAGTTCCCTTATTTCGTCATGAGCTGTT**TAGGCA**CAATAT |
| EcK07 | 129 | 112 | No | AAGACGGAAT**TTGTAC**GTTGCCGTATCCGTGTATGC**TAAGGT**GTTTTCGC |
| EcK07 | 123 | 25 | No | TTACGCAAGA**CGGAAT**TTGTACGTTGCCGTATCCGTG**TATGCT**AAGGTGT |
| EcK08 | 201 | 1389 | Yes | AGGTGAAAACAAC**TTGGTT**TAAATGGGTATTGTGTA**TAGTAT**TGAACAGG |
| EcK08 | 199 | 13 | No | GGAGGTGAAAACAAC**TTGGTT**TAAATGGGTATTGTG**TATAGT**ATTGAACA |
| EcK08 | 60 | 47 | Yes | GTAAGGAT**TCGTTG**GACTAGTTGTTTTTCTCTAAT**TATATT**GACCCATTA |
| EcK08 | 58 | 77 | No | TTGTAAGGAT**TCGTTG**GACTAGTTGTTTTTCTCTAAT**TATATT**GACCCAT |
| EcK09 | 120 | 22 | No | GCTAGGTG**TTGTAG**TATGTCCCGTGGGTATGTGTTAA**TGGTTT**CCAGTGG |
| EcK09 | 117 | 111 | Yes | GTTGCTAGGTG**TTGTAG**TATGTCCCGTGGGTATGTGT**TAATGG**TTTCCAG |
| EcK09 | 112 | 67 | Yes | GAAGGGTTGCTAGGTG**TTGTAG**TATGTCCCGTGGGTA**TGTGTT**AATGGTT |
| EcK09 | 96 | 492 | Yes | TAAGTTGGA**TTATCG**AGAAGGGTTGCTAGGTGTTG**TAGTAT**GTCCCGTGG |
| EcK09 | 94 | 28 | No | TTTAAGTTGGA**TTATCG**AGAAGGGTTGCTAGGTGTTG**TAGTAT**GTCCCGT |
| EcK09 | 54 | 1273 | Yes | GTCTAAGTG**TCGACT**TGTGAGTTGAACTTTTTCTAC**TATTTT**TAAGTTGG |
| EcK09 | 51 | 30 | No | GGGGTCTAAGTG**TCGACT**TGTGAGTTGAACTTTTTC**TACTAT**TTTTAAGT |
| EcK11 | 69 | 18 | No | CTCATTAGAC**TTGCCC**CGTGCTAGTATACCTCAG**TAACGT**TCCTGTTTGT |
| EcK11 | 54 | 214 | No | AGTC**TTGGTG**AGGATCTCATTAGACTTGCCCCGTGC**TAGTAT**ACCTCAGT |
| EcK12 | 122 | 17 | No | TGTAAAAT**TTGCAG**TTTCGGATGTTCGTTTCGGT**TATCAT**GGTTTGACGC |
| EcK12 | 120 | 291 | No | GCTGTAAAAT**TTGCAG**TTTCGGATGTTCGTTTCGGT**TATCAT**GGTTTGAC |
| EcL01 | 178 | 18 | No | ATATTAGTAA**CTGGAT**GTCCTCTATTGGTTTAAGGG**TACTAT**TCTAGAGC |
| EcL01 | 157 | 51 | Yes | GGGTTATGAAACAT**TTGACT**AATATTAGTAACTGGA**TGTCCT**CTATTGGT |
| EcL01 | 141 | 131 | Yes | GGTAAATTCG**TCAAAT**GGGTTATGAAACATTTGAC**TAATAT**TAGTAACTG |

Table 7: continued from the previous page.

| Clone[1] | Pos.[2] | # Starts[3] | Shifted[4] | DNA Sequence Region, –50 to +1 (5′→3′)[5] |
|---|---|---|---|---|
| EcL01 | 37 | 934 | Yes | CCGCTTCTAGAGG**TTGCCT**AATGTTTATCTTAATCG**TATATT**GGTTTTTC |
| EcL02 | 203 | 68 | Yes | TCGGCACGCGGAG**TCATTG**CAATAATGGGACCAGC**TATGGT**CATGGGGAG |
| EcL02 | 127 | 53 | No | TGACATG**TCATGT**CCCAATTAGGTGTTTGACCTG**TAAGTT**GACTTATGTA |
| EcL03 | 132 | 538 | Yes | TAGATTGTTTAA**TCGATT**TTCTGGAAAGAGTTGTGC**TACACT**GTCCGGGC |
| EcL04 | 81 | 14 | No | TACAATGATC**TGGTTT**TCAACACCTGATTTTTATAA**TGTAAT**TGTATAGT |
| EcL07 | 141 | 65 | No | TGCTGGCGT**TTGCCT**AGACATTAGGTGGTGGTAAGA**TGGGGG**ATATATCT |
| EcL07 | 137 | 313 | No | AGTGTGCTGGCGT**TTGCCT**AGACATTAGGTGGTGG**TAAGAT**GGGGGATAT |
| EcL07 | 135 | 144 | No | CGAGTGTGCTGGCGT**TTGCCT**AGACATTAGGTGGTGG**TAAGAT**GGGGGAT |
| EcL09 | 167 | 62 | No | CAAGG**TTGTCC**TGGTAAAGGGTTGAGGTAGCGGAC**TATACT**GTTTATTAC |
| EcL09 | 116 | 1962 | Yes | GATGCAGCCCTTC**TTGCGG**TAATTCGTGGTTGGTGG**TATTGT**ATTGCTTG |
| EcL10 | 187 | 50 | No | GGCCCGTGTCC**TGGTTT**AGTTCGTCGGAATTGTGG**TATATT**TATAATCAA |
| EcM01 | 169 | 265 | No | CTTCAGTGCAATTT**TCAATG**TGCACTTGTATAGGG**TACGTT**ATTGATTAC |
| EcM03 | 99 | 13 | No | GAGCATTTTCT**TTGTCT**TCTGGAGTTATTCGCGAT**TATCTT**ATTCTTTAG |
| EcM06 | 135 | 778 | No | GGTATTAAT**TTGGTG**TAGTTTTTGACGGTTTAGTC**TATGAT**CTATGTTGA |
| EcM09 | 178 | 27 | No | ATGCAT**TTGACG**ATATGCTTAAGGCGTATCCTAAG**TAAAAT**TCCGTTTGT |
| EcM09 | 155 | 55 | No | ACGATTCGTT**TTATGT**TGCGAGGATGCATTTGACGA**TATGCT**TAAGGCGT |
| EcM09 | 119 | 46 | No | CGGGTCCAGGG**TTGAAG**TGACCAGTGCCTTTGTTC**TACGAT**TCGTTTTAT |
| EcM10 | 204 | 53 | No | TATGGTTCCTCGT**TCGCGG**GATGAATCCATGTACAT**TATAGC**GCTGGAGG |
| EcM11 | 210 | 100 | No | AGTGT**TTAAGC**GCGGTTTTTGCGTTTTTGTTGGAT**TATAAC**CAGGAGGTA |
| EcM11 | 145 | 37 | No | AGATAGTG**TGAACT**CATGATAAGCTCGCTTGTGGTA**TAAACT**GGGGTTAC |
| EcM11 | 129 | 27 | No | ACGGTTTCC**TCGGGT**AAGATAGTGTGAACTCATGA**TAAGCT**CGCTTGTGG |
| EcM11 | 79 | 249 | No | GTGTAGTAGCTG**TTGTGG**ATTACACTTTTATATGT**CATCTT**CATTAGTGT |
| EcN01 | 203 | 71 | Yes | TAACTCAA**TTGTGG**TGGGGAGTGGGGTTTCCCGCCCC**TGGCGA**CGTGGAG |
| EcN01 | 149 | 164 | No | GAGCAAGTA**TGGTGT**TTTTCAATACTATTGTGTTG**TAGATT**TGGATTCAC |
| EcN04 | 150 | 144 | No | TGGCTCCTGGGCCTT**TCGATT**ATCCTCTGTTCTCAT**TATAAT**TGTCTCTC |
| EcN05 | 127 | 101 | Yes | GTGTAATATAAA**CGGAGC**TGAGTTCGTGCGTACAGTT**TACAAT**GAGATAT |
| EcN05 | 94 | 201 | Yes | GGATCGACTTT**TCGTGT**AATGTTTTCTACCGGGGTG**TAATAT**AAACGGAG |
| EcN10 | 145 | 611 | Yes | GCTAGTGTAAATT**TTGTAG**GTTGTTGTGAAAGATGT**TATTTT**CTGTGTTC |
| EcN10 | 112 | 13 | No | ACATCAA**TCATGG**GTAAAATTTCATATAAGTCTGC**TAGTGT**AAATTTTGT |
| EcN11 | 100 | 332 | No | TTAATAAATG**CTGTGT**AAAACATTATCATTGAGTG**TATATT**AGTTGTTAC |
| EcN12 | 158 | 26 | Yes | TGTTTCGGA**TTGGCT**CGCAGTGTGCGGGCGACAAC**AAGAAT**TAGTGGTGC |
| EcN12 | 155 | 17 | No | TTCTGTTTCGGA**TTGGCT**CGCAGTGTGCGGGCGACAA**CAAGAA**TTAGTGG |
| EcN12 | 153 | 13 | No | GGTTCTGTT**TCGGAT**TGGCTCGCAGTGTGCGGGCGA**CAACAA**GAATTAGT |
| EcN12 | 114 | 15 | No | CCTATTTTATAAG**TTGCGA**AGCTATAATGGACGGGG**CATGGT**TCTGTTTC |
| EcN12 | 100 | 325 | Yes | CGCACCTA**CTGTAC**CCTATTTTATAAGTTGCGAAGC**TATAAT**GGACGGGG |
| EcO01 | 70 | 18 | No | AGCATGCATT**TTATCT**GCTGGGTATCTTAGATGAAG**TAAGTA**CTTGAGGG |
| EcO03 | 193 | 27 | No | TGGCTTTG**TCGGGT**TAGTTCGATCGTGTGAGTTAGA**TCGCTA**AAGTTAGT |
| EcO04 | 81 | 23 | No | CGCGTAT**CCGTTT**TTTTATTCGACGTACACTGCTT**TACAAT**CGTTCTTGT |
| EcO07 | 181 | 14 | No | TGTGTA**TCGTGT**GTTAACGGGATCCTCTTTTCGTGTT**TAGTTA**AGAGCGT |
| EcO07 | 116 | 720 | No | TACCCAGTAGATT**TTGCCT**TGAATTGGTTTGTTTTT**TATACT**GTCTCTGT |
| EcO08 | 186 | 107 | No | AACACGCCACTCTA**TTGCTT**ATACTGACTGTCACT**GACTCA**GATTCGGGT |
| EcO08 | 170 | 60 | No | AGGTAT**TTGTTT**TTTTAACACGCCACTCTATTGCT**TATACT**GACTGTCAC |
| EcO09 | 204 | 35 | No | TTTTCGGGA**CCGGCG**TTAGGTTTTATGAGCCTACA**GATGTT**ATCCGGAGG |
| EcO09 | 163 | 150 | Yes | TATAAGGTACGTC**TTGGAT**AGGTTTTAGAAGTGTGGT**CATATT**TTCGGGA |

Table 7: continued from the previous page.

| Clone[1] | Pos.[2] | # Starts[3] | Shifted[4] | DNA Sequence Region, –50 to +1 (5′→3′)[5] |
|---|---|---|---|---|
| EcO09 | 52 | 56 | No | AGAGCGTTCT**TCGCTG**GAGCTAGAGACGTACTTATG**GATAAT**GAACTCAT |
| EcO12 | 207 | 116 | No | TTTTGAGTAGCGT**TTATTT**AAGGGCCTTTGTAACT**TAATAT**AGGAGTCGA |
| EcO12 | 201 | 201 | Yes | TGTGTTTT**TTGAGT**AGCGTTTATTTAAGGGCCTTTG**TAACTT**AATATAGG |
| EcP01 | 90 | 21 | No | CAATTTGTCTTAC**TTGCGT**CATAACAGGGGTTAAG**TAATGA**TTTAGTCTA |
| EcP03 | 130 | 167 | No | TACAGGCT**TCAATG**GGAGTTCGTTAGAGGTTATGA**TATAGT**GGTGTTTAG |
| EcP03 | 124 | 82 | No | TATGTATACAGGCT**TCAATG**GGAGTTCGTTAGAGGT**TATGAT**ATAGTGGT |
| EcP05 | 168 | 1514 | No | CCAAGGT**TCGCCT**TTTTTAGAGTTCTATTTTTTGT**TATTGT**ATGCTGCGT |
| EcP05 | 166 | 275 | No | TTCCAAGGT**TCGCCT**TTTTTAGAGTTCTATTTTTTGT**TATTGT**ATGCTGC |
| EcP06 | 189 | 32 | No | TAGAATCTTATGAGTAAGATCAATGTATCTTATTG**TGTAAT**GGGGGCGGT |
| EcP06 | 179 | 23 | No | ACATTTATAG**TAGAAT**CTTATGAGTAAGATCAATG**TATCTT**ATTGTGTAA |
| EcP06 | 153 | 815 | Yes | ATGCCAATTTA**TTAGGT**ATCTTGTCGACATTTATAG**TAGAAT**CTTATGAG |
| EcP08 | 190 | 438 | Yes | ATCGATTTCTT**TCAACT**TCACTGTTCGCTACAATGA**TATACT**CATTGGGC |
| EcP08 | 183 | 965 | No | TCACGAGA**TCGATT**TCTTTCAACTTCACTGTTCGC**TACAAT**GATATACTC |
| EcP08 | 181 | 985 | No | TTTCACGAGA**TCGATT**TCTTTCAACTTCACTGTTCGC**TACAAT**GATATAC |
| EcP08 | 131 | 13 | No | TTATTCA**TTGTCT**TCTTCATACGGGTATTTCTCGAC**TAAACT**AGATTCTT |
| EcP08 | 127 | 47 | Yes | CCCTTTATTCA**TTGTCT**TCTTCATACGGGTATTTC**TCGACT**AAACTAGAT |
| EcP09 | 192 | 340 | Yes | CCCTGCCAT**TCATTG**TCATAGCATTGGAGTTTCGC**TACTGT**TGACGTTGC |
| EcP09 | 169 | 14 | No | AAATTG**TTGACG**CGTGGAGTATTCCCTGCCATTCAT**TGTCAT**AGCATTGG |
| EcP09 | 163 | 88 | No | TTTTACAAATTG**TTGACG**CGTGGAGTATTCCCTGC**CATTCA**TTGTCATAG |
| EcP09 | 56 | 22 | Yes | TCTTGCTTG**CTACTT**ATCATGAGGTACGTTGCATTC**TATACT**GGGAGGAG |
| EcP10 | 127 | 91 | No | GCATAGCGG**CTGTAG**CACTTCTTCGGCGTGACCTG**TAGAGT**TTGGAGTGG |
| EcP11 | 174 | 18 | No | GTGCGTT**TTGTGG**GAGTGTCAGGTGTGATACAGTAG**GGTCGT**AAAGGTGT |
| EcP11 | 166 | 2255 | Yes | TGTGTAAGGTGCGTT**TTGTGG**GAGTGTCAGGTGTGA**TACAGT**AGGGTCGT |
| EcP11 | 135 | 15 | No | TCCACTTTG**TTGACT**TATGAGCATTCAATATTGTG**TAAGGT**GCGTTTTGT |
| EcP12 | 123 | 503 | Yes | ACGTTAATGGGTC**TTGTTT**AGATCTTTTATCGTGGG**TAAGAT**CGAGTTGT |
| EcP12 | 105 | 20 | Yes | TGCTAATC**TCGGGG**GCTAACGTTAATGGGTCTTGTT**TAGATC**TTTTATCG |
| EcP12 | 66 | 35 | Yes | TGTATATTCTC**CGGAAT**AGTCTATGGTATGCCGAGT**TCTTGC**TAATCTCG |

[1] Clone: The ID of the identified *E. coli* clone.

[2] Pos.: Position of the TSS, within the ArtPromU sequence.

[3] Starts: Number of mapped RNAseq reads starting at the stated position.

[4] Shifted: Was the TSS shifted one position to the left (in case of neighbouring read stacks of similar height)?

[5] The artificial promoter sequence spanning from –50 to +1 (TSS). The identified motifs (by Improbizer[20]) are underlined, and coloured in blue and grey, representing different motifs. Darker colours represent stronger matches to the identified motif, TTGTGT and TATAAT.

##### 1.4.6 Motif analysis of the artificial promoters.

To identify potential promoter motifs in the regions upstream of the identified TSS, the tool Improbizer/Ameme[20] was used. The regions 50 nt upstream of the TSS were ordered in descending order based on the amount of read starts for the given TSS and Improbizer was run with standard parameters except that the parameter for "constrainer" was set to 100. The results of the motif search are listed in Table 7. As the identified consensus motifs, TTGTGT and TATAAT, strongly resembled the known *E. coli* $\sigma^{70}$ binding site, the motifs identified were parsed and split into six bins. These bins were created based on two categories: First, based on the rough equivalent of the promoter strength, i.e., the number of read starts; and Second, based on the presence of an extended –10 motif, i.e., a TGn directly upstream of the –10 box. The resulting bins were visualised via WebLogo[16], and the results are displayed in Figure 3.

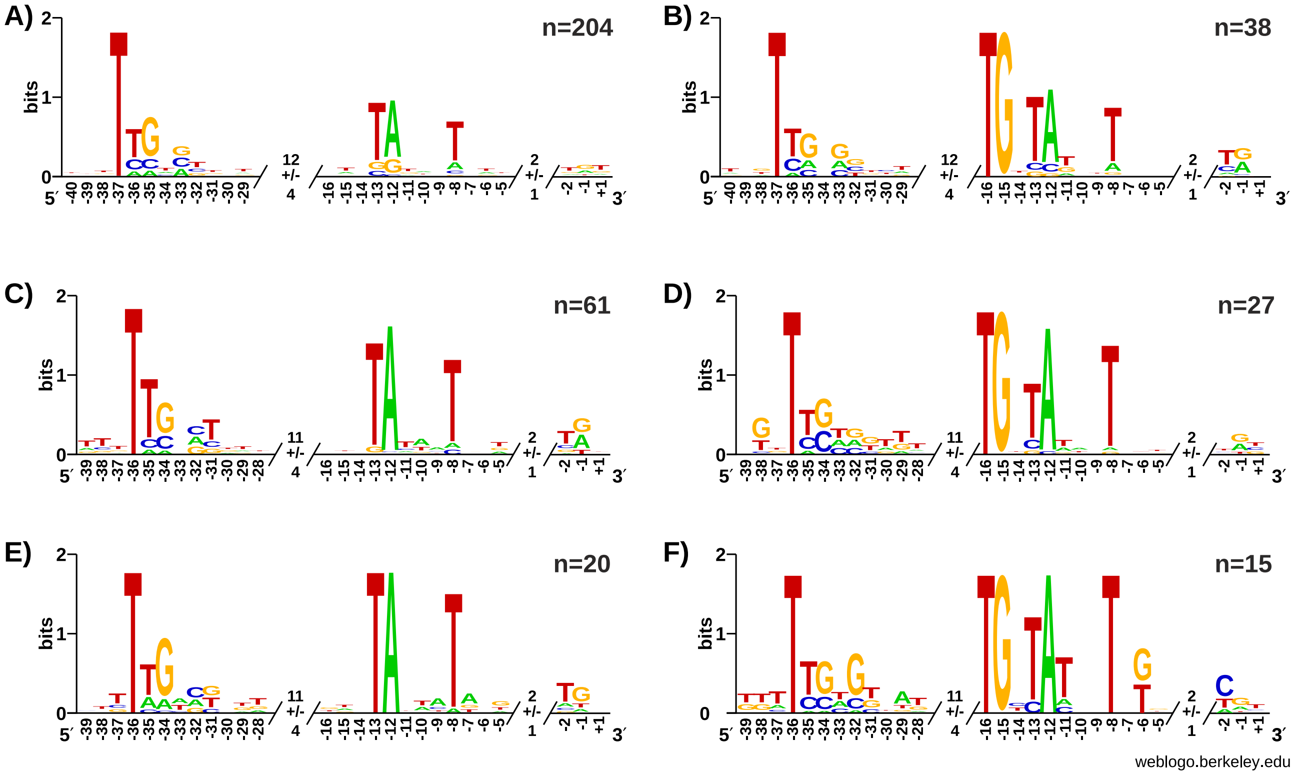

Figure 3: The WebLogo image of ArtPromU sequences with different promoter strengths identified in *E. coli*. The DNA sequences upstream of the identified TSS in the ArtPromU sequences was scanned for motifs using Improbizer [20]. The sequences were realigned based on the identified motifs, cut down to the core motifs including 3 nt up- and downstream, and binned based on relative "transcription strength", i.e. the number of mapped reads starting at each TSS as well as the presence or absence of the −10 extended motif TGn. For each bin, the median distance of the motifs was also calculated, the resulting numbers are given in the "spacers". The numbers on the x-axis are based on these median distances. Motifs of "weak" promoters (10 to 99 mapped reads) without (A) and with (B) −10 extended motif. Motifs of "medium" promoters (100 to 999 mapped reads) without (C) and with (D) −10 extended motif. Motifs of "strong" promoters (1000 mapped reads and more) without (E) and with (F) −10 extended motif. The number of sequences used in each alignment are depicted with "n".

##### 1.4.7 Motif analysis of the artificial 5′ UTRs.

To identify potential motifs in the regions downstream of the identified TSS, also known as the initially transcribed region, the tool Improbizer/Ameme [20] was used. The region spanning from +1 to +25 were analysed searching for one motif and one occurrence. This search resulted in the motif GGATGT and the result of this motif search for each artificial 5′ UTR is listed in Table 8.

Table 8: The list of all the motifs identified within the artifical 5′ UTRs.

| Clone[1] | Position[2] | # Starts[3] | Shifted[4] | DNA Sequence Region, +1 to +25 (5′→3′)[5] |
|---|---|---|---|---|
| EcC09 | 180 | 15242 | Yes | TAAACGAA**GTAGGT**CAATCCGGAGTA |
| EcA12 | 176 | 7647 | No | TGACGG**GTAAGT**AGAGTGGGTAAAGG |
| EcJ12 | 95 | 3351 | Yes | TTGTACTCTAAA**CGGTTT**GCATCTTG |
| EcA12 | 178 | 3320 | Yes | ACGGGTAAG**TAGAGT**GGGTAAAGGAG |
| EcE07 | 117 | 3227 | No | CCACTAT**CGCCCT**AGCACGGTGGTAG |
| EcE09 | 125 | 3126 | No | TGAAAGG**CGGGTT**ACTTACGCTTATC |
| EcC09 | 177 | 3081 | Yes | CCGTAAA**CGAAGT**AGGTCAATCCGGA |

Table 8: continued from the previous page.

| **Clone**[1] | **Position**[2] | **# Starts**[3] | **Shifted**[4] | **DNA Sequence Region, +1 to +25 (5′→3′)**[5] |
|---|---|---|---|---|
| EcI04 | 183 | 2801 | Yes | CTGTCGGTT**CGCTGT**TGGAGGGGTAT |
| EcE03 | 117 | 2640 | Yes | AAGTCAACCT**GGCCAT**AATTCGGTTG |
| EcP11 | 166 | 2255 | Yes | TAAAGGTG**TTCTGT**TGTAGGAAATCC |
| EcI11 | 59 | 2138 | Yes | TTGGGTGATTTC**GGGAGT**TTGTGGGC |
| EcE01 | 31 | 2096 | Yes | TTGGATC**TTGCGT**CGTTCTGGTAGCT |
| EcG07 | 126 | 2051 | Yes | CTAGAGT**TTCTGT**TCAGTATTGGTTA |
| EcL09 | 116 | 1962 | Yes | GGCAAGGTTGTCCTGGTAAAGGGTTG |
| EcF03 | 96 | 1893 | No | TAATAA**TTGCAT**CTCTCTGGACGAAC |
| EcD09 | 66 | 1869 | No | CGATGTTG**CGACTT**TTATTAAAATCG |
| EcC07 | 169 | 1667 | No | TCTTAGGAAT**TAATTT**TTGTAGCATA |
| EcJ12 | 172 | 1653 | Yes | GGAACAGTG**TAATAT**GAGTATGTTGG |
| EcC02 | 90 | 1581 | Yes | TTGCGGCTG**CAGGTT**GAATGTTGTGG |
| EcP05 | 168 | 1514 | No | TCTAAAT**TTCGAT**TGTCGATCTAAAA |
| EcG10 | 175 | 1480 | Yes | ATGATGAAC**GTAGAT**CGTTGACTGAG |
| EcC10 | 139 | 1306 | Yes | CGAGGTGATGT**TGCGAT**TGAAGCTCA |
| EcH12 | 101 | 1287 | Yes | TTTACGA**CGAATT**AGAAAACGTGGTG |
| EcK09 | 54 | 1273 | Yes | GATTATCGAG**AAGGGT**TGCTAGGTGT |
| EcE07 | 90 | 1117 | Yes | GAGTATC**CTGTGT**TAATTTGGGACAT |
| EcE07 | 115 | 1101 | No | TACCACTAT**CGCCCT**AGCACGGTGGT |
| EcF02 | 104 | 1097 | No | TAGTCTA**TGATGT**AGATATGGACTTG |
| EcE10 | 182 | 1010 | Yes | CAGAACGGTTCTGCTAACGGAGGTTT |
| EcJ09 | 160 | 1006 | Yes | CACGTGTACTAAGGGGCGACCGGTAC |
| EcP08 | 181 | 985 | No | CTCATT**GGGCGT**GATGAGGAGGGGTA |
| EcP08 | 183 | 965 | No | CATT**GGGCGT**GATGAGGAGGGGTATA |
| EcD08 | 78 | 961 | Yes | GTCGCTTGCGT**CTGTGT**CGTGATCAT |
| EcL01 | 37 | 934 | Yes | CCTAT**GACCAT**AGTTATAACATTCGA |
| EcI03 | 135 | 923 | No | GGAGCGGTC**GTGCAT**AAGCCGCGTGT |
| EcE07 | 102 | 826 | Yes | TTAATTTG**GGACAT**ACCACTATCGCC |
| EcP06 | 153 | 815 | Yes | GTAAGAT**CAATGT**ATCTTATTGTGTA |
| EcM06 | 135 | 778 | No | AAGGTTCAT**CTAGTT**TCGCCGCAGGG |
| EcA10 | 132 | 739 | No | CCGGTAA**GTGTTT**CTTTGGCCAGACT |
| EcO07 | 116 | 720 | No | TTTTCGTAAG**TGGTGT**TGTGTATCGT |
| EcF02 | 106 | 656 | Yes | GTCTATGAT**GTAGAT**ATGGACTTGTG |
| EcN10 | 145 | 611 | Yes | CACATC**GAAGGT**ATTCAAGAGTTAGT |
| EcH09 | 51 | 585 | Yes | GGGTTCTTTTTTCTCGGAGGGTCTAA |
| EcL03 | 132 | 538 | Yes | CTTGAGGATAT**TTAGCT**CTCAACTGT |
| EcP12 | 123 | 503 | Yes | TACAGGGGGGGC**AACAAT**AATCAAGG |
| EcK09 | 96 | 492 | Yes | GGTATGTGTTAA**TGGTTT**CCAGTGGA |
| EcE10 | 113 | 469 | Yes | GGGCGTCTAGT**AGGCCT**CTTCCGCTT |
| EcB04 | 192 | 458 | No | -TACTGGTT**GGAGGT**TCTCTATGATG |
| EcH08 | 184 | 446 | No | TGTAGCG**GAAACT**ATGGGAGGGCAAA |
| EcP08 | 190 | 438 | Yes | -CGTGATGAGGAG**GGGTAT**AATGATG |
| EcF02 | 122 | 421 | No | TGGACTTGT**GAGCCT**AATGTATAGAG |
| EcD01 | 191 | 408 | No | -GTTTAGA**TGGAGT**CGTGGTATGATG |

Table 8: continued from the previous page.

| Clone[1] | Position[2] | # Starts[3] | Shifted[4] | DNA Sequence Region, +1 to +25 (5′→3′)[5] |
|---|---|---|---|---|
| EcG07 | 171 | 405 | Yes | GAACGGTAGTTAG**TGGGCT**CTGAAAT |
| EcF04 | 178 | 404 | No | GGCCTTGTCAT**ATCGAT**CGGAAGGAG |
| EcH03 | 168 | 402 | Yes | GAGTTATG**TGGTTT**TATAACTCGATG |
| EcK03 | 91 | 334 | No | TCTT**CAATTT**CTCATGGGAATTACGT |
| EcN11 | 100 | 332 | No | CTTTCATCC**TAGTCT**CGATTTGCTTT |
| EcI07 | 110 | 325 | Yes | ACGCGAATAGAGG**GACAGT**GTTCGAT |
| EcN12 | 100 | 325 | Yes | GCATGGTT**CTGTTT**CGGATTGGCTCG |
| EcL07 | 137 | 313 | No | TATCTGCGTG**GGGTAT**CGGTAGTGAT |
| EcB01 | 101 | 307 | No | TTAGCTAGT**TAGTGT**TTGGTGTAGTC |
| EcF09 | 120 | 290 | Yes | CTAGGGATGTG**CTAGCT**TGAAAGGTG |
| EcI04 | 181 | 284 | No | CTCTGTCGGTT**CGCTGT**TGGAGGGGT |
| EcP05 | 166 | 275 | No | CGTCTAAAT**TTCGAT**TGTCGATCTAA |
| EcM01 | 169 | 265 | No | CGTCGT**GGATGT**CTCAGTCACGACCT |
| EcM11 | 79 | 249 | No | TACGGTTTCCTCGGGTAAGATAGTGT |
| EcK05 | 128 | 239 | No | TACTGG**AAAAGT**TGGTCGTCAGTTAG |
| EcD09 | 68 | 226 | No | ATGTTG**CGACTT**TTATTAAAATCGGT |
| EcB08 | 61 | 221 | No | GGCTT**TGGTCT**TAGTGGCGGCACGTC |
| EcJ04 | 164 | 221 | Yes | CATATAGGAT**GAATTT**ATACAAAAAG |
| EcK11 | 54 | 214 | No | TAACGTTC**CTGTTT**GTGGAGTGATAT |
| EcC05 | 51 | 212 | No | TAAGTGTGTA**GGCCCT**AGTTTTTGCT |
| EcK05 | 139 | 212 | No | TTGGTCGTCAGT**TAGTGT**GGCTGCCT |
| EcF09 | 123 | 209 | No | GGGATGTG**CTAGCT**TGAAAGGTGCAG |
| EcC04 | 117 | 204 | No | TTTGACTTTTCGT**CGACAT**TCATGGG |
| EcN05 | 94 | 201 | Yes | GCTGAGTTC**GTGCGT**ACAGTTTACAA |
| EcA12 | 174 | 199 | No | AGTGACGG**GTAAGT**AGAGTGGGTAAA |
| EcG08 | 133 | 195 | No | TAGACCG**CACGGT**GGGAAGTCACTGT |
| EcF08 | 86 | 185 | No | TGGCTTCCTCGA**GTGGTT**CGGATTTG |
| EcK06 | 117 | 183 | Yes | TTGGTTCTGCAC**CAAGTT**TTTGTGTA |
| EcC10 | 137 | 176 | No | TTCGAGG**TGATGT**TGCGATTGAAGCT |
| EcF12 | 56 | 168 | No | GTTCGATGGGTTA**GGCGAT**CGGTGCA |
| EcP03 | 130 | 167 | No | GGGTGGGCATT**GTACTT**GAGTAATTA |
| EcE12 | 158 | 164 | No | ACGTCTGTC**TGGCAT**GGGGTATGAAT |
| EcN01 | 149 | 164 | No | CTGTGTAA**CTCAAT**TGTGGTGGGGAG |
| EcC11 | 165 | 163 | Yes | AGCACTCG**GTAAAT**GAGAGAGCAACG |
| EcD04 | 58 | 161 | No | TCATCTTTG**CGGCCT**TCACAATGAAG |
| EcE01 | 95 | 154 | Yes | TCCGGTTGG**CGGCCT**TATTCCGGCGA |
| EcO09 | 163 | 150 | Yes | ACCGGCG**TTAGGT**TTTATGAGCCTAC |
| EcH08 | 147 | 148 | No | CACGGAGTGG**CTCGAT**GTCCTGTTAA |
| EcG12 | 176 | 144 | No | TGCTTCGTAG**GAGTCT**TAGTTTGAGG |
| EcL07 | 135 | 144 | No | TATATCTGCG**TGGGGT**ATCGGTAGTG |
| EcN04 | 150 | 144 | No | CCGGTATT**TTAAAT**TACTTGGGCTTC |
| EcI12 | 47 | 135 | Yes | ATGGTTGA**GGCCAT**CTCTTAAGGGTA |
| EcH05 | 133 | 134 | Yes | TCACCTCGAA**CGGCAT**CGGGGTCTTG |
| EcD05 | 91 | 132 | No | TGGGTT**CGAGAT**CCCCAGTATGACTC |

Table 8: continued from the previous page.

| **Clone**[1] | **Position**[2] | **# Starts**[3] | **Shifted**[4] | **DNA Sequence Region, +1 to +25 (5′→3′)**[5] |
|---|---|---|---|---|
| EcL01 | 141 | 131 | Yes | GGATGTC**CTCTAT**TGGTTTAAGGGTA |
| EcF02 | 166 | 127 | Yes | ATGGGTGCT**CAGTGT**GCAAATTGGGC |
| EcI12 | 133 | 127 | No | GGTCGAT**TTACAT**GCGAACGCCTTCG |
| EcC02 | 148 | 124 | No | TTTTCTCTTCCAA**CGGTAT**TTTGTAT |
| EcC02 | 146 | 123 | No | GGTTTTCTCTTCCAA**CGGTAT**TTTGT |
| EcH12 | 78 | 123 | No | ACTTGAA**TGGTAT**CGTTTCGTCGTTT |
| EcB01 | 115 | 122 | No | TTTGGTGTAGTC**GTAAAT**TATTTTTA |
| EcC08 | 106 | 120 | Yes | GGATAGAG**TTCCGT**TAGTGTGTTGAG |
| EcE04 | 179 | 120 | Yes | TCGAAATACTTG**TACTCT**GAAGGAGA |
| EcI03 | 95 | 117 | No | CGGTAGATT**GTGCGT**GACCTATTGCG |
| EcJ12 | 100 | 113 | No | CTCTAAACGGT**TTGCAT**CTTGGCAAC |
| EcK07 | 129 | 112 | No | CCTAGC**TGGTGT**TTAGGGTATGGCGT |
| EcD08 | 177 | 109 | No | TCAAGGCAATGAG**TGCTAT**AATTGGA |
| EcH12 | 98 | 109 | No | TCGTTTACGA**CGAATT**AGAAAACGTG |
| EcO08 | 186 | 107 | No | TTATC**CGCTCT**ACGGGAGGAATATTA |
| EcN05 | 127 | 101 | Yes | TCTCTGGTTTGT**TGGACT**GAGCTAGA |
| EcA01 | 98 | 95 | Yes | -CATTTTAGG**GAGTAT**CTCCATGATG |
| EcH05 | 143 | 91 | Yes | CGGCATCG**GGGTCT**TGGCTCACCGCA |
| EcP10 | 127 | 91 | No | GGCGGGTCAAT**CAAAGT**CAAACAGGG |
| EcF10 | 122 | 89 | No | TACCATGTGA**CTAGGT**TCAAGTTTGA |
| EcP09 | 163 | 88 | No | GCATTGGAGT**TTCGCT**ACTGTTGACG |
| EcD01 | 188 | 86 | Yes | GTGGTTTAGA**TGGAGT**CGTGGTATGA |
| EcB11 | 167 | 85 | No | CACTCG**GTAAAT**GAGAGAGCAACGCT |
| EcB07 | 135 | 83 | Yes | TTGAGAGG**TGACCT**GCTCTGGCCCCG |
| EcC08 | 122 | 82 | Yes | GTGTGTTGAG**GGAGGT**TGTTACATTG |
| EcP03 | 124 | 82 | No | TGTTTAGGGT**GGGCAT**TGTACTTGAG |
| EcB04 | 178 | 81 | No | TGATAACCC**AGGTGT**ACTGGTTGGAG |
| EcF08 | 88 | 81 | No | GCTTCCTCGA**GTGGTT**CGGATTTGCT |
| EcG02 | 125 | 81 | Yes | ACTTTC**CGGTCT**AGGCTGATCTGAAT |
| EcI12 | 135 | 81 | No | TCGAT**TTACAT**GCGAACGCCTTCGGA |
| EcJ10 | 22 | 81 | No | CCTGTCAAT**CGATGT**GTGGGGAGTTC |
| EcK08 | 58 | 77 | No | TTAAGCCCATC**TTCTAT**GTCTCAGGC |
| EcH05 | 121 | 76 | No | CGTTCTA**CGATAT**CACCTCGAACGGC |
| EcF04 | 175 | 71 | Yes | TTAGGCCTTG**TCATAT**CGATCGGAAG |
| EcG07 | 122 | 71 | Yes | AATGCTAGAGT**TTCTGT**TCAGTATTG |
| EcH02 | 125 | 71 | Yes | ACTTTC**CGGTCT**AGGCTGATCTGAAT |
| EcJ09 | 60 | 71 | No | TGGTTCTT**GTAAGT**TTTTGTAAATGC |
| EcF01 | 171 | 70 | No | TCACATGT**GAACAT**GTGTTAGGTATG |
| EcA02 | 63 | 68 | No | ATCTGTG**CTCTCT**TATTGGTAACAAT |
| EcC02 | 177 | 67 | No | CTCTATTC**GGCTAT**AACATAGTGGGA |
| EcD07 | 127 | 67 | No | GGCGGGTCAAT**CAAAGT**CAAACAGGG |
| EcB04 | 140 | 66 | Yes | GTTGTACT**CTGTAT**GGTTTCCTCTTA |
| EcC07 | 184 | 66 | Yes | TTTGTAGCA**TAATGT**AGGAGCTCGAT |
| EcE07 | 186 | 66 | Yes | GGATATCCACGT**GGAGGT**CGAAGATG |

Table 8: continued from the previous page.

| Clone[1] | Position[2] | # Starts[3] | Shifted[4] | DNA Sequence Region, +1 to +25 (5′→3′)[5] |
|---|---|---|---|---|
| EcC05 | 190 | 65 | No | TACTATCTTAT**GGAGGT**AACTAATGA |
| EcG02 | 122 | 65 | Yes | GTTACTTTC**CGGTCT**AGGCTGATCTG |
| EcL07 | 141 | 65 | No | TGCGTG**GGGTAT**CGGTAGTGATATCG |
| EcC07 | 65 | 64 | Yes | GGTGAATTCG**TTAGAT**GGCTGAGGTT |
| EcH02 | 122 | 64 | Yes | GTTACTTTC**CGGTCT**AGGCTGATCTG |
| EcI07 | 144 | 63 | Yes | GAAATTAT**TGGTAT**GTATCGTACCGT |
| EcC04 | 112 | 62 | Yes | CCTTGTT**TGACTT**TTCGTCGACATTC |
| EcH08 | 71 | 62 | No | CTTTCCTCG**TGGGGT**CACTGCTGTCG |
| EcL09 | 167 | 62 | No | CTGTATGC**GGGAGT**TTTGCGTCGATC |
| EcE04 | 56 | 60 | No | TGACTAT**GGGTGT**CGTTTTGAGGTGC |
| EcO08 | 170 | 60 | No | CTGACTCAGATT**CGGGTT**ATCCGCTC |
| EcC09 | 101 | 59 | No | TATCGT**CTAATT**GTCGGGTGGGTTTG |
| EcF01 | 111 | 59 | No | CTCCTT**TTGGAT**TCGGCTACGCCTTG |
| EcB11 | 165 | 56 | No | AGCACTCG**GTAAAT**GAGAGAGCAACG |
| EcE07 | 100 | 56 | No | TGTTAATTTG**GGACAT**ACCACTATCG |
| EcI10 | 83 | 56 | No | AGATATTT**CTGCTT**GGTCCGGCGAAT |
| EcO09 | 52 | 56 | No | TCGTCT**GGATAT**GTTTGATTTTCTAC |
| EcA05 | 168 | 55 | Yes | TTGAG**CAGCAT**GGTCTACTCTAGGTC |
| EcA06 | 71 | 55 | No | AGGGTT**TGATAT**GGGATGATCAGTCA |
| EcJ02 | 77 | 55 | No | TTCACTC**TTAGGT**TGGAGTCGATTGC |
| EcM09 | 155 | 55 | No | TATCCTAAG**TAAAAT**TCCGTTTGTCA |
| EcE04 | 185 | 53 | No | TACTTG**TACTCT**GAAGGAGACGATTG |
| EcF08 | 155 | 53 | No | CTTAGGGAGC**TTACCT**GATGTCTGTA |
| EcH06 | 71 | 53 | No | AGGGTT**TGATAT**GGGATGATCAGTCA |
| EcL02 | 127 | 53 | No | ACTGGCTACGGC**GGCGAT**TTATTTTA |
| EcE09 | 127 | 52 | No | AAAGGCGGG**TTACTT**ACGCTTATCGG |
| EcH07 | 71 | 51 | No | AGGGTT**TGATAT**GGGATGATCAGTCA |
| EcL01 | 157 | 51 | Yes | TTTAAGGG**TACTAT**TCTAGAGCTATT |
| EcA11 | 48 | 50 | No | TATGGGTCTG**AGCGCT**GAGAACTACG |
| EcL10 | 187 | 50 | No | AGAGAAT**TTCTAT**GGAGAGTGTTAAT |
| EcA07 | 71 | 49 | No | AGGGTT**TGATAT**GGGATGATCAGTCA |
| EcB04 | 190 | 49 | No | TGTACTGGTT**GGAGGT**TCTCTATGAT |
| EcH08 | 69 | 49 | No | TTCTTTCCTCG**TGGGGT**CACTGCTGT |
| EcK08 | 60 | 47 | Yes | AAGCCCATCTT**CTATGT**CTCAGGCTC |
| EcP08 | 127 | 47 | Yes | TTCTTTTTCA**CGAGAT**CGATTTCTTT |
| EcM09 | 119 | 46 | No | TGTTGC**GAGGAT**GCATTTGACGATAT |
| EcC07 | 29 | 44 | Yes | GTTTACTTGC**GTGGAT**GTATTAGCTT |
| EcE09 | 149 | 44 | Yes | TCGGCGG**CGAGCT**GCACGGGCGCTTG |
| EcG12 | 107 | 44 | No | GATGTTTCTTA**GGGGAT**GCCTGACAT |
| EcE03 | 155 | 43 | Yes | CGCGGTTA**ATGGGT**GCCCGTATTGTG |
| EcH12 | 56 | 43 | No | TTAATC**GAGTAT**ACTAGGTTCGACTT |
| EcC04 | 192 | 42 | Yes | -TTGTAGTG**GGAGAT**CTACTATGATG |
| EcH05 | 139 | 41 | No | CGAACGGCAT**CGGGGT**CTTGGCTCAC |
| EcE06 | 88 | 40 | No | GGGGTATGTA**CACAGT**TCGTGCTTAT |

Table 8: continued from the previous page.

| Clone[1] | Position[2] | # Starts[3] | Shifted[4] | DNA Sequence Region, +1 to +25 ($5'\to3'$)[5] |
|---|---|---|---|---|
| EcC08 | 115 | 39 | No | TCCGTTA**GTGTGT**TGAGGGAGGTTGT |
| EcI07 | 142 | 39 | No | TCGAAATTAT**TGGTAT**GTATCGTACC |
| EcK06 | 76 | 39 | No | TCCTATA**CACAGT**TAATTGTTTGCGT |
| EcA12 | 171 | 38 | Yes | GTGAGTGACGG**GTAAGT**AGAGTGGGT |
| EcG11 | 176 | 38 | No | TTGGCAT**TTGGCT**CACACTCATATGG |
| EcC08 | 109 | 37 | No | TAGAGTTCCGT**TAGTGT**GTTGAGGGA |
| EcD06 | 162 | 37 | No | TGGAGTTGG**CTACGT**GGTGGCGTTTA |
| EcF08 | 152 | 37 | No | AAACTTAG**GGAGCT**TACCTGATGTCT |
| EcM11 | 145 | 37 | No | CTTCTT**GAAGCT**TTGAAGTGTTTAAG |
| EcF03 | 150 | 36 | No | TGTTTGCT**GTGCCT**TCTTTTTCGGAG |
| EcG08 | 135 | 36 | No | GACCGCACGGTG**GGAAGT**CACTGTGG |
| EcG08 | 131 | 35 | No | AATAGACCG**CACGGT**GGGAAGTCACT |
| EcP12 | 66 | 35 | Yes | GGGGGC**TAACGT**TAATGGGTCTTGTT |
| EcC09 | 174 | 34 | Yes | TAACCGTAAA**CGAAGT**AGGTCAATCC |
| EcE04 | 176 | 34 | Yes | ATTT**CGAAAT**ACTTGTACTCTGAAGG |
| EcK06 | 154 | 34 | No | CCAAGTAT**CACGCT**TTGTTTATCGAA |
| EcC04 | 115 | 33 | No | TGTTTGACTTTTCGT**CGACAT**TCATG |
| EcB11 | 163 | 32 | No | TAAGCACTCG**GTAAAT**GAGAGAGCAA |
| EcE01 | 114 | 32 | Yes | CCGGCGAGTAGTCGGCTGCTACGTGC |
| EcP06 | 189 | 32 | No | TTTACAGTTAA**GGAGGT**GTTTTATGA |
| EcB01 | 165 | 31 | No | TTCACAGTACT**TGCTAT**CGGAGGTTA |
| EcE03 | 164 | 31 | Yes | TGGGTGCCCGTA**TTGTGT**CCTTCTTT |
| EcF07 | 189 | 31 | Yes | TGGGTGTGGAA**GGAGAT**CATTGATGA |
| EcC02 | 190 | 30 | No | TAACATAGTGG**GAGGGT**CACTATGAT |
| EcE07 | 87 | 30 | Yes | TTAGAGTATC**CTGTGT**TAATTTGGGA |
| EcG01 | 158 | 30 | No | TCGTCAA**GGATGT**TGCTTGGTCCATG |
| EcH03 | 101 | 30 | No | ATCACATT**TTGGCT**GATATTCGTGGT |
| EcK09 | 51 | 30 | No | TTGGATTATCGAGAAGGGTTGCTAGG |
| EcA02 | 117 | 29 | No | TGAACGTACAAGA**GGAGAT**ACCCGAT |
| EcK05 | 126 | 29 | No | TATACTGG**AAAAGT**TGGTCGTCAGTT |
| EcK09 | 94 | 28 | No | TGGGT**ATGTGT**TAATGGTTTCCAGTG |
| EcB07 | 22 | 27 | No | GAGTGTATTTGCG**GAGCGT**TGCTAAT |
| EcI12 | 131 | 27 | No | CGGGTCGAT**TTACAT**GCGAACGCCTT |
| EcM09 | 178 | 27 | No | TCATTGTG**CTCGGT**CCGAAAAAGGAG |
| EcM11 | 129 | 27 | No | GTATAAAC**TGGGGT**TACTTCTTGAAG |
| EcB04 | 180 | 26 | No | ATAACCC**AGGTGT**ACTGGTTGGAGGT |
| EcD04 | 190 | 26 | No | TGGGTTGAGA**GGAGGT**TTTGAATGAT |
| EcD04 | 103 | 26 | No | TTCGTTTCTTTC**TTCCGT**TTGTGCAG |
| EcE01 | 101 | 26 | No | TGG**CGGCCT**TATTCCGGCGAGTAGTC |
| EcE10 | 191 | 26 | No | TCTGCTAAC**GGAGGT**TTCGAATGATG |
| EcE10 | 109 | 26 | No | CTAA**GGGCGT**CTAGTAGGCCTCTTCC |
| EcF09 | 146 | 26 | No | CAGGTCC**CCCGAT**GCGCGTTTTTGA |
| EcH03 | 160 | 26 | Yes | GTTAACTAGAG**TTATGT**GGTTTTATA |
| EcN12 | 158 | 26 | Yes | CGGGATGA**GTACGT**CTTAGGTTGGT |

Table 8: continued from the previous page.

| Clone[1] | Position[2] | # Starts[3] | Shifted[4] | DNA Sequence Region, +1 to +25 (5′→3′)[5] |
|---|---|---|---|---|
| EcA12 | 192 | 25 | No | -TGGGTAAAGGA**GTGAGT**GTATGATG |
| EcC11 | 163 | 25 | No | TAAGCACTCG**GTAAAT**GAGAGAGCAA |
| EcC11 | 97 | 25 | No | AGGGAG**CACTTT**TTAGGGCTGGTTAT |
| EcK07 | 123 | 25 | No | TTTTCGC**CTAGCT**GGTGTTTAGGGTA |
| EcA12 | 184 | 24 | No | AAGTAGA**GTGGGT**AAAGGAGTGAGTG |
| EcB11 | 101 | 24 | Yes | AGCACTTTTT**AGGGCT**GGTTATCGCC |
| EcC04 | 174 | 24 | Yes | TTTTTCA**TTCAGT**GCAACTTGTAGTG |
| EcG12 | 44 | 24 | No | AGCAATTGC**CTGAAT**ATACTGGTTTG |
| EcJ12 | 191 | 24 | Yes | -ATGTTGGGG**GAGGAT**GGGTATGATG |
| EcA02 | 84 | 23 | No | ACAATTT**TACTAT**CGCTAAATGGGCT |
| EcC08 | 133 | 23 | Yes | GAGGTTG**TTACAT**TGCTTGGTTGTAT |
| EcF03 | 113 | 23 | Yes | TGGACGAACTTGT**CGGGCT**ATTGTAG |
| EcG01 | 161 | 23 | No | TCAAGGATG**TTGCTT**GGTCCATGGAT |
| EcG07 | 117 | 23 | No | ATGTGAATGC**TAGAGT**TTCTGTTCAG |
| EcK01 | 120 | 23 | No | CGGGGT**GTCATT**CACGGCAGCGGGGT |
| EcO04 | 81 | 23 | No | TAACAAGTC**GTCGTT**TGTGTCTTGGC |
| EcP06 | 179 | 23 | No | ATGGGGG**CGGTTT**ACAGTTAAGGAGG |
| EcA10 | 178 | 22 | No | TCATGTCACG**GTGGGT**TAATTGGGAG |
| EcG10 | 146 | 22 | No | TTTTCATTG**TTGTGT**TATCGTGTCAT |
| EcH03 | 99 | 22 | No | CAATCACATT**TTGGCT**GATATTCGTG |
| EcJ11 | 38 | 22 | Yes | GAGCTTCGG**CTCGTT**GGCGTGGAATC |
| EcP09 | 56 | 22 | Yes | GTTGA**TGCCCT**TTGTAAGTGGGCTTA |
| EcB07 | 110 | 21 | No | TTCATGCATTA**CAGTAT**CGCTAGAGT |
| EcB11 | 91 | 21 | No | TCTGTAAGGGAG**CACTTT**TTAGGGCT |
| EcC02 | 184 | 21 | No | CGGCTA**TAACAT**AGTGGGAGGGTCAC |
| EcC04 | 144 | 21 | No | GAGGATCTGGC**AGATGT**TTGGGTTTT |
| EcC07 | 174 | 21 | No | GGAATTAATTTTTG**TAGCAT**AATGTA |
| EcC11 | 102 | 21 | No | GCACTTTTT**AGGGCT**GGTTATCGCCT |
| EcF11 | 11 | 21 | No | CTGCTTTTA**TGGAGT**GTATGAATGAT |
| EcH05 | 128 | 21 | No | CGATATCACCTCGAA**CGGCAT**CGGGG |
| EcK04 | 90 | 21 | No | TAGGTCG**TTACAT**GTGCATCTTGAGT |
| EcP01 | 90 | 21 | No | AACTGTTTTCGT**TAAAGT**GTATGTAG |
| EcE12 | 153 | 20 | No | AGGTGACGT**CTGTCT**GGCATGGGGTA |
| EcF01 | 166 | 20 | No | TTTGGTCACATGT**GAACAT**GTGTTAG |
| EcF02 | 150 | 20 | No | TTCTTTGTTT**GGGTAT**ATGGGTGCTC |
| EcH05 | 173 | 20 | No | TACGGCAAC**CTATGT**TAACTAGATTT |
| EcP12 | 105 | 20 | Yes | GTGGGTAAGAT**CGAGTT**GTACAGGGG |
| EcA05 | 185 | 19 | No | CTCTAGGT**CAAAGT**CGGAGTGTTTAT |
| EcA12 | 189 | 19 | No | GAGTGGGTAA**AGGAGT**GAGTGTATGA |
| EcC11 | 91 | 19 | No | TCTGTAAGGGAG**CACTTT**TTAGGGCT |
| EcH02 | 63 | 19 | No | CGGTTTG**GGATTT**GTTGGGAACGGCT |
| EcB11 | 97 | 18 | No | AGGGAG**CACTTT**TTAGGGCTGGTTAT |
| EcG10 | 149 | 18 | Yes | TCATTG**TTGTGT**TATCGTGTCATTCG |
| EcL01 | 178 | 18 | No | CTATTTGTAGGC**AAGGAT**AGTGGGAG |

Table 8: continued from the previous page.

| **Clone**[1] | **Position**[2] | **# Starts**[3] | **Shifted**[4] | **DNA Sequence Region, +1 to +25 (5′→3′)**[5] |
|---|---|---|---|---|
| EcP11 | 174 | 18 | No | TTCTGTTGTA**GGAAAT**CCGCTAAAGG |
| EcA02 | 121 | 17 | No | CGTACAAGA**GGAGAT**ACCCGATGATG |
| EcB01 | 146 | 17 | No | TCTCAAA**TGACTT**TTCGCATTCACAG |
| EcC07 | 189 | 17 | No | AGCATAATGTA**GGAGCT**CGATCATGA |
| EcF03 | 93 | 17 | No | TTGTAATAA**TTGCAT**CTCTCTGGACG |
| EcG10 | 178 | 17 | No | ATGAAC**GTAGAT**CGTTGACTGAGGAG |
| EcJ12 | 93 | 17 | No | CGTTG**TACTCT**AAACGGTTTGCATCT |
| EcN12 | 155 | 17 | No | GTGCGGGATGA**GTACGT**CTTAGGTTG |
| EcB01 | 104 | 16 | No | GCTAGT**TAGTGT**TTGGTGTAGTCGTA |
| EcB09 | 14 | 16 | No | GAATTGCGCC**AAATAT**GTCCGTATTT |
| EcG01 | 166 | 16 | No | GATGTTGCTTG**GTCCAT**GGATGAGTT |
| EcA01 | 93 | 15 | No | GAGTCCATTTTA**GGGAGT**ATCTCCAT |
| EcC07 | 187 | 15 | No | GTAGCA**TAATGT**AGGAGCTCGATCAT |
| EcC08 | 170 | 15 | No | TGAGTTTCTT**GTATAT**CTGAATCGAT |
| EcD01 | 182 | 15 | No | TGAATAGTGGT**TTAGAT**GGAGTCGTG |
| EcD04 | 55 | 15 | No | TCATCATCTTTG**CGGCCT**TCACAATG |
| EcE07 | 177 | 15 | No | TTGGACATT**GGATAT**CCACGTGGAGG |
| EcF03 | 152 | 15 | No | TTTGCT**GTGCCT**TCTTTTTCGGAGGA |
| EcI12 | 76 | 15 | No | TACTGATTCG**TTGGTT**GATTTGATAG |
| EcN12 | 114 | 15 | No | CGGATTGGCT**CGCAGT**GTGCGGGCGA |
| EcP11 | 135 | 15 | No | TGGGAGTGTC**AGGTGT**GATACAGTAG |
| EcF08 | 150 | 14 | No | ACAAACTTAG**GGAGCT**TACCTGATGT |
| EcF12 | 153 | 14 | No | TTGGCGTG**CTATCT**TCCTCATCAGCT |
| EcH02 | 130 | 14 | No | CCGGTCTAGGC**TGATCT**GAATATGGT |
| EcI04 | 191 | 14 | No | -TCGCTGTTGGAG**GGGTAT**AATGATG |
| EcI07 | 113 | 14 | No | CGAATAGAGGGA**CAGTGT**TCGATATC |
| EcL04 | 81 | 14 | No | TTCGTTCAGT**TTATGT**TCTCAATTTT |
| EcO07 | 181 | 14 | No | TTGGTCTAGTT**GTGGTT**TAGGAGTTA |
| EcP09 | 169 | 14 | No | GAGTTTCGC**TACTGT**TGACGTTGCTG |
| EcB04 | 171 | 13 | No | TCAAGCTTGATAACCCAGGTGTACTG |
| EcC02 | 182 | 13 | No | TTCGGCTA**TAACAT**AGTGGGAGGGTC |
| EcE04 | 188 | 13 | No | TTGTACTCTGAAGGAGACGATTGATG |
| EcE07 | 189 | 13 | No | TATCCACGT**GGAGGT**CGAAGATGATG |
| EcE07 | 169 | 13 | No | TGGTAGTGTT**GGACAT**TGGATATCCA |
| EcF12 | 176 | 13 | No | GCTCGTCATAGTT**CAAGTT**TGGGAGG |
| EcG08 | 145 | 13 | No | TGGGAAGT**CACTGT**GGTTGTGGTGGC |
| EcJ09 | 126 | 13 | No | TACCAG**TACTGT**TACAAGTGTCATAC |
| EcM03 | 99 | 13 | No | GGTGCGGTT**CTATGT**GTACGAATCGT |
| EcN10 | 112 | 13 | No | TAGGTTGTTGTG**AAAGAT**GTTATTTT |
| EcN12 | 153 | 13 | No | TGGTG**CGGGAT**GAGTACGTCTTAGGT |
| EcP08 | 131 | 13 | No | TTTTCA**CGAGAT**CGATTTCTTTCAAC |

[1] Clone: The ID of the identified *E. coli* clone.

[2] Position: Position of the TSS, within the ArtPromU sequence.

[3] Starts: Number of mapped RNAseq reads starting at the stated position.

[4] Shifted: Was the TSS shifted one position to the left (in case of neighbouring read stacks of similar height)?

[5] The part of the artificial 5′-UTR sequence spanning from +1 (TSS) to +25. The identified motifs (by Improbizer[20]) are underlined and coloured in grey. Darker colour represent stronger matches to the identified motif, GGATGT.

##### 1.4.8 Determination of relative presence of clones with functional ArtPromU sequences.

To determine what fraction of a GeneEE DNA library is harbouring clones with functional ArtPromU sequences, we have created two libraries which were based on the plasmid pRL101 (Table 4), that carries the antibiotic marker kanamycin on the backbone along with the *β-lactamase* gene that confers ampicillin resistance. pRL101 was PCR amplified by the primer pair BsaI-Bla-F and BsaI-Bla-R amplifying the entire plasmid excluding the native *β-lactamase* promoter. Both the PCR product and the GeneEE(±SD)BsaI inserts were digested with BsaI and DpnI overnight. Both the vector and insert were purified and the vector was CIP-treated (NEB) at 37 °C for 1 hour and purified. The ligation mix was made using 1:10 molar vector:insert ratio in a total volume of 30 µL using the T4 Quick Ligase (NEB). Ten µL of three aliquots were then transferred into the cloning host *E. coli* via chemical transformation. Both libraries along with a no insert control were plated out both on LA plates supplemented with 50 µg/mL kanamycin, and LA plates supplemented with 50, 500, and 1000 µg/mL ampicillin. The observed number of resistant colonies are listed in Table 9.

Table 9: The relative presence of *E. coli* clones with ArtPromU sequences within a GeneEE DNA library.

| | **Antibiotic markers and concentrations (µg/mL)** | | | |
|---|---|---|---|---|
| | **Km (50)** | **Ap (50)** | **Ap (500)** | **Ap (1000)** |
| **pRL101+GeneEE(−SD)** | | | | |
| Biological replica 1 | 74067 | 27222 | 4056 | 2078 |
| Biological replica 2 | 59800 | 27333 | 2756 | 856 |
| Biological replica 3 | 50711 | 23711 | 2533 | 1156 |
| Average[1] | 61526 | 26089 | 3115 | 1363 |
| Ratio[2] | 1.00 | 0.42 | 0.05 | 0.02 |
| **pRL101+GeneEE(+SD)** | | | | |
| Biological replica 1 | 24444 | 6522 | 1633 | 533 |
| Biological replica 2 | 29878 | 11700 | 1267 | 411 |
| Biological replica 3 | 56978 | 19244 | 1822 | 889 |
| Average | 37100 | 12489 | 1574 | 611 |
| Ratio | 1.00 | 0.34 | 0.04 | 0.02 |
| **pRL101 (No insert control)** | | | | |
| Biological replica 1 | 1633 | 111 | 56 | 0 |

[1] The average was calculated based on three biological replicas.

[2] The averages were normalised by setting the number of colonies identified in Km (50) plates to 1.

##### 1.4.9 The construction of GeneEE plasmid DNA libraries for the functional screening for inducible gene expression in *E. coli.*

For this construction, the *xylS* gene with its native promoter was PCR amplified from pHH100 by the primer pair XylS-EcoRI-F and XylS-XbaI-F, introducing the recognition sequences for EcoRI and XbaI, respectively, (Table 1). EcoRI and XbaI digested vector, pRL101, and the insert carrying the *xylS*, were then ligated using 1:3 molar vector:insert ratio in a total volume of 20 µL using the T4 Quick Ligase (NEB). The mixture was then transferred into the cloning host *E. coli* via chemical transformation and grown on LA plates supplemented with 50 µg/mL kanamycin. The *xylS* coding sequence carries a BsaI recognition sequence and in order to eliminate this site the entire plasmid was amplified by the primer pair XylS-delBsaI-F and XylS-delBsaI-R (both primers carry a single nucleotide point mutation, hence eliminating the BsaI recognition sequence), by following the Overlap Extension PCR cloning method[9], and the resulting linear PCR product was transformed to *E. coli* via chemical transformation and grown on LA plates supplemented with 50 µg/mL kanamycin. This resulting plasmid, pJN101, was then PCR amplified by the primer pair BsaI-Bla-F and BsaI-Bla-R amplifying the entire plasmid excluding the native *β-lactamase* promoter. Both the PCR product and the GeneEE(–SD)BsaI were digested with BsaI and used in cloning as a vector and insert, respectively. The ligation mix was made using 1:5 molar vector:insert ratio in a total volume of 20 µL using the T4 Quick Ligase (NEB). The mixture was then transferred into the cloning host *E. coli* via chemical transformation grown on LA plates supplemented with 50 µg/mL kanamycin. The resulting library was consisting of ∼30,000 transformants.

###### 1.4.10 The functional screening for artificial inducible gene expression in *E. coli.*

For the functional screening for clones with inducible gene expression phenotpye, the library, consisting of 30,000 clones, was plated out on two sets of LA plates, one set with (1 mM *m*-toluic acid) and another set without inducer. Each set was consisting of nine LA plates supplemented with 2, 3, 4, 5, 6, 7, 8, 9 and 10 mg/mL ampicillin. A 96-well microtiter plate, containing LB supplemented with 50 µg/mL kanamycin, was inoculated by picking random colonies grown on plates with inducer and ampicillin concentrations ranging from 6 to 10 mg/mL. The plate was incubated overnight at 37 °C under 800 rpm constant agitation. The cells were then replica plated using a 96-pin replicator to two sets of plates, with (1 mM *m*-toluic acid) and without inducer, to 18 plates each plate supplemented with 2, 3, 4, 5, 6, 7, 8, 9 or 10 mg/mL ampicillin. In total 27 clones were found to be showing an inducible phenotype and all 27 clones were Sanger sequenced (GATC Biotech, Germany) using the primer pRL101-Bla-Seq (Table 1). This led to seven unique constructs A5, A7, C12, H3, C4, F2 and F11 (Table 10). In order to confirm the XylS-dependent induction the *xylS* coding sequence with its native promoter was deleted from the corresponding constructs by digesting with XbaI and NdeI, blunting the linear DNA with T4 DNA polymerase (NEB) and ligated using the T4 Quick Ligase (NEB). The mixture was then transferred into the cloning host *E. coli* via chemical transformation and grown on LA plates supplemented with 50 µg/mL kanamycin. As for the final confirmation, all 16 clones were grown in a 96-well microtiter plate containing LA supplemented with 50 µg/mL kanamycin and after overnight growth they were replica plated onto two sets of plates with (1 mM *m*-Toluic acid) and without the inducer, each set with LB plates supplemented with ampicillin concentrations ranging from 2 to 10 mg/mL (Table 11).

Table 10: The ArtPromU sequences that lead to inducible phenotype in *E. coli*.

| Clone | DNA Sequence (5′→3′)[1] |
|---|---|
| A5 | GGTATATTTCGTGAGTTGTTCGGTCAAATACTGAACCTTTTTGACTTCGCTTAAATTAACGATGTATTA |
| | ATACACCGTGTTTGAGTTGTTAGGTGTGTTGTGACAGAATGACCTGTACTAGCGGGAGGACACGTAAAT |
| | TTTGGTCCCGTTATAACTTATCATGCCATTCCTGGCCGAACGAACTATGAATTCTATGAAAATG |
| A7 | AGATGTAAAGGGACGTCAGCCATCACTTCCTTCTCGATGGCAGAAATCATTTTGCAGTGTACCTGAAGT |
| | GTCGATTTCTTCTTTAAAATTGTGTTTTAGTTGATGCAGCTATAATTTATCCGTAACGGATGATGCGGG |
| | GAATTATTTATGGTACCTTGCTTCACTAAGTGATTTGTATCCTAACTTGTATATATAACTTATG |
| C12 | TTCCAAAATGTAGTATGACTAAATTAGATGATTACTCCAGAAGTTTTTCAAGGTAGCTTTTTAGTGAGT |
| | TTGATTTTGTTAGTAGGCTGTGTATCCTGCTGGACAGATCGAGAAAATTGTCGTAAAAAGTCTGTCGTC |
| | TCTGTTAATGTGTTCTCAGGTGTCCTGTTTATATCGTTGTGATGATCAGAAGTCTATTAGGATG |
| H3 | CATCAACTATATTGCTTACGGCGATTCTCTGTGCTCGAATCTAGTGTAATTGTTTTCGTATTTTACTAA |
| | CTTCCGATGTGCACTTATAATTCTGTCAATTCATGGTTATTTCGGCATATATGACCAGTCATTGATGTT |
| | ATCGAGGGCCGTCGTATAAACGGGGAGATCTAAGAGTCGTAGGCTTATGAGGTGTTTCATAATG |
| C4 | GGGCAACCTCAGGTATTACATGCGGCTCAGGTTGTACTTGATTAGACTTCTTGTCCTCGTTATGATGGT |
| | GCCTCTTTCGTGTTATTGCGCTTGGGACGCTGTTATGAGAAACTCATATGTGTATCATCAGCCTCGGCG |
| | GCAATATTCCAAGTATGCTTTTAGGTATTGATTCTCAGGCTTGTGTGATAAATGTTTGCAGATG |
| F2 | CTCATTTAGTTTAATAGCGCATTTTCGTGTAACGTATGATAGGTCAGCTACTTTTATTTATTGGTGAGC |
| | CTATTATGTAATCGTCCATTAGAACGGGGCCCGTCAGATCATTAGCAGCCTAATTTTGTGGATTTCTTT |
| | CGGGCTTTGTCGTATTATCTAACCCAATTTAGCTTTGCGGTTGTTGTGGGGAGCTCGTAGGATG |
| F11 | AAAGAACTTGAGTAATAGGGAAGAATTTTTGTAAAATGTATGAAAGAATTATGATGGATCTATTTCTAG |
| | GACGTCCGGTACGGATGGTAGTAAAAATTCAGATTCTTGGTTAAGGCTGTTTGTTTCCGGGACCTCAAC |
| | TGTCATGGGGCCACCCCCGGCGTTACTATTCGTTCTATAAGTCAACTTCTTAATATGCATGATG |

[1] The ATG sequence at the 3′-end of each ArtPromU sequence is the start codon of the *β-lactamase* gene.

#### 1.5 *Pseudomonas putida*

##### 1.5.1 Growth conditions.

*P. putida* cells were grown in LA or LB at 30 °C, supplemented with 50 µg/mL kanamycin when selection was required as described in the text.

##### 1.5.2 The functional screening of GeneEE plasmid DNA libraries in *P. putida*.

For the selection of functional GeneEE sequences in *P. putida*, using mCherry as the reporter, the same library that was established in *E. coli*, described in subsection 1.4, was used. A volume of 10 mL of LB was inoculated with 100 µL of cells, from the frozen culture harbouring the GeneEE plasmid DNA libraries, and was grown for 2 hours at 37 °C, and the plasmids were purified with QIAprep Spin Miniprep Kit (Qiagen) and quantified with Nanodrop (ThermoFisher). One µL of plasmid (25 ng/µL) was electroporated (2.5 kV for 2 mm gap cuvette; 200 Ω, 25 µF) to electrocompetent *P. putida* with the Gene Pulser Xcell Electroporation System (BioRad) using a previously established method[21]. The electroporated cells were then added to pre-warmed 1 mL of LB and were incubated for 1.5 hours at 30 °C. The cells afterwards were plated on LA supplemented with 50 µg/mL kanamycin. Among the obtained colonies (Figure 5) several colonies with visibly red colour to the naked eye were picked and plasmid DNA was isolated with QIAprep Spin Miniprep Kit (Qiagen). These plasmids were then Sanger sequenced (GATC Biotech, Germany) using the sequencing primer pLT101-mCherry-Seq (Table 1). The identified ArtPromU sequences with their corresponding phenotypes (mCherry read outs) are listed in Table 12.

Table 11: The ampicillin resistance phenotypes of the clones with inducible phenotype identified in *E. coli*.

| | **Plasmids with XylS**[1] | | **Plasmids without XylS**[2] | |
|---|---|---|---|---|
| | − inducer[3] | + inducer[4] | − inducer | + inducer |
| **Clones** | **Ap resistance, mg/mL**[5] | | | |
| A5 | 2 | 4 | 2 | 4 |
| A7 | 2 | 4 | 2 | 4 |
| C12 | 2 | 4 | 2 | 4 |
| H3 | 2 | 4 | 2 | 4 |
| C4 | NG[6] | 4 | NG | NG |
| F2 | 2 | 6 | NG | 2 |
| F11 | 2 | 6 | NG | 2 |
| Control[7] | NG | NG | NG | NG |

[1] Indicates the condition where the *xyls* gene is present in the plasmids.
[2] Indicates the condition where the *xyls* gene is absent from the plasmids.
[3] In the absence of inducer.
[4] In the presence of inducer, 1 mM *m*-toluic acid.
[5] The numbers indicate the highest ampicillin concentrations at which growth was observed. The range of concentrations used were 2, 3, 4, 5, 6, 7, 8, 9 and 10 mg/mL.
[6] No growth.
[7] The control construct is the re-circularised plasmid with no insert.

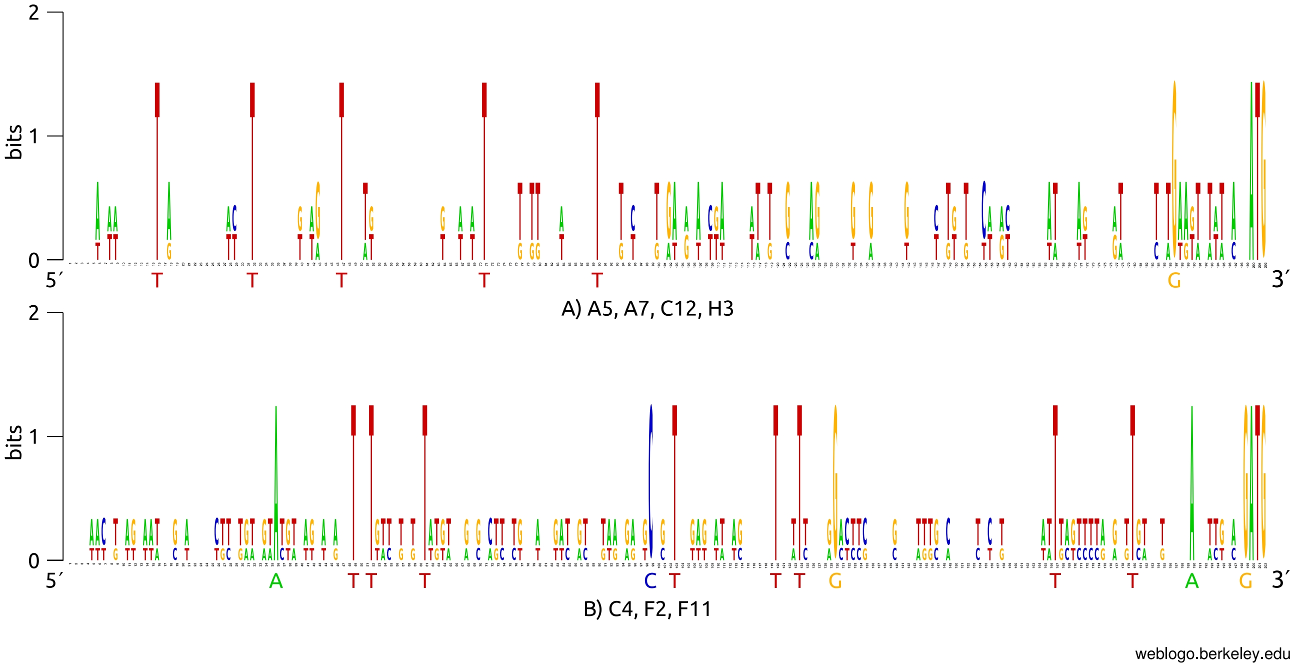

Figure 4: The WebLogo image of ArtPromU sequences that led to inducible phenotype in *E. coli*. The multiple alignment of the ArtPromU sequences of the clones A5, A7, C12 and H3 is depicted in part A; and the clones C4, F2 and F11 are in part B. For clarity, the conserved nucleotides are depicted in larger font size below each part.

Table 12: The ArtPromU sequences identified in *P. putida*.

| **Clone** | **FIM**[1] | **DNA Sequence (5′→3′)**[2] |
|---|---|---|
| PpL | 13 | AAATCCTTCATCGTCAATGGATGTTCGGATGCGCGGGCAGCTCCGGAGATGTAGAATG |

Table 12: continued from the previous page.

| **Clone** | **FIM**[1] | **DNA Sequence (5′→3′)**[2] |
|---|---|---|
| Pp06 | 19 | TAGCAATATTCTATTTTGTGTTGCGTATTTAATTTAATGCAGTTACGTTCCCCGCTTGGGATTT<br>GGCCTTTTGTACATTCTGTGGTTTTTGCTGAGGATTTGTTGCGTGCTATGTAACGAGGGTGGTA<br>CTCCAAATTGTTGTACTAAGGGGTTTCTTGGTATGTAATGGGAGGTTGTATTTATTCTTTGTAT<br>TTTGAAGGGAGCTCTATAATG |
| Pp01 | 20 | ACAGGAAATGGGAGTTAGGATCACCCAGCGAATTGTGATCATTATTGCTTAGATGTATGGTATA<br>GAATTTCTTGTTTGTTTTTTACCTGTGTGGTTAAAATGCCAGGAAAGGTTGAAATGAGTCAGCA<br>CTGACCCCATTGATAACTGTCTGTCGATCGGATACCGTTGACTGGCTGCCTTCTAGTTCGGCTG<br>CGTTGCGAGGAGACGACACATG |
| PpM | 20 | TTTGTGTTAAGGACTTTGTTGGAAGAAAATGCGGGTTATTGTGTTGGTTAATTGTTGTGATCGG<br>GCGAGTTCAACTTTCGTCACACCGTCTTTATTATTAATTTTATTGTCTCCCTCATGCATAACTG<br>GCTGATCGTTGTATGTCCGTTACTCAATTGGGAGATTTAGTATG |
| Pp09 | 21 | TTTTATTAGATTTTCTGTGCTTCCAAATGGTCCCGTTTTAATTTTTCATTTCAGTATGGTCAGA<br>TTATGTATAATGTTGTTCTCGTAGCGGTTAATTACGAGGCCGGGGGAGATGGTAGTACGCAGTC<br>GCGCTGTGGTGGGTTGAGATGGGTTAGCATTCACGGCCTGGCGTTCTATAATTTTTACAATTTG<br>CCGCCAATGGAGCGGTATAATG |
| Pp11 | 30 | AAGCTGGGTTGCCGGTACCTCACTGGGCTGTGACGGAGATGTGGTGGTGTCGGTTGGGCTGGTG<br>CTCTCTTTCGATGCCCTAGGGAGGTCGCTGCTATTATGCCCGACATGTAGATGTGGGAGATATG<br>CATGTCACGGAGGACATCGGCATGGAAGGCCGCCTTAGGCGAGTTGTGTAGTGCGTAGTCGCTT<br>AGGCCTGTGGAGTCTTATGATG |
| Pp07 | 34 | TAATTCTTGCGTGAAAGAGTTACCTACCCACTCTCTTCTCAATTCTGCGCAGAGGGCTAATGTT<br>AACTGTCGGATTACTTAAAATTTTTTTACATAAACTCTTCTCACTACGTTAGATTATTTAGGT<br>TTATGGAACCGGGAGCAATTCTGTTCTGGCGTGTTTCTCAGTCGGCTAAGGCTTATTCGCGGGA<br>GGTGAATCATG |
| Pp04 | 50 | CGCATTTTAAGCTTCTATAGGCTGGTGGAGCTCCTGCGGCTCCATGTTGTATTATTGTGGAAAA<br>TTGTCTGAACTGACGGTAAGTTTTCATGTTACTGTCGTTCGGGGTATGCCCCTATTATATGATT<br>TATGGGTAGAACTAGTATTTCCAGCGATTTTACCTCTTACTGGGCCACGTATGCTATTGTGGAG<br>GCTAAGTATG |
| Pp08 | 56 | AAAAAGTAATTTGTAAAAAAGCAGACTCACGGTTGCGATTGCACCCAGGGTGTTCGGGGCGGTT<br>TCTTCTTGGTCATCTTATATTGAGGGGGGCTGTCTTGATTTATATAGTTTTACTGTCCTCCGAT<br>CTCCTTTCTCAAATAGTTCGAAGTGTATGGTTTACTTGTTTGTTATCATGGGACGGGTCTGAGT<br>GAGGAAATGGAGAACGACAATG |
| PpH | 86 | AATGAGTGATTCTCAATAAGATTTAACGTTTTATGTCGTTTTTTGTAGAAGATGCGATCATTAA<br>ACATAGTGGTTTTCGAAGATAGGTTAGCTTGGTTGTGTTATGCTGACGTTGCTGACCAGTTGGG<br>AGCGGTATGCCAGGCTGTGTGGCGCCTCAATCCTAAGGTCTATCGGGTGAGGCGCTTCATTCAT<br>GATGGTAAGGAGAAACTAGATG |
| Pp03 | 100 | GGGCTAAGGGTAATTCTTCTGGATTTAAAAAATGCCTTTTTATTGTCTGGTTCTGTGGGAACAG<br>CCGATGCGGTCTGGTTGTTCTAATTGCGCGTCGTTTCAGCCTATGTGCTAGGGTCTGTCCTAGT<br>GCTTCCAAACAATGGTATTCTCTCATTTGGCACTAATGGATCTGTGATTAGTACGCGGTATGTG<br>GGCTTAGGGAGCAATATAATG |

[1] Normalised fluorescent intensity measurement read outs.

[2] The ATG sequence at the 3′-end of each ArtPromU sequence is the start codon of the *mCherry* gene.

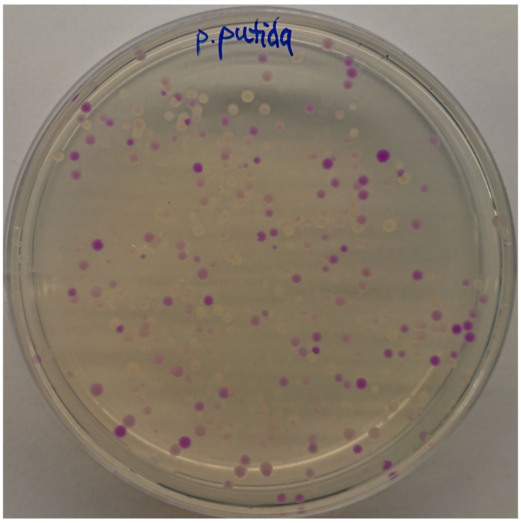

Figure 5: *P. putida* cells expressing mCherry at varying levels.

#### 1.6 *Thermus thermophilus*

##### 1.6.1 Growth conditions

*T. thermophilus* cells were grown in Thermus Broth (TB, 8 g/L bactotryptone, 4 g/L yeast extract, 3 g/L NaCl, all dissolved in mineral water) or Thermus Agar (TA, TB + 15 g/L agar). A range of kanamycin (30, 60, 90 µg/mL) were added to the growth medium when selection was required as described in the text. TA plates were incubated in a moisturised plastic container to avoid agar plates from drying out.

Stock pellet of *T. thermophilus* from –20 °C was thawed at room temperature and was inoculated to 1 mL of TB and subsequently was transferred to a 125 mL flask containing 20 mL of TB. The strain was cultivated under constant mild agitation (150 rpm) at 65 °C.

##### 1.6.2 The construction of GeneEE plasmid DNA libraries in *E. coli*.

For the construction of GeneEE plasmid DNA libraries in *E. coli*, an *E.coli*-*T. thermophilus* shuttle vector, pMK184, was used (Table 4). The pMK184 plasmid, providing the thermostable kanamycin resistance as the reporter, was PCR amplified using the primer pair $Km^T$-F and $Km^T$-R (Table 1) excluding the native PslpA promoter controlling the expression of the thermostable kanamycin marker. The reverse primer $Km^T$-R carries the BioBrick Prefix[1] as a flanking sequence that serves as a homologous sequence in Gibson cloning. This amplicon was used as the backbone and the GeneEE(+SD)$Km^T$ as the insert. The Gibson ligation mixture was made using equimolar amounts of purified vector and insert (totalling 100 ng DNA) in a total volume of 5 µL, added to a volume of 15 µL of Gibson assembly mix. The mixture was then transferred into the cloning host *E. coli* via chemical transformation and grown on LA plates supplemented with 50 µg/mL kanamycin. The resulting library was consisting of ∼1,500 transformants which were scraped of the plate, followed by plasmid isolation with QIAprep Spin Miniprep Kit (Qiagen) and quantification by Nanodrop (ThermoFisher).

##### 1.6.3 The functional screening of GeneEE plasmid DNA libraries in *T. thermophilus*.

For the functional screening of the GeneEE plasmid DNA libraries, the purified plasmids from the *E.coli* transformants were transformed, using natural competence, to *T. thermophilus* as previously described[10]. A fresh pre-warmed TB was re-inoculated with an overnight culture of *T. thermophilus* in a 1:50 dilution ratio. The strain was cultivated in a 65 °C shaking incubator until an $OD_{550}$ nm of 0.4 was reached. From this culture 0.8 mL of the culture was aliquoted to a new 12 mL tube and 200 ng of the GeneEE plasmid DNA libraries isolated from the *E. coli* clones were added. The mixture was incubated for four hours at 65 °C and afterwards was plated on selective TA supplemented with 30, 60 and 90 µg/mL kanamycin. The plates were incubated overnight at 65 °C. Several clones were randomly picked from the TA plates and plasmids were isolated with QIAprep Spin Miniprep Kit (Qiagen) and were Sanger sequenced (GATC Biotech, Germany) using the sequencing primer pMK184-Km$^{T}$-seq (Table 1). The ArtPromU sequences with their corresponding phenotypes (kanamycin resistance levels) are listed in Table 13.

Table 13: The ArtPromU sequences identified in *T. thermophilus*.

| **Clone** | **Km**[1] | **DNA Sequence (5′→3′)**[2] |
|---|---|---|
| Tt01 | 60 | ACAGCGTTTGCAATGGTAACTCGCGTTTATGCTATACTTGAAAGCGAACTAGCATTAGAACTCTA<br>AAAGGTGATTGTAAGTGACGATGGAGTGTTAATCTTAGGGGGGGGACGCGATTAGATTAGGCGCA<br>TCCGTCTAGTTAAGTGTTGGGCCGTCAGAGGACGGCAAGAACTTCGGTGGTTAGATTGGTCATGT<br>GAGCAGGAGTCATGCAATG |
| Tt02 | 90 | ACTTCTGGCGTACTTTCATGCAGTCAGAATACGATCCTTGGAAAACTACATGCTTAATAAAGGGA<br>GAAAAGGGTCATGTTCTTCGGAAAGGTCGAACATTGTTGATGTCGATTGGTGACTTGTTGTTTTG<br>ATTTATGATAGGTTGTATGGGTCGTGGGCTGAGAAGGCTCTAGACTTGTCGAGTAGTACGGTCAT<br>GAGGGGGAGCGTGTATATG |
| Tt03 | 60 | GttGGTCAGGTATACTTTGCTTAACGTTTGGTTTATTTTCTTTATTTTTTAAGATGTCCTAGGTT<br>AAATTATTCTGCTTTTTCTTGCGGATTAGGGCGTTGCTCGTTTAGGATAGTAATTTTATAATGAG<br>GTTGTGCGTAGCGGCCATGATGTGTTCCATGTATGACACTAGAATGGGGTGAGCGGGCCTGTTAT<br>GGTGCGGAGTGATCTTATG |
| Tt04 | 90 | TTACCATTTCAGCCGCGGTCTTTTATAATTCGGTAGCGGTGCGTTACGGGGGTTAAGGTGACGGT<br>CGTCACGTGGTTGGAGACACGGTTCGGGATTGTGTTCTATCATTGATGGGGGTAGAAGGGAGTAC<br>ATTTAGTGGTCGTCCTTTGGCTGCCTGGTTTGAGGGAAAATAATTGCTGACTGCTGTAGCGGGAA<br>AGGCCGGAGGGGCTGGATG |
| Tt05 | 60 | AGGTGGGGTACGCTGTTTTTATACTTTGTTGGTCTTTATTGTGTTATAGTGATCGGAAGATAGGA<br>AAAATAGTTGTGTGATATGCTGGGATTTGTGTGGTGGTTGTCTAGCAGGCTTTGCAGTTAGACGT<br>GGACATTTGGTGGGGGGGCGTACCGGGTCAGGGTTGTGTGGTAAGTAATTGCGGGCTTAAATTCG<br>GTTGTGGAGAGGGTGAATG |
| Tt06 | 90 | ATATTATGGAGTGTTGAGTACGCTTCACCAGGTGGGGGCGTGGTATCATGCGAAGATGTTAGTTT<br>TTGCATTCTTCTGAAAGGGTCGATAGAAGTATACTTTATGTTTAGGGGATGGATATGACCGGTTT<br>TTTCCGTAGGTAATTAGTTAACGCTGTTCGAGTTTTAGCGGGTAGTCTGAGTGGCATGTACGGGG<br>CCTTTGGAGGTGGGTGATG |
| Tt07 | 60 | GAACGTGTATCTCTTTTTGTGATAGGATTGGTCTTGCTATTGACGTTAACGTGGTGATTGTTAGA<br>CTGTTACTCTGGGTGTCTAAATTGTTGTCTACAAATTCATCGGTATCGAAAGTTAGCAGTGTGTC<br>GTTTGTTTTATAACGGTGACGTTGCATATTTCGTTGCGCTCGGCGATAGTTGATTCTTTTCGTAG<br>CGCTAGGAGGACGAGGATG |
| Tt08 | 60 | TTGTTATTGTTGATTTTGGAATGCTCTGTGTAGTATCTGGCTTTTCCTGTTTTAGTTTTGGGGTA<br>ATATGTTATACATGATGCGATCAGTCACTATTCGGGTAACTGTTATGGGTTATGGGTTCTTTGAT<br>AATGATGTGAGTGGAGCTTCAGGGAAGACTAAATAAGGCGATGGGATCCTGCGAACTAAGATTGC<br>ATAGGAGGGTTAGGATG |
| Tt09 | 90 | ATTGGATGACCATCTATTCATTTCACTTGTCTTACGGCAGGGTTGCTGTTCTTCTGGTTTGCTTA<br>ATTTGATGCTTTTGGCTCTAGCCCTATAGGGATCCTGACATTGTTGTATAATGTTAATAGTAGGG<br>TATAGTTGTCTCTAGTGTATGCTAACGGTGGGATTATTCGGATGGCCGTGCGGGACTGTCGGGTT<br>ACGGAGGAGTGGCAGAATG |
| Tt10 | 60 | GCTGCGATTATGGTATATTGAATTCTGGTCATAGCAGTAGCGTGAGTATGTAGAGCTCCCGTTGT<br>GGAGTACTGGAGGAGAGACGTTATG |

Table 13: continued from the previous page.

| | | |
|---|---|---|
| Tt11 | 90 | TCTTGAATATTAGTGTACAATATTGGTTTGTAGTGCTCAGGTTTGTTAGTGTGGTTGCTCGTGGG GAGTGCGCGTATG |

[1] Kanamycin concentrations used for the selection (µg/mL).

[2] The ATG sequence at the 3′-end of each ArtPromU sequence is the start codon of the thermostable *kanamycin* gene.

#### 1.7 *Corynebacterium glutamicum*

##### 1.7.1 Growth conditions.

*C. glutamicum* cells were grown in Brain-Heart Infusion Broth (BHIB, 37 g/L Brain-Heart Infusion mix [Difco], 91 g/L sorbitol) or Brain-Heart Infusion Agar (BHIA, 37 g/L Brain-Heart Infusion mix [Difco], 10 g/L agar) at 30 °C, supplemented with 15 µg/mL chlorampenicol when selection was required as described in the text.

##### 1.7.2 The construction of GeneEE plasmid DNA libraries in *E. coli*.

For the construction of the GeneEE plasmid DNA libraries the pXMJ19 plasmid was amplified by the primer pair pXMJ19-Chl-F and pXMJ19-Chl-R (Table 1) leading to the amplification of the entire pXMJ19 plasmid excluding the native *chloramphenicol* promoter. This primer pair also introduces BsaI restriction sites within the overhangs used for the BsaI-based restriction cloning. After the PCR, the PCR product was digested with DpnI and BsaI at 37 °C for 3 hrs and purified with QiaQuick PCR purification kit (Qiagen). The GeneEE(–SD) segment was used as insert and ligated with the pXMJ19 plasmid amplicon in a 1:7 vector:insert ratio, overnight at 16 °C with T4 DNA ligase (NEB). The ligation mixture was first heat-inactivated at 65 °C for 20 min, and then transformed to chemically competent *E.coli* cells, leading to ∼1000 transformants/10 µL. The final library consisted of ∼2000 clones. The resulting transformants were scraped from the agar plates and plasmid DNA was isolated with QIAprep Spin Miniprep Kit (Qiagen).

##### 1.7.3 The functional screening of GenEE plasmid DNA libraries in *C. glutamicum*.

For the functional screening of the GeneEE plasmid DNA libraries, the purified plasmids from the *E.coli* transformants were electroporated to *C. glutamicum*, as described previously[18]. The electroporated cells were plated on LA plates supplemented with 15 µg/mL chloramphenicol, leading to library of ∼200 transformants in *C. glutamicum*. Among the obtained clones, 10 clones were randomly selected and plasmids were isolated with QIAprep Spin Miniprep Kit (Qiagen), and were Sanger sequenced (GATC Biotech, Germany) using the primer pXMJ19-Chl-Seq (Table 1). The identified ArtPromU sequences are listed in Table 14.

Table 14: The ArtPromU sequences identified in *C. glutamicum*.

| Clone | DNA Sequence (5′→3′)[1] |
|---|---|
| Cg01 | TGCCCGGGCTCGAGTTCAAGGATTGCGACAATCTGTTCTGTGGCAGCGTTTGAGCTTATGTGATAATC TGAATTGTAAGTGAGTATCGTAAGGGATTTTCATACTAGTTGAGTTTACAGTAAAGTTAAGTGTGTAT CGCTAGTTAACTAATCGGGACTTGAAATTCTAGCAAGAGTAGTGTCCCGGCACAGAACAAGGAGTGTT ATGCATG |
| Cg02 | AGTGCCCTTAATAAGTCCATTTGATTGTTCCATTATAGTCTAGATACGTCCTGGTATTTTGCAATTAG GATTGGATGTAGTCCGTAGCAGTATTTTTTTATGTGTATAGATTTACAATGAGTGTCGTATATTTTGC AGATGCCTCCGGGATTGGACTGGTATCAATATTCACGGTAGCTTTGAGAATTATGAGTAGGGAAGGAC AAATG |

Table 14: continued from the previous page.

| | |
|---|---|
| Cg03 | AGTGCCGGCCTGACAATGGATGGTATTGGATCTTAATTGCAGTAGCTTTCTGTCTGCCGATTATCAAG |
| | AATTTCTTGACAGGTGCGAAGGTAGTCACCCTGGTGAGGCATAGTAGTTCGGATGCATATTGCAAAGA |
| | GTTTTCGTGGGGAAGTTGCGAATGGTCGTAAGAATGGTTGCTCAACGTCGGTAACTTCTCTACAATGG |
| | GTATGCATG |
| Cg04 | AGTGCCAAAGTCCATGGATTTGTAGTTACGTTTATTGGTTGTTCTCATTGGGTGACTATTAAGATGCG |
| | ATCCATGATGTTTTAGGGCTGGTATGATATCGAGAGGTTTTCATATCATTTCTATGTATGTCCGGTGA |
| | GATCTTTAATATTCTGGGATCATTGATGACTCCAACACTTGATATATTGATACGTTTTAATTAGTATT |
| | CAATGAATG |
| Cg05 | AGCATCAACCCATTGCCACATCCTACTGAAAGGATCACCATTACGTGTTACTACATGGGTCGCTCACC |
| | GCCGACTTCCGCATGACCATACGCAAATTTCTTACCTAGGGCCGTTCGATCATACTTGCAATGCACCC |
| | AATATGCAGTAGAATCCGATACATCAAAACACCAATTATCAATACTACATAAGGACACCAAGGATTCG |
| | AGGCAAATG |
| Cg06 | AGCATTACCTGTGAACCTTCAAAAGCGTGACTTGCTTCCGTGAAAGGCATGAGTCACTAGTTTAGTGC |
| | GATGAAGTCACGCCTGTGTTAATATACATAAATACCTAACATTATCGCAATTTACTAAACCCCCCCGT |
| | AATAAACTCTGTAGTATGATGGAGTTGCACAGATTAATACAAAAAGATAAACACAGATCTGAAAATGA |
| | GGCAGATG |
| Cg07 | AGCATGGAGAAAAAAATCACTGGAAAACGTTCTTCGGGGCGGTCTCATGCCAGTGCCGGTTTTGAAAG |
| | GGTTCATAGATTTGTGTAGTTACCTTGGAAAACAGTCGGGCGAACATGCATAATAAGTTTGCCTATAT |
| | ATTGTGACTTGCCAGTACCTGGTTCGAGATGGAGATGTGTTGTTTTATATAAGGAGCATAACGTGTTA |
| | CGATAGTCTCCGTTGAAGAACACTTTAATTATGTTTGTATGCTGATGATGAGGATGTATG |
| Cg08 | AGCATTACCTGTGAACCTTCAAAAGCGTGACTTGCTTCCGTGAAAGGCATGAGTCACTAGTTTAGTGC |
| | GATGAAGTCACGCCTGTGTTAATATACATAAATACCTAACATTATCGCAATTTACTAAACCCCCCCGT |
| | AATAAACTCTGTAGTATGATGGAGTTGCACAGATTAATACAAAAAGATAAACACAGATCTGAAAATGA |
| | GGCAGATG |
| Cg09 | TGCCTGCAGATATAGAGGAAATGGAACGTAAAGATTGATATGGGAATAGGCTTCATATGAATAAGTGC |
| | TCAGTAATGTTCCCTAATGCCTCGCTCGAAATTGCCTAGTTATTCCTACGGGGCTTTTGCTTCTATTT |
| | ATGATGCGCTTATACGAGGTCATGTAGTTTTTGTGTCCAGCGCTATGTCTCAGTGTCAGACGATTTGG |
| | ATGTGCCACACACGTGCGGACCAGTATTGGAAATGAGTGTAGCCGGCTCGATATCGTTGAGTTCATAG |
| | AGTGGAACTTATAGTTCGCTAGAATGGCATGGACACGTTTGGGGTTTGGACCTGTTTCGAGGATTCCA |
| | TGGTTGATGGTTGGAGCGTCTTTTTGTACAGAGCAATCTTGATTATTAGTTTTGGCTAGCGTAAGATG |
| | GCGATGCATG |
| Cg10 | CAGCATGGAGAAAAAAATCACTGGAAAACGTTCTTCGGGGCGGTCTCATGCCAGTGCCGGTTTTGAAA |
| | GGGTTCATAGATTTGTGTAGTTACCTTGGAAAACAGTCGGGCGAACATGCATAATAAGTTTGCCTATA |
| | TATTGTGACTTGCCAGTACCTGGTTCGAGATGGAGATGTGTTGTTTTATATAAGGAGCATAACGTGTT |
| | ACGATAGTCTCCGTTGAAGAACACTTTAATTATGTTTGTATGCTGATGATGAGGATGTATG |

[1] The ATG sequence at the 3′-end of each PromU sequence is the start codon of the *chloramphenicol* gene.

#### 1.8 *Streptomyces albus & Streptomyces lividans*

##### 1.8.1 Growth conditions.

The *Streptomyces* strains were grown in Yeast Extract Tryptone medium (YET, 16 g/L Tryptone, 10 g/L, Yeast Extract, 5 g/L NaCl). For sporulation, *S. lividans* TK24 was grown at 30 °C on ISP4 agar (BD Difco ISP medium 4), *S. albus* J1074 at 30 °C on Soy Flour Mannitol Agar (SFMA, 20 g/L soy flour, 20 g/L mannitol, 20 g/L agar). Conjugation reactions were performed using the same media supplemented with 10 mM $MgCl_2$ at 30 °C.

##### 1.8.2 The construction of GeneEE plasmid DNA libraries in *E. coli.*

For the functional screening of the GeneEE plasmid DNA libraries in *E. coli*, *S. albus* and *S. lividans*, a shuttle expression plasmid based on pKC1218 (Table 4) was modified as follows:

The pKC1218 backbone was amplified using primers pKC-Km-F and pKC-Km-R (Table 1) introducing 40 nt homology overhangs to a 1100 bp SalI-fragment of pHH100 containing the *aph(3')* kanamycin resistance gene, and the cassette was cloned between oriT and SCP2 replicon by *in vivo* homologous recombination in *E. coli*[22] yielding pKE101.

pKE101 was linearized by PCR using primers BsaI-Am5-F and BsaI-oriV3-R (Table 1) introducing BsaI-overhangs on both ends. The product was phosphorylated and re-ligated and the circularised product digested with BsaI. GeneEE(–SD)BsaI was digested with BsaI and ligated directly upstream of the promoterless *aac(3)IV* gene, leading to GeneEE library in pKE101. This library was then transferred to turbo competent *E. coli* C2984 cells (NEB) by chemical transformation. Transformants were selected on agar plates containing LA supplemented with 50 µg/mL kanamycin. The entire population of obtained transformants (lawn on plate) was pooled and plasmid DNA isolated with QIAprep Spin Miniprep Kit (Qiagen). 1 µg of plasmid DNA was then transformed to chemically competent *E. coli* S17-1 cells for conjugal transfer of the library to the *Streptomyces* strains. *E. coli* S17-1 transformants were selected on LA supplemented with 50 µg/mL kanamycin. For the GeneEE sequences that lead to functional ArtPromU sequences both in *E. coli* and *Streptomyces* strains, transformants were also selected on LA supplemented with 50 µg/mL apramycin.

##### 1.8.3 The functional screening of GeneEE plasmid DNA libaries in *S. albus* and *S. lividans.*

For the functional screening of GeneEE plasmid DNA libraries based on apramycin resistance phenotype, the GeneEE plasmid DNA library established in *E. coli* was conjugated from *E. coli* S17-1 to *S. albus* and *S. lividans*. Conjugation reactions were performed as described previously[23] with minor modifications. S17-1 library transformants (∼20,000 transformants for both libraries) were pooled by adding 3 mL LB to a 14 cm agar plate and the cells brought into suspension using a sterile glass rod. A volume of 100 µL of the obtained suspensions was used to inoculate 25 mL LB and incubated at 37 °C for ∼2.5 h until the cells had reached an $OD_{600}$ nm of 0.4. The cells were centrifuged at 2,000 xg for 5 min at room temperature and the obtained pellet re-suspend in 2 mL fresh LB and placed on ice. Spore suspensions of two freshly sporulated plates of *S. albus* and *S. lividans* were prepared using 4 mL $sdH_2O$ and the obtained suspensions filtered through sterile cotton wool. A volume of 50 µL of these spore suspensions were added to 500 µL 2x YT medium and incubated at 50 °C for 5 min to induce germination. The spore suspensions were cooled under running water before 500 µL of *E. coli* suspension was added, and the resulting suspension was mixed by inversion before spreading it onto two 9 cm agar plates (SFMA + 10 mM $MgCl_2$ for *S. albus*, and ISP4 + 10 mM $MgCl_2$ for *S. lividans*). Conjugation plates were incubated at 30 °C for 16 hours before overlaying them with antibiotic solutions yielding final concentrations of 30 µg/mL nalidixic acid and 50, 250 and 500 µg/mL apramycin in agar media. The plates were then further incubated at 30 °C until exconjugants appeared (3 days). Single exconjugant colonies were transferred to fresh plates supplemented with corresponding concentrations of apramycin.

Single colonies of exconjugants were colony PCR-amplified with the primer pair 5511-F and 234-R (Table 1) amplifying a 710 bp fragment surrounding the ArtPromU sequence region. Single colonies were picked into 100 mL of 200 mM lithium acetate and 1% SDS and incubated at 70 °C for 5 min. 300 mL of 96% ethanol were added, the suspension vortexed and centrifuged at 15,000x g for 3 minutes. The pellet was washed with 70% ethanol, dried and dissolved in 20 mL sterile deionized water. 1 mL of the resulting solution was used as template for PCR reactions using Taq polymerase (NEB). Amplicons of the expected size were extracted from 0.8% agarose gels and purified using the QiaQuick Gel Extraction Kit (Qiagen), and were Sanger sequenced (GATC Biotech, Germany) with the sequencing primers 5654-F-Seq or Am-5c-R-Seq (Table 1). The identified ArtPromU sequences with their corresponding phenotypes (apramycin resistance levels) are listed in Table 15 and 16.

Table 15: The ArtPromU sequences identified in *S. albus*.

| **Clone** | **Am**[1] | **DNA Sequence (5′→3′)**[2] |
|---|---|---|
| Sa01 | 50 | TGTGATGGAAGATATTAGTGAGAAGTCTCTGATGCGAAGTTTTTAGTTGGCCCCGGTTGTTGCTGT<br>TATACAATATCTAACAATCCCCGACGAGCAATACACCGGGTTCCAGCACCTGGGACATGTTTTACG<br>CGTCTGAAAATATGATTAGAGAGGAGTAGTGAGTGTCTCGAGGACTGATG |
| Sa02 | 50 | GAACATATGGGAGGAAGATGGATACCCTATGTGTTCCCGGTATGTATCATGTGATATATCGGTTCA<br>ATGGCACGAGACAACGAGAGGGACAGGTCTAAAGGCCATTATATGGGCGGTAGCGTATGTTTGTGT<br>ATTCGTTAGGTCAGTTGTGGTATCAACAATTAAAGTTCAACGTCTATGCTAGTATGGGGGCCAAAT<br>AGATG |
| Sa03 | 50 | CTCTGTGTTACCAGATCTTTAAGGTATAGACAGGTGTGTTCGAGCGATGGATTTTTTTTTTCTTTT<br>AACTCATTGTAGCGCTAAAACTGTTGGTCGTATGCGGGCTATTGGTACATGTGAAGGAGTCACAAG<br>TCTGGCAAGCAGGTTGTGTCATGTGTAGGTACCAGAATAAGCAGGCATAGGGTAGACTGGAGGGGA<br>TTATG |
| Sa04 | 50 | TCCTTACACATTGGTATTTTTCTCCTAGTTGCCTGTCTTTGCAATCAGTACTCTTGACCGTTTTTT<br>CCATGGATCAAGAGTTAAAGGGTGGGGGTGAATGTACTCAAATTATGGTTCAGTGGGGGTAATATC<br>CGTTCTGACTGTGAACATTAGCACAGTAGTGTGGCGGACGGCGGCGTTGTTGGTATAGTATGGGCT<br>TGATG |
| Sa05 | 50 | TGCCCTTTCATTAAGGTAATGGAAGCGGGCATGGTATGGTACGTTTCGTTGATCAAGTTGTTTTAA<br>CAGTTTTTGAGGGTAATTTGCGTATCCCGTAGTATAATTATGCTTTGTGCGTCAAGATCATGGGGG<br>CTGAGACCTTGGTGTTCCCGGCCTACCGAGCGGCTCTTAGACAAAAATTTAGGGTGAAATGAGAGG<br>TCATG |
| Sa06 | 50 | TTCTCTTATCAATAACTTAGTAAAGACCTTGGGAGACAATCGGGAGATGCCGTGACGTGGGTTGCG<br>AGTTGGGTACGTCCTTTACGGAAGGTCATGCGTGGTAAGTTGCGGTGCGTATGGAGATAAGATAAG<br>GTGGAGCATG |
| Sa07 | 50 | AGTTTCGCATATTAATAATATAAGTGTATATGCTAGACTTTTAAGCGGGTTTTATGATAGTGTATG<br>GGAACGTGCGGCGGCAGTTTATATTGGAGTTGTAAGAGGGCATCGTAGACGCTAAGGTGGTCCAGT<br>GGATGCTAGGAATACTTGGACTATGCCAGAGGCTAGTTAAGAGCGGCACGTTTTTGGTGGGCGGAA<br>AGATG |
| Sa08 | 50 | GCTTATATGGTGGTTGTGCTGGTGCTTTCTCTTTTAGAGGCTGCGATGCTCGTAATGTCAGTTGAA<br>CGATTGACAAGGGAGAAGCCAGGTACTTATGGTACCAGGTTATGACGCCATGTGCTGTGGAGCACC<br>GGTTTCAAGCAGACGTGGCACATACTGTAGGAGCGAAGCATATGTGCAGGTGGCCAGCGGAGAGAA<br>ATATG |
| Sa09 | 50 | CTCTTGATGGGACTGTTGGAGGGCGTATACGTGGATTATTAATTTGTTGAACCGTGGCCAGTTTCC<br>GTTGATCTAGACTTTTTTGCGGTGTCGCTGTCTCATTTATTTTTATCTGTAAATCAGATTTACTCT<br>CTACTACCCGTATTAAGTCACGAGTGCCGTATAATTCTTAGTGTCTGTGTGATACAGTAACTAGGG<br>CTATG |
| Sa10 | 50 | AAGGAAAATCAAGAGTTTGAACGGCTTGAGTGAACAACATCTCGACTAGACTGAACGATTTCATGT<br>CAAACTCGTGTTCGGGATCCGCTATGTATGGATGTGTTGAGGCTTTTAATGTTATCATTAGAGCTA<br>GTGTGGTGTGTACTCAAGTTTTAAATCGCATGCCCGGGTGAGTCAATTGACTGGAGGTCGCGGATA<br>TCATG |

Table 15: continued from the previous page.

| | | |
|---|---|---|
| Sa11 | 50 | GTTCCAGTTTCGGAGCAGTGCCATTGGTCGAATGAGTTCTCTGGTGAACGAGGCGTTAATAATTTT<br>CGGTGTCAAGTAGTCGATATTCCTGGTTTTTAATTATATGATCTTGTAATTATACATGGCCTTCCT<br>TCTTGAAATGCCAGAGTAGTATTCGACTTTATGGATTGAGACGAGGACGAATGTTGAGTCGGTGGG<br>GGATGCTGCAGCGGCCGCTACTAGTAGGTCTCGCATATATATGAGATACGGTACACTGGATAACAT<br>CCAGATTCTGAGGGTGTGAGCAGAGTGGAGCAGAATTGTTCCGACGCGAGATCGTCTTGGTGCTAG<br>ATAATGCTTAAGGATTGAATTATTCCTGGCGTAAATGTCAATATTGGAGCGTATTATGATG |
| Sa12 | 50 | AGAATTTTTGTTGCTTGTAAATTATATGTTCAAAAGATTTGAGTGTATGATCAGTGTGTGGTATGG<br>TGCGTTTGTCCTTTTAGCCCCTGTCTTTTTTCTCGGGCGATGTGTGTTGCACGTGTAAATGATTCC<br>TGGCTCCCATTAATAAGCTGGGTATTAGACTTCGGCGTGAAGTTAATGGAAGACGGGGTAGACGAC<br>GGATG |
| Sa13 | 50 | AAGCTAACTGGTGCCTGAGTGGATCTTCGAATTATGTTTCTCAGATCTTTTTTTTTGTCTTGCTTG<br>TATTTGTTGCGTCTATTAGATTCTAGATGTATAATAATATGCTCGGCCTGTTTTTCCATAATAAGC<br>TAGTCACAGTATCTTGGCGACTTCATTGTGGAGCATTTGGGTGGCCCTAGTTCAGTTTGGTGTGCG<br>CAATG |
| Sa14 | 50 | GTAGTTTTGTCTATTGGTTAAATTCTTCGGTAAGCGGTCATCGACGATACATCCATGCTAGGTGCA<br>TTTGTGCGTGGGCAACCATTTTCGTGTAGGTAATTGATACCATGGTATATGTAATGTGACACAGGC<br>ACACGTCCGCGTATACATCGTATTCAATGATTCCGGAATGACGTGGGCTATACTATCATAGTCGTT<br>AGATG |
| Sa15 | 500 | TTCTCTTATCAATAACTTAGTAAAGACCTTGGGAGACAATCGGGAGATGCCGTGACGTGGGTTGCG<br>AGTTGGGTACGTCCTTTACGGAAGGTCATGCGTGGTAAGTTGCGGTGCGTATGGAGATAAGATAAG<br>GTGGAGCATG |
| Sa16 | 500 | CTACCATGGCAGGGCTTTTACGTGGATATCTTCTTTGAGTGTTTGCCTATCGGTTTGTGATATGCT<br>TAGATCTATCAGGTGTCGACTTCCGTACTTGTTGGTAATGCTCAGATCATCAATCGAAGGGCGAAT<br>GTTGTACAATCAGTATGTGACGGTCTGAGGGGGACGGTTTTGTATGGGGCGCCGGGGAGTTGGTTA<br>GTATG |

[1] Apramycin concentrations used for the selection (µg/mL).

[2] The ATG sequence at the 3′-end of each ArtPromU sequence is the start codon of the *aac(3)IV* gene.

Table 16: The ArtPromU sequences identified in *S. lividans*.

| **Clone** | **Am**[1] | **DNA Sequence (5′→3′)**[2] |
|---|---|---|
| Sl01 | 250 | AGGGACCCTAAGACGCTTTGGCTTTCTGGAGTTTCAGGTACAGGTTTCATATCTGTATTGTTATA<br>CTTTGTCGACGAGAGTTGTAGGACTGATAAATTGCGTTAGATTCTACAGTCAACGTGCGTTGGGC<br>CTATGGATTCGGCGTCTTTGTCCAAAATTTGTCGAGTTCCTTCGGGCGATGTTTGGAGGTATCTG<br>TAAAGATG |
| Sl02 | 500 | CTACCATGGCAGGGCTTTTACGTGGATATCTTCTTTGAGTGTTTGCCTATCGGTTTGTGATATGC<br>TTAGATCTATCAGGTGTCGACTTCCGTACTTGTTGGTAATGCTCAGATCATCAATCGAAGGGCGA<br>ATGTTGTACAATCAGTATGTGACGGTCTGAGGGGGACGGTTTTGTATGGGGCGCCGGGGAGTTGG<br>TTAGTATG |
| Sl03 | 500 | TCGGTCGCGGAGTGAATATCCCCTGTTGCGAGGTATAAGTGTATTCTTGCGTATTGGGAGATTGT<br>GGTCGCAGATCGTATGCTTGATCTGGTGTCAATTTAGGGAAGTAATTACCTGACGAGAGGTCGTT<br>GTCTCGGTTTGAGTATAAGGCTATATGTGTGTTTTGCGTGTTGGGGCCAAGATGCGCATAAGATG<br>ATGGTATG |

Table 16: continued from the previous page.

| | | |
|---|---|---|
| Sl04 | 500 | GAAATATCTCTCTGTCTTGAACGGGGTTGTAAGGGATATAATTCCTAGGATCGTTGACTCTCCCG GACGGTTCTTTCGGGCGGTTTTCTCGTTGTTCCAGGTTGGGAGTGTTCTGGCTTTTGAACCTCTT GCATGCTGTCAGGGGTGGTTGTGTATAATCGCTGAGTTGTTCATACAAATTGGTTGGGAGTGTTC GCAAGATG |
| Sl05 | 500 | TTCTCTTATCAATAACTTAGTAAAGACCTTGGGAGACAATCGGGAGATGCCGTGACGTGGGTTGC GAGTTGGGTACGTCCTTTACGGAAGGTCATGCGTGGTAAGTTGCGGTGCGTATGGAGATAAGATA AGGTGGAGCATG |
| Sl06 | 500 | CTAGGTGCAGCACGCATTGGAGACGAGTAGTGTACTTGAAGGGGGGTGATATGATGCGATCCGTT TATTGTCCTATGGCGCGAATCTAACGTGCGTGTGGTTGCAGGAATAACGTATACCTTCTTTTTGG ATCCTTGGGGGTATCGAATATATACGCGCCTCGAATAGGGAGCTTAGATG |
| Sl07 | 500 | TGTTGTCCGAGTCTTTATCTCACGTTGTGATGGGTCCCTGTTTTCTTCGGTGGAGGCTTTGAAGG AATTCGTCCACAAGAGTTAATAAGCGCCCTCGCACCGTTTGAATAGCGTTTCGTTGTATGTTGTA TGTGTTAGGAACGTCATTGTTTCCATATTCAGGGTAGGGGGGGTCACTATGGTCTACTTAGCGCA TGGGGATG |
| Sl08 | 500 | TTCTCTTATCAATAACTTAGTAAAGACCTTGGGAGACAATCGGGAGATGCCGTGACGTGGGTTGC GAGTTGGGTACGTCCTTTACGGAAGGTCATGCGTGGTAAGTTGCGGTGCGTATGGAGATAAGATA AGGTGGAGCATG |

[1] Apramycin concentrations used for the selection (µg/mL).

[2] The ATG sequence at the 3′-end of each ArtPromU sequence is the start codon of the *aac(3)IV* gene.

#### 1.9 *Saccharomyces cerevisiae*

##### 1.9.1 Growth conditions.

*S. cerevisiae* cells were grown in Yeast Extract Peptone Dextrose Broth (YEPDB, 20 g/L bacto peptone, 10 g/L yeast extract, 20 g/L dextrose) or YEPD agar (YEPDA, 50 g/L YEPDB, 15 g/L agar) (Sigma-Aldrich) at 30 °C. For the functional screening of GeneEE segments controlling the expression of *tryptophan* gene the Drop-out media (0.68 g/L yeast nitrogen base powder [Sigma-Aldrich], 0.5 g/L glucose, 1.92 g/L of Yeast Synthetic Drop-Out Media Supplements without tryptophan [Sigma-Aldrich]) was used.

##### 1.9.2 The construction of GeneEE plasmid DNA libraries in *E. coli*.

For the construction of GeneEE plasmid DNA libraries in *E. coli*, an *E. coli*-*S. cerevisiae* shuttle vector, pENZ004, was used (Table 4). The pENZ004 plasmid provides the homologous regions suitable for genomic integration, via gene replacement into chromosome II; and the *tryptophan* gene used for the functional screening in *S. cerevisiae*. pENZ004 was PCR amplified using the primer pair Trp-F and Trp-R (Table 1) leading to the amplification of the entire plasmid excluding the native *tryptophan* promoter. The reverse primer Trp-R carries the BioBrick Prefix[1] as a flanking sequence that serves as a homologous sequence in Gibson cloning. This amplicon was used as the backbone and the GeneEE(–SD)Trp as the insert. The Gibson ligation mixture was made using equimolar amounts of purified vector and insert (totalling 100 ng DNA) in a total volume of 5 µL, added to a volume of 15 µL of Gibson assembly mix. The mixture was then transferred into the cloning host *E. coli* via chemical transformation and grown on LA plates supplemented with 50 µg/mL ampicillin. The resulting library was consisting of ∼10,000 transformants which were scraped and the plasmids were isolated with QIAprep Spin Miniprep Kit (Qiagen) and later quantified with Nanodrop (ThermoFisher).

##### 1.9.3 The functional screening of GeneEE plasmid DNA libraries in *S. cerevisiae*.

The GeneEE plasmid DNA library isolated from the scraped *E. coli* cells was linearised with SfiI in CutSmart Buffer at 50 °C. *S. cerevisiae* cells were grown in 5 mL of 2x YEPDB on a rotary shaker at 200 rpm at 30 °C. To determine the titre 100 µL of the *S. cerevisiae* culture was inoculated into 900 µL of 2x YEPDB in a spectrophotometer cuvet and measured at $OD_{600}$ nm (an $OD_{600}$ of 1 equals to $3x10^7$ cells/mL). Cells were inoculated to 25 mL of pre-warmed 2x YEPD to a titre of $5x10^6$ cells/mL, and were grown for 4 hours at 30 °C, with agitation until titre reaches $2x10^7$ cells/mL. The cells were harvested by centrifugation at 3,000 xg for 5 min and the pellet was washed twice in 25 mL of sterile deionised water, and were resuspend in 1 mL of 0.1 M LiOAc. The DNA was transformed to *S. cerevisiae* using the established LiOAc protocol[24], and the resulting transformation mixture was plated on Drop-out media and the plates were incubated at 30 °C for 4 days. To sequence the ArtPromU sequence, crude yeast genomic DNA extractions were performed as described previously[25]. The genomic DNA was then PCR amplified using the primer pair Hom-F and Hom-R (Table 1). The resulting amplicon was PCR purified with QiaQuick PCR Purification Kit (Qiagen) and was Sanger sequenced (GATC Biotech, Germany) with the primer pENZ004-Trp-Seq (Table 1). The identified ArtPromU sequences are listed in Table 17.

Table 17: The ArtPromU sequences identified in *S. cerevisiae*.

| **Clone** | **DNA Sequence (5′→3′)[1]** |
|---|---|
| Sc01 | AAGAGTCGCGCTCAGTTTTCGGTTGCTTCCTGGTTATCTAGAGGGAATTTCGATGATGCATTTGCCTA<br>AATACTAGCTAGTGTCAAGAGATGATATTATGTCTTATTTAGACTTAGCGCGGAACTAAGAATTGTCG<br>TTTATTTACTGTCTATGAACAATTTCGTGGACTGTGTAATAGTTGTAACTCGGTAGATG |
| Sc02 | GTCTGTGTCATGCTACAAGGGTGACGGTGCAAGTATTTTGAGATCACAGTCGTACATGCGTCCGCATT<br>TCTTGAATATTTGCACTATCACAACTATATTACTGTTCTGACAAAAGTCATCTTATTTTGATTAATGT<br>TGGGGGGGTGTAGTAATGGACGCGCTAGTAATTTGTAGGTTTGTGTCAGAAAGCTCATCCGAGATG |
| Sc03 | TATTGGTAATAGTACCGGTTGTAAAGTTGCCGCTGTCGAACGTGGTTATTTATTTGGTGTTACAATCT<br>GGGCCGCGGGTTACTCTTTTATCTCAAGTATTTGTGTGGTTGTCGTTTCGCAAGCTATGTAACTCAGT<br>CTAGACGATTGGACTAGAGGTAGGATATTTGTCGGAGATTACTATTGGGAGTTTCTTGTAGAGTATG |
| Sc04 | AGGCTTGCCGCAGAGTGACTGATCGGTATATTCGTTATTATCTAATTTTCTTGTTTGGCGGATGGAAT<br>ACGATAGCGTTGATTGATGTTCGTTTTTCTTGTGCAGTGGCGGTTTGGGTGTTAGGGGTAAGTGATCG<br>TGAAAGAATCAACGGTTAGATCGAAATGAAGTGAGATTCAGTGTTGGGGATGGAAATGCTAAGGATG |
| Sc05 | CAACTGGCGCGGCAATGTTGTTACTTTTAAGAGTTTTCTTCTCGATCCTTACCAAAGGAATGGATTCT<br>TATTGAGATTTAAGCTGGCTTATGTTGTAAGTCTGTTACAATAAGTCGTGCGAAGGCGGTTTACGAAG<br>AAGGTAGTCTGGGAGTCCTGTACTTGTTTATCATCTATTATTTGTTTTTCCAAGACCGGGGTTGATG |
| Sc06 | GCCCGGTTACCGGGGGTATTGGCGAAAAATTAAGTGGTGAGCTGCGAGGTCTGGGTTACAGTGGTGAC<br>CTTTTATAGTATCATACTTTAGTTAGGTTGGATATCGATATCCATTTAAGGGAATTATCCATTCCACT<br>CTGGTTCTTCCTATGTGAAGAAGATATTGGATTATTAGTTTAGCGGGAGCGTTATCTTATTTATATG |
| Sc07 | GTGATTGTTGCTAAGCCGCAGAGGGGTTTCAGTGGGAAACGATAATCAGATACATGGGGGTTATGAAA<br>CCTTCTAGCTGGTAGTTTGCTTTTGCGATTTAAAGGTTGACGATGCGGTCGACTTCTCATCTGTTGAT<br>GAGGGTTTAGGGTGTGGCATCCTTTACTACGGCTTGAGCAAAGAGTACTGTAGTCTCGTGGTATATG |
| Sc08 | GTACATTATTTCGTCATCTGCAACGAAATTAACTTTTTGCGAATATTTAGGTTGATGTATTTGCCATG<br>GGAAAATGAGTAAAATTTAGCGGTCTGGGCATGCCTCCACAGGGTGCATAGTATGGTGAGGCTGTGGA<br>ATGCACGTTGTACGATTCAGTATATTAAGATTATTAATTGACTCGGTAGGGAAGTGGCGTGGGTATG |
| Sc09 | TCGCGTACTTTATGCTCACGTTATGCGACCACGTAAAAGGCGCGCGGTATGATATGCAGGGCTTACAT<br>TCTTTTTAAAAATCCGACGCTCGGGTTTTGGTGATGGGTTCTTTCGTTGACCATTGTTGCTGATGGGC<br>TTTTCGTTAAAATAATGGTAAGGATTCAGTTTAGGGGTTTTACTGTTTGAAGATGGCTTTATGTATG |

Table 17: continued from the previous page.

| | |
|---|---|
| Sc10 | CGTCTTAGGCTGGAAAATTCAGGTGGCGAGGATGGCTTTAGTTTTGAAATGAAACATTCGATTTTCTT |
| | GCTGCGGAGATCCCTTTTCTTCCGTTTTTTTAAAAATGCAGAGGATGTTCAGAATGCGGTCAGCTACA |
| | ATTGTGTTGGTGTGGGATGGAATTGGTTGTTAGGGGAGTGATATCAGGTGAAAAGTAGGAGTGGATG |

[1] The ATG sequence at the 3′-end of each ArtPromU sequence is the start codon of the *tryptophan* gene.

[3] *Pseudomonas putida* KT2440 chromosome, complete genome
URL http://www.ncbi.nlm.nih.gov/nuccore/NC_002947.4. 1002825811.

[4] *Thermus thermophilus* HB27, complete genome
URL http://www.ncbi.nlm.nih.gov/nuccore/AE017221.1. {:itemType: dataset}, 46197919.

[5] Corynebacterium glutamicum ATCC13032, complete genome
URL http://www.ncbi.nlm.nih.gov/nuccore/BA000036.3. 42602314.

[6] Chater, K. F. & Wilde, L. C. *Streptomyces albus* G mutants defective in the SalGI restriction-modification system. *Microbiology* **116**, 323–334 (1980).

[7] Hopwood, D. A., Kieser, T., Wright, H. M. & Bibb, M. J. Plasmids, recombination and chromosome mapping in *Streptomyces lividans* 66. *Microbiology* **129**, 2257–2269 (1983).

[8] CEN.PK2-1c-euroscarf. URL http://www.euroscarf.de/plasmid_details.php?accno=30000A.

[20] Improbizer. URL
https://users.soe.ucsc.edu/ kent/improbizer/improbizer.html.
