## Supplementary material for "A universal method for gene expression engineering": perl script: motifs

use strict;

# simple script to parse multiple files with FLASh joined read data
# to identify the most likely sequence belonging to a certain position 

# hash to store the parsed data
my %data;

# helper hashes to identify the number of rows and columns in the setup
my %rows;
my %cols;

foreach my $file (@ARGV) {
    # get the ID 
    my ($id) = ($file =~ m/(\w+)_comb.fasta/);

    # if the ID is a char, the data comes form a row, otherwise it is a column
    if ($id =~ /[A-Z]/) {
	$rows{$id} = 1;
    } else {
	$cols{$id} = 1;
    }
    
    open FILE, $file;

    # go through the reads, grep out the N block
    # requires perfect matches to the flanking sequences
    while (<FILE>) {
	my ($seq) = m/GCCGCTTCTAGAG(\w+GGAG\w{7})ATGGTTTCTAAAG/;
	if ($seq) {
	    $data{$id}{$seq}++;
	}
    }
}

# hash to store the best hits found
my %bhits;

# go through the data, first on row
foreach my $row (sort(keys(%rows))) {
    # check each sequence found in a row
    foreach my $seq (sort(keys(%{$data{$row}}))) {

	# discard spurious matches (CAVEAT: this might cause problems in low abundance pools)
	my $rowcount = $data{$row}{$seq};
	next if ($rowcount < 5);
	
	# go through the data, second on row
	foreach my $col (sort by_number(keys(%cols))) {

	    # discard spurious matches (CAVEAT: this might cause problems in low abundance pools)
	    my $colcount = $data{$col}{$seq};
	    next if ($colcount < 5);
	    
	    # if a sequence is found more often in both, row and col, it replaces the previous best hit
	    # CAVEAT: imbalanced pools could cause trouble here
	    if (($rowcount > $bhits{$row.$col}{rc}) && ($colcount > $bhits{$row.$col}{cc})) {
		$bhits{$row.$col}{rc}  = $rowcount;
		$bhits{$row.$col}{cc}  = $colcount;
		$bhits{$row.$col}{seq} = $seq;
	    }
	}
    }
}

# the sequence downstream of the N block, needed for mapping of TSS data
my $end = 'ATGGTTTCTAAAGGTGAAGAAGACAACATGGCTATCATCAAAGAATTTATGCGTTTCAAAGTTCACATGGAAGGTTCTGTGAACGGTCA';

# print the best hits: FASTA to STDOUT, table to STDERR
foreach my $id (sort(keys(%bhits))) {
    print ">$id rowcount=$bhits{$id}{rc} colcount=$bhits{$id}{cc}\n$bhits{$id}{seq}\n$end\n";
    print STDERR "$id\t$bhits{$id}{rc}\t$bhits{$id}{cc}\t$bhits{$id}{seq}\n";
}	


# numerical sorting
sub by_number {
    $a <=> $b
}
