## Supplementary material for "A universal method for gene expression engineering": perl script: starts

use strict;
# script to parse TSS data and output FASTA and table for further analysis

# path to Statistics::Descriptive;
use lib "/vol/biotools/share/scripts/";
use Statistics::Descriptive;

# sequence upstream of the N block (for promoters at the border of the cloned region 
my $static = "tttccttggtgtatccaacggcgtcagcgaattcgcggccgcttctagag";

# hash to store the plate data (position, FIM and sequence)
my %seqs;

# first file has to be a TSV containing position, FIM, and sequence
open FIMS, "$ARGV[0]";
while (<FIMS>) {
    chomp;
    my @fields = split(/\t/);
    $seqs{$fields[0]}{fim} = $fields[1];
    $seqs{$fields[0]}{seq} = $fields[2];
}

# hash to hold the mapping data
my %data;

# total number of mappable positions
my $total_len;

# initialize the arrays in which the mapped start positions will be stored
foreach my $id (keys(%seqs)) {
    my $len = length($seqs{$id}{seq});
    @{$data{$id}} = (0) x $len;
    $total_len += $len;
}

# read in the SAM file
open DATA, "$ARGV[1]";
# counter of total hits observed
my $total_hits;
while (<DATA>) {
    # skip headers
    next if (m/^\@/);
    chomp;

    # split the SAM string
    # [1] holds the binary flag
    # [2] holds the sequence id
    # [3] holds the offset (position of the TSS)
    # [5] holds the mapping info
    my @fields = split(/\t/);

    # consider only mappings on the forward strand
    next unless ($fields[1] == 0);

    # consider only mappings where the beginning is mapped 
    if ($fields[5] =~ m/^\d+M/) {
	my $id = $fields[2];
	
	# remove spurious hits downstream of the N block
	next if ($fields[3] >= length($seqs{$id}{seq}));
	next unless ($seqs{$id}{seq});
	$data{$id}[$fields[3]]++;
	$total_hits++;
    }
}


# do the simulation: scatter the total number of reads randomly over the total number of mappable positions
my @sim = (0) x $total_len;
for (my $i; $i < $total_hits; $i++) {
    $sim[int(rand($total_len))]++;
}

# do the statistics
my @simcounts;
foreach my $count (@sim) {
    push (@simcounts, $count);
}
my $stat = Statistics::Descriptive::Full->new();
$stat->add_data(@simcounts); 
my $mean = sprintf('%0.0f', $stat->mean());

my $threshold = sprintf('%0.0f', (3 * $stat->standard_deviation() + $stat->mean()));

# hash to store data for FASTA output
my %om;

# now parse process the data for each sequence
foreach my $id (sort(keys(%seqs))) {

    # the seqquence, add the ATG of mCherry
    my $seq = lc($static . $seqs{$id}{seq}) . 'ATG' ;

    # the FIM data
    my $fim = $seqs{$id}{fim};

    # shift flag: was the TSS shifted to the position upstream?
    my $flag = "No";
    # flag for multiple TSS in a given sequence
    my $seen;
    # the string with all mapped starts for a given sequence, CSV format
    my $hits = join(',', @{$data{$id}});

    # string that holds the output for the table
    my $tssfields;
    
    # total number of mappings for valid TSS for a given sequence
    my $tsstotal = 0;

    # go through the positions, from 3' to 5'
    for (my $pos = $#{$data{$id}};  $pos > 0; $pos--) {
	# if the number of mappings starting at the given position is higher than the random background...
	if ($data{$id}[$pos] > $threshold) {
	    # and the increase is steep considering the position directly upstream, consider $pos to be a valid TSS 
	    if ($data{$id}[$pos] >= 3 * $data{$id}[$pos-1]) {
		
		$tsstotal += $data{$id}[$pos] ;
		$tssfields .= "\t\t\t\t" if ($seen);
		my $down = uc(substr($seq, $pos + 50, 1)) . substr($seq, $pos + 51, 25);
		my $up = substr($seq, ($pos + 1) , 49) . uc(substr($seq, $pos + 50, 1));
		$tssfields .= "\t$pos\t$data{$id}[$pos]\t$flag\t$up\t$down\n";

		push (@{$om{$data{$id}[$pos]}}, (">$id\_$pos\_$data{$id}[$pos]\_$flag\n$up" )) unless (length($up) < 50);

		# reset the shift flag
		$flag = "No";
		# set the multi TSS flag
		$seen = 1;
		$pos--;
	    } else {
		# in case of "fuzzy" TSS, shift the TSS one position upstream 
		$data{$id}[$pos-1] += $data{$id}[$pos];
		$flag = "Yes";
	    }
	}
    }
    
    print "$id\t$seq\t$hits\t$tsstotal\t$fim$tssfields";
    print "\n" unless ($tssfields);
}

foreach my $val (sort by_number(keys(%om))) {
    foreach my $line (@{$om{$val}}) {
	print STDERR"$line\n";
    }
}


sub by_number {
    $b <=> $a
}
